# PhorEau: a new process-based model to predict forest functioning, from tree ecophysiology to forest dynamics and biogeography

**DOI:** 10.1101/2025.01.28.635328

**Authors:** Tanguy Postic, François de Coligny, Isabelle Chuine, Nicolas Martin StPaul, Xavier Morin

## Abstract

Climate change impacts forest functioning and dynamics, yet significant uncertainties persist regarding the interactions between species composition, demographic processes, and environmental drivers. While the effects of climate change on individual plant ecophysiology are better understood, few robust tools integrate these processes dynamically, hindering accurate projections and recommendations for long-term sustainable forest management. Forest gap models strike a balance between complexity and generality and are widely used in predictive forest ecology. However, their lack of explicit representation of critical processes—such as plant phenology and water use—limits their ability to fully capture tree sensitivity to climate change, calling into question the robustness of their future predictions. Therefore, incorporating trait- and process-based representations of climate-sensitive processes within gap models is a crucial step toward generating realistic predictions of forest evolution under climate change.

In this study, we coupled the ForCEEPS gap model, validated across a broad range of forest types and environmental conditions in Europe, with two process-based models: a plant phenology model (PHENOFIT) and a plant hydraulics model (SurEAU), each parameterized for the main European tree species. We then evaluated the performance of the resulting PHOREAU model across multiple processes, metrics, and time- and spatial-scales, thereby minimizing the risk of equifinality.

PHOREAU demonstrated robust capabilities in predicting fine hydraulic processes at both the forest and stand scales for various species and forest types. This, combined with its enhanced ability to predict stand leaf areas from inventories, led to modest improvements in annual growth predictions compared to the original ForCEEPS model and a strong capacity to predict potential community compositions.

By integrating recent advancements in plant hydraulics, phenology, and competition for light and water into a dynamic, individual-based framework, the PHOREAU model bridges the gap between trait diversity and long-term forest productivity and resilience. It offers insights into complex emergent properties and trade-offs linked to diversity effects under extreme climatic events, with significant implications for sustainable forest management strategies.

## 1 Introduction

Forests cover approximately 30% of the Earth’s land surface, hosting the majority of terrestrial biodiversity. They are crucial carbon sinks (Pan *et al*., 2011) play a vital role in climate regulation (Chapin III *et al*., 2008), and provide essential ecosystem services to humans (Nadrowski, Wirth and Scherer-Lorenzen, 2010). However, climate change poses significant risks to forests, including disruptions to forest dynamics (McDowell *et al*., 2020), as increasingly extreme environmental conditions have profound effects on forest structure and composition as well as on forest functioning (Allen *et al*., 2010). Despite this, there remains a lack of robust tools to effectively investigate the combined impacts of biodiversity and climate change on forest dynamics and functioning. In particular there is a key challenge inherent to using forest models, calibrated on historical data, to predict the evolution of European forests in the uncertain transition period that is facing Europe over the coming decades (Parmesan, Morecroft and Trisurat, 2022). As Europe trends towards generally drier conditions and as the limiting resource over which individual trees compete may shift from light to soil water (McDowell *et al*., 2020), the robustness of predictions will depend in large part on whether models are able to mechanistically account for the impact of water stress in relation to stand composition (Brodribb *et al*., 2020; McDowell *et al*., 2022). Instead of postulating general *a priori* species complementarity effects in resource usage based on past patterns, process-based modelling must strive to capture how individual trees harness and compete for light and water *in natura:* it is only then that local complementarities between certain species, in certain climate conditions, and for certain stand characteristics might be predicted, to be then possibly applied to forest management policies.

Depicting and understanding the role of diversity in ecosystem functioning has been a key focus of ecological studies for at least two decades (Kinzig, Pacala and Tilman, 2002; Hooper *et al*., 2012; van der Plas, 2019). In forest ecosystems, the importance of the role of diversity — both structural and compositional — on productivity and wood biomass has been firmly established by numerous studies over a wide range of conditions and methods (Nadrowski, Wirth and Scherer-Lorenzen, 2010; Morin, 2011; Paquette and Messier, 2011; Liang *et al*., 2016; Ratcliffe *et al*., 2017). In addition, there is some evidence that tree diversity could modulate the resistance and recovery of forest productivity under stress (Ammer, 2019; Jourdan, Lebourgeois and Morin, 2019; Schnabel *et al*., 2021; Blondeel *et al*., 2024), although there is no strong consensus on this point, especially regarding possible underlying processes (Decarsin *et al*., 2024). Yet despite these patterns, there remains a scarcity of data regarding the actual differences in functioning of monospecific and mixed forests, and their relative response to changing climate conditions. For these reasons, and because experimenting composition effects in mature forests is especially difficult, the evaluation of diversity effects in forest ecosystems has come to rely increasingly on process-based models (Bohn and Huth, 2017; Maréchaux and Chave, 2017; Jonard *et al*., 2020; Morin *et al*., 2021). The prospective power of these models make them key tools in testing various hypotheses on the diversity-functioning link (Maréchaux *et al*., 2021), but also in evaluating forest management practices that incorporate species mixing (Jourdan *et al*., 2021) and more generally in simulating forest-response to the long-term impacts of climate change.

While the diversity-productivity relationship is well evidenced — a global meta-analysis has shown mixed-species stands were on average 25% more productive than their respective species’ monocultures (Zhang, Chen and Reich, 2012) —, data regarding the link between species diversity and the ability to withstand extreme climatic events is more scarce and contradictory. Where some studies have linked forest diversity to a lessened sensitivity of tree growth to drought (Lebourgeois *et al*., 2013; Anderegg *et al*., 2018; Serrano-León *et al*., 2024) others have found this relationship to be strongly context-dependent (Grossiord *et al*., 2014; Forrester *et al*., 2016; Jactel *et al*., 2017), and restricted to dry environments. Moreover, with the rapid shift in climatic conditions, it would be a mistake to assume that the same patterns of diversity-productivity and diversity-resilience relationships used to support the stress-gradient hypothesis (Bertness and Callaway, 1994) will apply in the next decades to newly drought-prone sites, where water resource limitation has not had the chance to shape the co-evolution of the local species over the past millennia. In fact, the same structural and specific complementarities that are currently responsible for increasing the productivity of existing mixed temperate forests through a better usage of the light resource could become a source of vulnerability, as competition for water intensifies proportionally to the density and foliage areas of the stands (Jucker *et al*., 2014; Haberstroh and Werner, 2022; Decarsin *et al*., 2024).

Although there is a lack of knowledge regarding the effects of species mixing on forest resistance and resilience to drought, trait-data describing the hydric functioning of tree species has been steadily accumulating. Thereby a great variety of water-stress adaptation and drought response strategies among species have been identified (Brodribb, 2009; Martin-StPaul, Delzon and Cochard, 2017): these include traits linked to the allocation between transpiring and conducting surfaces, stomatal control and conductance (Johnson et al., 2012), water storage, root-to-shoot ratio, specific leaf area, safety margins (Martin-StPaul, Delzon and Cochard, 2017), and rooting depths (del Castillo *et al*., 2016). It is these traits and their variability which ultimately account for many of the plant-to-plant interactions responsible for water-competition reduction and facilitation (De Cáceres *et al*., 2021; Moreno-de-Las-Heras *et al*., 2023; Mas *et al*., 2024); however, understanding their net impact in existing forests is complicated by environmental and structural variability among stands, and more generally by the fact that the most common available indicators — growth and mortality — integrate over time many processes that are difficult to unravel.

It is this dynamic and integrative effect of species-mixing on medium-term drought-resilience which both most directly concerns forest management strategies elaborated today, and for which it is most difficult to formulate *a priori* recommendations. Decoupling the effects of hydraulic trait diversity from forest structure (foliage area, tree density) involves significant methodological difficulties (Forrester and Pretzsch, 2015), and is further complicated by the feedbacks between traits and stand structure (Guillemot and Martin-StPaul, 2024), as trees have been shown to adapt hydraulic traits such as leaf thickness or root-to-shoot ratio to forest structure (Limousin et al., 2012; Martin-StPaul et al., 2013). Furthermore, even disregarding species diversity, the relationship between forest structure, density and productivity is itself poorly understood: there is no consensus on the link between tree-size heterogeneity and productivity (Pretzsch and Biber, 2010; Bourdier *et al*., 2016; Dănescu, Albrecht and Bauhus, 2016), and while stand density has been statically correlated with increased growth (Reineke, 1933; Forrester, 2014), it is the overall dynamic interactions between these factors that must be understood (Morin *et al*., 2025). The prohibitive cost of testing all the factors affecting forest functioning (species diversity, stand structure and density, response to climate and soil conditions, effect of management…) in experimental or observational studies justifies the use of forest ecosystem models (Pretzsch, Rötzer and Forrester, 2017), which are able to replicate *in silico* the complex plant-to-plant interactions that regulate competition for above- and belowground resources, evaluate potential facilitation and competition reduction processes, and integrate them over time in stand structure dynamics that account for trade-offs between drought-resistance and productivity.

Furthermore, while many studies highlight the role of species diversity in forest functioning, it is important not to lose sight of the fact that the presence of a species in a given forest is itself the result of a historical process conditioned both by site conditions and species coexistence mechanisms. Phenological processes (including seed production, leaf dormancy and resistance to frost) have been shown be major factors in determining species distribution (Chuine, 2010); but the site conditions underlying them are themselves shaped directly by forest structure and functioning. Because of these complex retroactions, and in a dynamic context where tree phenology itself shifts with changing climates (Cleland *et al*., 2007), species diversity must be integrated not as an input but as an emerging factor of the modelling framework.

Integrating trait-based phenology is an important next-step in improving the ability of gap models to represent competition for light. Recent gap models (Maréchaux and Chave, 2017; Morin *et al*., 2021) by explicitly modelling crown sizes and species shade tolerances, have focused on capturing the processes through which canopy packing and spatial niche partitioning can emerge. However, space is not only the dimension through which plant species partition resources – time is also an important vector of asymmetry through which different species can coexist in by exploiting different niches (Gotelli and Graves, 1996). Relative shifts of even a few days in leaf phenology – either through earlier budding or later senescence – have been shown to have major impacts on plant growth, by allowing otherwise shaded understory plants to receive full sunlight (Jolly, Nemani and Running, 2004). As warming climate conditions advances the phenology of certain species, increasing productivity (Park *et al*., 2016) at the expense of additional vulnerability to spring frosts (Lopez *et al*., 2008), accurately integrating phenological responses of individual species is necessary to make robust future predictions over a wide range of forest types.

Compared to other forest ecosystem models that have been used to evaluate diversity effects for many species combinations, stand structures, and environmental conditions (Forrester and Tang, 2016; Morin et al., 2021), the PHOREAU model places a large emphasis on an integrative approach that considers the interactive and dynamic effects of several processes. Including a detailed representation of plant water use and competition for the water resource, it extends the scope of classic gap models relying on competition for light modulated by climate and soil conditions. The PHOREAU framework thus presents a coupling between recent advances in the process-based modelling of plant water relations under conditions of extreme drought (Cochard *et al*., 2021a; Ruffault *et al*., 2022) with state-of-the art phenology (Chuine and Beaubien, 2001) and light competition (Morin *et al*., 2021) in an individual-based gap-model able to consider all types of forest structures (Morin *et al*., 2025) and forest management (Jourdan *et al*., 2021). The validity of this approach is underpinned by accurate measurement of the underlying hydraulic traits for a wide range of species (Kattge *et al*., 2020),, and the explicit representation of the mechanisms regulating plant water uptake, evapotranspiration, and cavitation (Cochard *et al*., 2021b).

Even though many of the intermediary and background processes have already been independently validated, their novel coupling into the integrative PHOREAU model warrants a new validation, which we present here. This multi-stage validation relies on predictions ranging from daily hydraulic processes, yearly productivity, pluri-annual mortality, and long-term structural- and species compositions. Once validated, this model can be used to shed light on some of the many pending interrogations regarding diversity and forest functioning, such as the impact of extreme droughts on the outcome of diversity effects (Piedallu *et al*., 2023) or the role of complementarity in leaf phenology on growth in mixed stands (Morin, 2011). More generally, this new integrative tool is a unique asset for tackling issues ranging from the physiology of individuals to the functional ecology of forests and the biogeography of species.

## 2 Presentation of the PHOREAU model

Resulting from the coupling of three process-based models, an exhaustive presentation of PHOREAU would have involved rewriting the descriptions of these models, already available in their respective publications. To avoid this pitfall and for the sake of clarity we have chosen to summarize only the main processes of each model, and instead to focus on the integration methodology and the new processes allowed by the coupling. Refer to Figure 1 for a breakdown of the coupling between the ForCEEPS, PHENOFIT and SurEau models which constitute PHOREAU.

**Figure 1.**
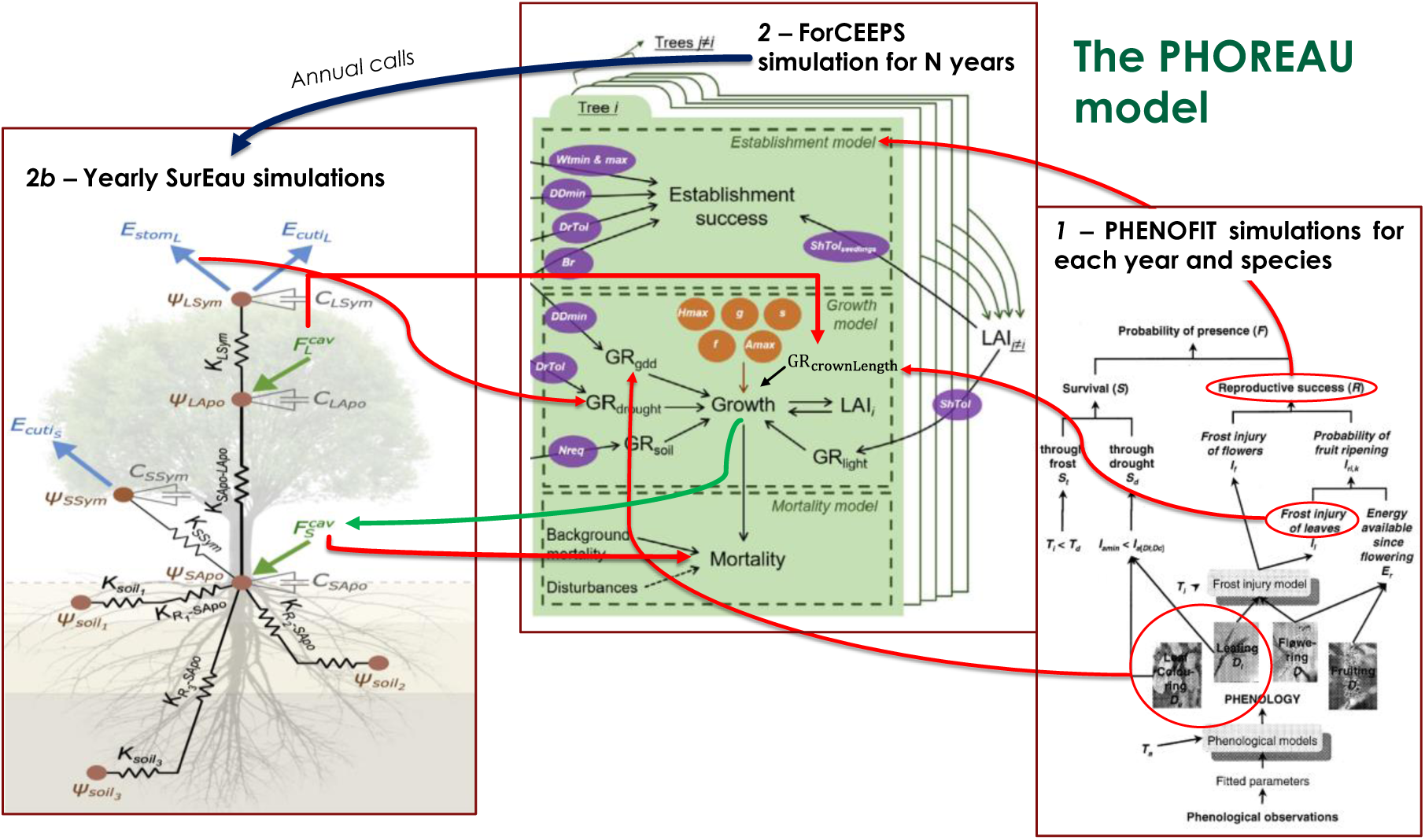
Detailed representation of the processes included in the ForCEEPS and PHENOFIT models. Red circles indicate outputs used for the coupling, and red lines their destination in the ForCEEPS simulation. Original figures are taken from Morin et al. (2021), Chuine and Beaubien (2001), and Ruffault et al. (2022), where parameters details can also be found.

### 2.1 The forest community gap-model

#### 2.1.1 Presentation of the ForCEEPS model

In PHOREAU, forest dynamic processes (growth, mortality and recruitement) are all managed by the ForCEEPS model (Morin *et al*., 2021). ForCEEPS (Forest Community Ecology and Ecosystem Processes) is a gap model that relies on a few ecological assumptions to simulate the dynamics of tree establishment, growth and mortality in independent small patches of land, that are aggregated to derive properties at the forest scale. While the model is not spatially explicit at the patch level, it is individual-based: two trees of the same species and the same age can have different growth rates under the same climate, depending on the specific patch-level biotic constraints of light-competition. Derived from the FORCLIM model (Bugmann, 1996; Didion *et al*., 2009) The ForCEEPS model was developed with the aim of simulating forest dynamics under a wide range of environmental conditions while limiting the need for prior calibration, and was designed to be equally able to simulate planted, managed, or natural forests (Morin *et al*., 2020, 2025). A schematic representation of the different processes and how they interact can be found in Figure 1.

Tree growth is computed at a yearly time-step in two phases. First maximum diameter increment is calculated using an empirical equation as shown in Eq. 1, as a function of trunk diameter at breast height at the start of the year, and a maximum species growth rate *g*_*s*_. *b*_*s*_ and *c*_*s*_ are species specific allometric parameters, and *H*_*max*_ the maximum height reachable by that species. Height is directly linked to diameter following another species-specific allometric parameter.

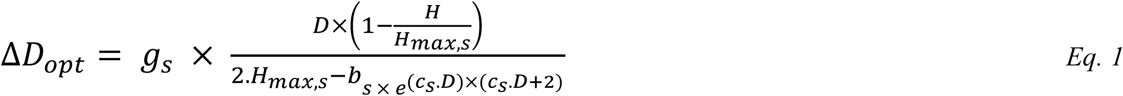

Realized growth is determined from optimal growth after reduction by a series of growth-reduction factors (bounded between 0 and 1) following a modified geometric mean, as shown in Eq. (1)

Drought, growing degree days, and soil reduction factors range from 0 to 1 are determined by site soil and climatic conditions, and modulated by species-specific parameters. The other factors represent biotic constraints related to light availability. *GR*_*light*_ represents the immediate effect of competition for light, and depends on the cumulated leaf area above or at the same level as the considered tree. *GR*_*crown*_ represents the long-term effects of crown size reduction on the capacity of trees to grow and assimilate carbon. In the original ForCEEPS framework, trees crowns were represented as downwards-pointing triangles; the ratio of crown height to tree height *c*_*s*_ is adjusted based on a factor that decreases from a species-specific maximum to minimum value as the tree experiences increasing shade (see Figure 1).

Similarly, tree establishment is regulated by winter temperature, growing degree days, light availability, and stand browsing intensity (Eq. 38). The number of potential seedlings for a given species depends on maximum site density and its shade tolerance parameter, shade intolerant having a greater regeneration potential (Eq. 37). The survival of each potential seedling is controlled by a stochastic process itself regulated by the reduction factors listed above. If selected, the sapling is initialized with a DBH of 1.27 cm. Tree mortality is the combination of a stochastic background process combining stand density and tree longevity, and a growth-related mortality that represents stress-caused tree death linked to biotic and abiotic constraints.

A full description of the ForCEEPS model developed on the Capsis modeling platform (Dufour-Kowalski *et al*., 2012) that was used as a base for this study can be found in Morin et al., (2021). In the following sections, we present new developments that have been included in the ForCEEPS model, independent from the coupling with SurEau and PHENOFIT.

#### 2.1.2 Decoupling tree height from diameter: light-dependent plasticity

The predictive power of gaps models is tied with their representation of stand structure. Yet most classic gap models, including ForCEEPS, do not simulate a dynamic tree height, instead inferring it from the tree trunk diameter through an allometric relationship. It follows that for a given species, every individual follows the same height-diameter trajectory. While this is consistent with the fact most forestry surveys report basal diameter without height, this means that the models cannot represent site effects on maximum height, as well as the effects of competition for light on the height-diameter relationship. In reality dominated understory trees tend to allocate more carbon to height growth than diameter growth. Conversely, trees in low-density or thinned forests have greater diameter growth and slower height growth (Oliver and Larson, 1996). Furthermore, this sensitivity of growth allocation to competition for light is more marked in shade-intolerant species (Delagrange *et al*., 2004).

The effects of competition for light on growth allocation are crucial for understanding stand dynamics, as small initial differences in height tend to increase with time unless corrected by greater height growth. Forest managers have long known that tree maximum height varies from site to site with tree age and density (Fortin *et al*., 2019), and forest growth models often use different height-diameter depending on site conditions (Mehtätalo, Miguel and Gregoire, 2015). Attempts to implement dynamic height growth in gap models have been shown to increase the realism of simulated stand structure, without reducing general applicability. For instance Rasche *et al*. (2012) have implemented such a dynamic height in the ForClim model on which ForCEEPS is originally inspired. Instead of the static relationship between diameter and height (*h*), height increments are calculated at each time-step Δ*H* = *f*_*h*_ Δ*D* through a function *f*_*h*_ that distributes growth between diameter and height growth according to a competition-for-light driven parameter *s*, which replaces the original fixed species-specific allometric parameter. Since the yearly diameter increment uses previous-year height in its calculation, its formulation also had to be adapted to account for the fact that height is dynamic and no longer directly calculated from diameter. These adaptations have been used in our modified ForCEEPS model, albeit with two important modifications.

Firstly, the parameters of the growth-distribution coefficient *g*_*s*_ were adapted to be more conservative, and better reflect the species-specific relationship that had already been parametrized:

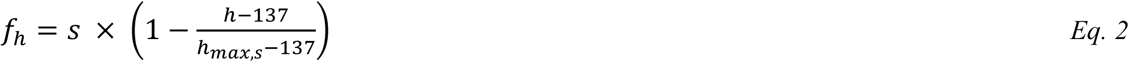

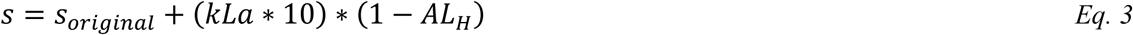

where *kLa* is the species shade-tolerance, *h* the tree height in centimeters, *h*_*max*,*s*_ the maximum species height, and *AL*_*H*_ the light availability at the top of the tree crown.

Secondly, we adapted the yearly growth equation. In the original formulation by Rasche *et al*. (2012), because yearly growth is calculated on the basis of total diameter at the start of the year, a tree that allocated more growth to its height than to its diameter due to competition would have less total growth the following year, all else being equal. This is a result of the simplifications of the ForClim model, in which the diameter increment is calculated on the basis of previous year diameter instead of the previous year volume. This means tree biomass is only dependent on tree diameter, disregarding its height. Given long enough timescales, this effect has major implications, as originally taller but thinner trees end up with smaller final height and diameters than in the original formulation. A possible solution would have been to replace trunk diameter by volume in the growth equations; but this would have meant reshaping the model from the ground up, and making it less applicable to classic forestry datasets, as actual volume data are rarely available. A static set (*D*_*static*_ and *H*_*static*_), calculated from the old equations and static allometry relationships, that were only used as an *ad-hoc* proxy for real tree volume in the updated diameter increment equation (Eq. 4) and the calculation of slow-growth mortality (to avoid killing off trees that allocate too much growth to height); and a real set (*D* and *H*), using the updated equations and dynamic allometry, that was used in all other cases including the light-competition module.

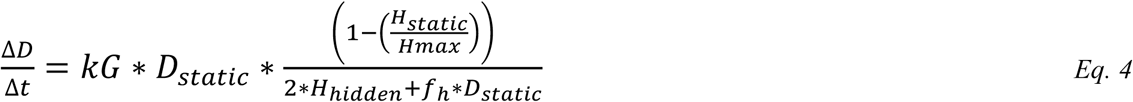

#### 2.1.3 A modified light-competition module

In addition to the dynamic representation of tree height, several modifications were implemented to better represent the complex interactions between stand structure and competition for light. This proved necessary when integrating our mechanical transpiration-driven approach to water fluxes, as stand leaf area is one the main driver of embolism in the SurEau model (Cochard *et al*., 2021b). Preliminary results indicated a poor capability of the ForCEEPS to reproduce observed leaf area indices from stand inventory, in both relative and absolute terms. The refinements to the light-competition module presented below and summarized in Figure 2, were tested with a large-scale validation protocol as detailed in section 4.6.

**Figure 2.**
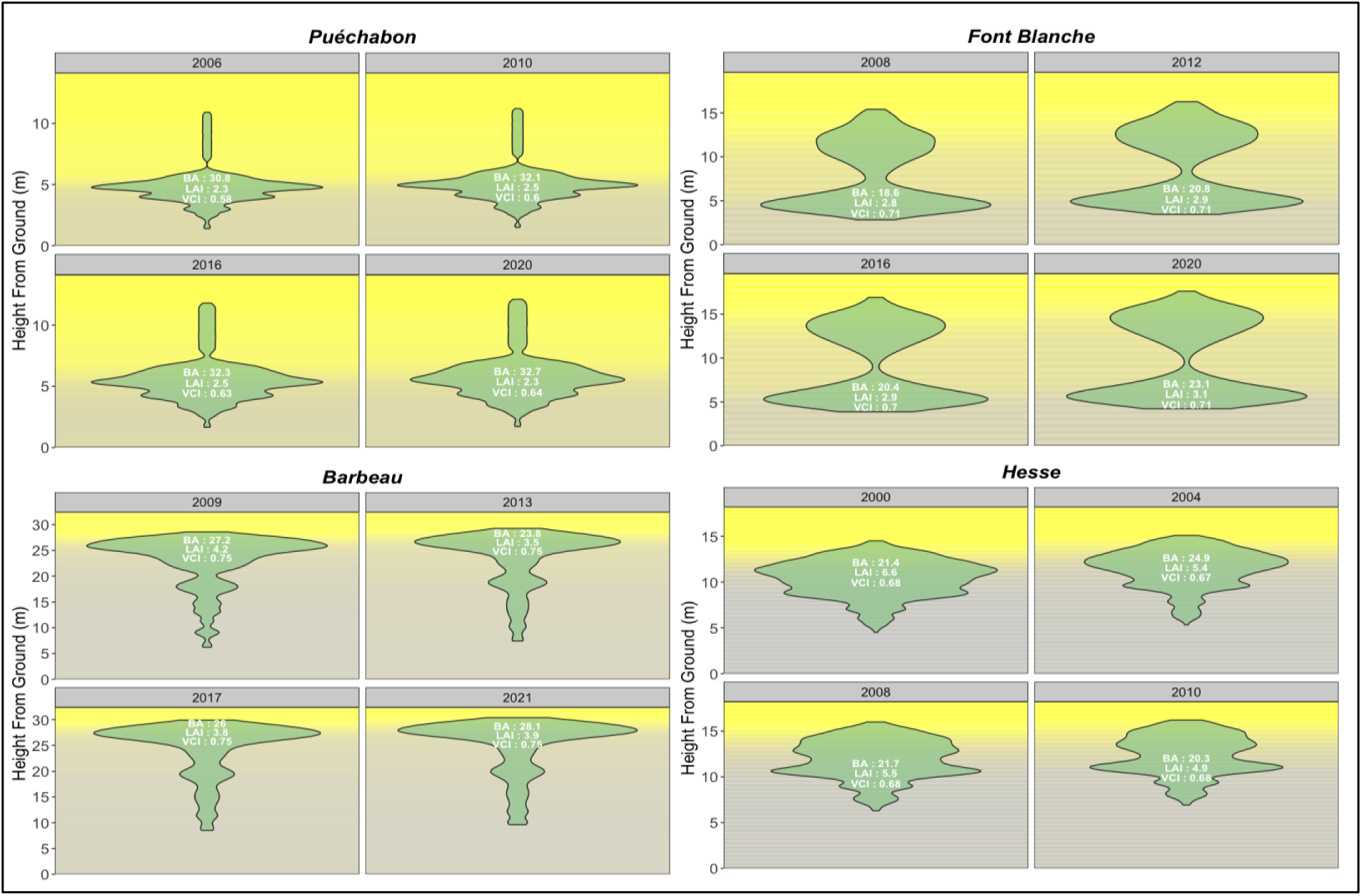
Predicted distribution of stand leaf area and light availability. This figure illustrates the vertical gradient of predicted light availability indices of the four considered ICOS sites for specific simulation years. The light availability is presented over the aboveground profile, divided into 0.1 m layers. In addition, the area of each shape in the layers represents the predicted aggregate leaf area. Refer to Figure X for light availability index gradient. The figure also includes global annual stand parameters. For details on the calculation of the Vertical Complexity Index (VCI), refer to Annex X.

**Figure 3.**
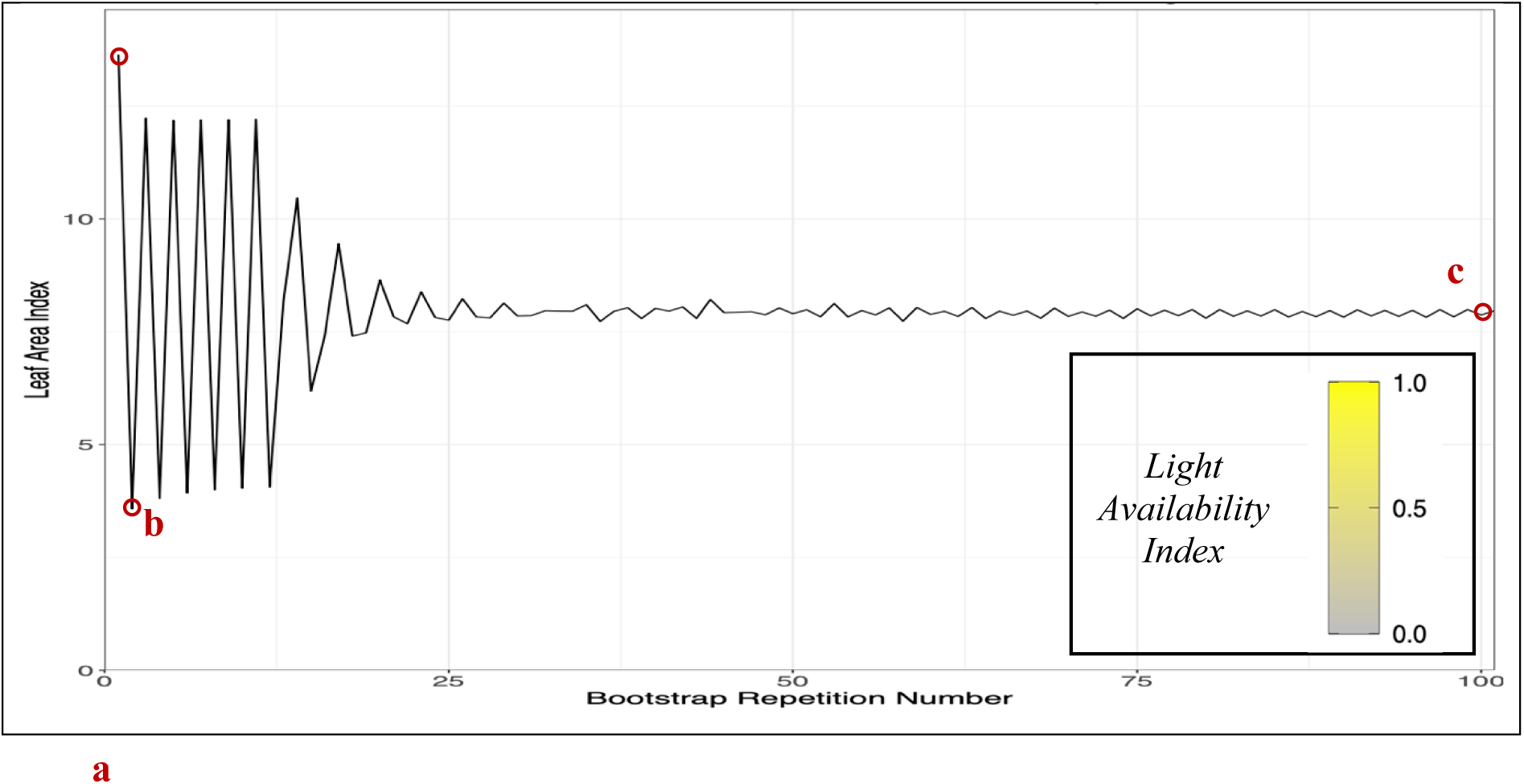
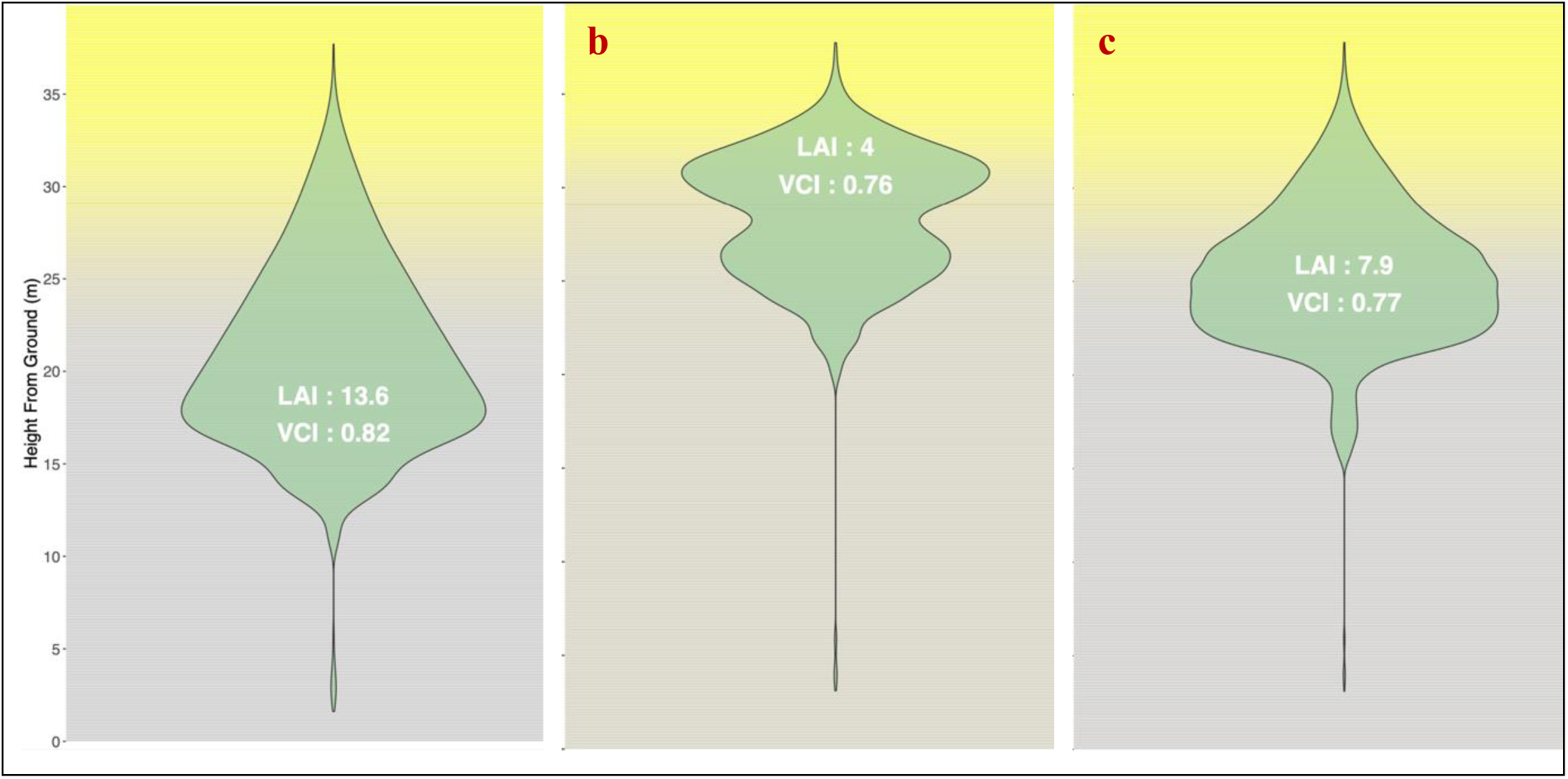
Illustration of PHOREAU canopy bootstrap algorithm. Top: one-sided leaf area indices predicted by the PHOREAU bootstrap algorithm, initialized with a Picea abies dominated inventory (RENECOFOR EPC 39a, 2003). Bottom: three snapshots of predicted foliage area and light availability vertical stratification at different steps in the algorithm. For details on the calculation of the Vertical Complexity Index (VCI), refer to Annex X.

In addition to the dynamic representation of tree height, the evolution of tree crown length was modified to better represent the feedbacks between stand structure and competition for light. In PHOREAU, light availability impacts growth directly and indirectly: directly through the shading growth reduction factor, and indirectly through the crown-length growth reduction factor, which represents long-term crown recession due to shading. Individual tree crown lengths are calculated as the product of tree height, and a variable ratio. It is this ratio that changes according to the light exposition of the tree, between two extreme species-specific values as described in Morin *et al*. (2021). In the original ForCEEPS framework, seedlings started with a crown ratio set at the species maximum, which then decreased over the tree’s lifetime with shading; in particular, this formulation assumes that the crown ratio can only ever decrease or stay the same from one year to the next, with no possibility of reversion when more light becomes available.

It is this possibility of crown ratio reversion we have implemented in PHOREAU. A constantly decreasing crown ratio assumes no increase in light availability over a trees lifetime, disregarding the impact that the death or removal of one tree can have on its neighbours by enhancing light availability and leading to larger crown sizes and denser canopies (see Juchheim, 2020, and Saarinen *et al*., 2022). We have consequently adapted the original ForCEEPS crown ratio equation to reflect this, with a yearly increase capped at 5% of the difference between the previous-year crown ratio, and the potential crown ratio given current light availability. We are aware this approximation does not take into account the fact that younger trees recover their crowns better due to having more remaining growth potential (Hynynen, 1995).

#### 2.1.4 The boostrap algorithm

In the PHOREAU framework, the leaf area is updated at the end of the year, after each tree’s crown length has been updated according to the light availability. However, the light availability that is used to calculate the new crown lengths is the result of the stand area of the previous year, which is itself the result of the previous year’s crown lengths. This asynchronicity means that – disregarding other processes like growth regeneration and mortality – the estimation of stand area will oscillate around an equilibrium state. While this equilibrium state is dynamically stable, the oscillations for the first few years are large enough to be significant. This is especially problematic when starting the model from an inventory: because actual crown lengths are rarely available, the model is forced to initiate the crown at the maximum species’ value; the resulting very low light availability means that the following year the crown lengths will be reduced by a large factor, which means that more light will be available the year after that, causing a new spike in stand leaf area. It is to correct for this effect that we implemented a bootstrap algorithm where, before the first year of the simulation, multiple iterations of the light competition module are run until the shift in stand area between two successive iterations becomes negligible.

#### 2.1.5 Phenology-based temporal niche partitioning for light

In ForCEEPS, the way the light availability of each canopy layer is determined by the above total leaf area of the other layers, combined with differentiated shade tolerances between species, allows emergent complementarities between in a multi-specific context between shade tolerant and intolerant species, resulting on average in greater total stand leaf area and productivity at the stand level. But alongside spatial complementarities, there exist temporal complementarities in species usage of light linked to their phenology (Gotelli and Graves, 1996).

The PHOREAU model, by integrating leaf phenology simulated in PHENOFIT, is able to account for these temporal effects. This required an in-depth reworking of the light-competition module. Instead of calculating layer light availabilities at the yearly time-step, daily light availabilities are now calculated by summing the crown areas of the taller layers excluding dormant trees (this assumes that trees are either dormant or active: in reality leaves bud and fall over the course of a few days, and senesced leaves can cast shadow for weeks before falling off). The final tree light availability is calculated by summing, over all its layers, for all the days for which it is itself active, each daily layer light availability. To correct for the fact that tree growth is dependent on heat as well as sunlight, this sum is weighed using daily mean temperatures. Finally, the leaf areas used in the light availability calculations are reduced by the PHENOFIT leaf loss index in case of frost.

#### 2.1.6 Species-dependent crown shapes

An accurate representation of crown shapes is an integral component to any model of light competition and canopy interactions between trees (Krůček *et al*., 2019). In reality the crown shape of any given tree is a complex combination of genetic, allometric, and environmental factors, as crown shape varies across species, age groups, climate, local conditions and the shading status of the tree (Oliver and Larson, 1996). Canopy packing in mixed forests can be partly attributed to this heterogeneity and plasticity of crown shapes, as trees suffer relatively less competition for a given foliage density (Longuetaud *et al*., 2013).

Crown-shape representation in PHOREAU iterates on the ForCEEPS framework, which already allowed for stratified distributions of foliage area over a vertical axis (Morin *et al*., 2021). Compared to the previous iteration, PHOREAU allows trees to have other crown shapes than the default inverse-cone – such as conical or ellipsoidal shapes. This is meant to represent broad patrons in crown geometry observed at the European Scale, such as the fact species present in higher latitudes or latitudes tend to have more columnar or conical crowns to capture light coming from a perpendicular angle, whereas species as lower latitudes are more frequently flat-topped for maximum exposure (Kuuluvainen and Pukkala, 1989).

While the lack of explicit tree positions prevent PHOREAU from recreating the asymmetrical crown shapes which result from horizontal constraining between crowns (Niklaus *et al*., 2017), this simple approach allows for a more accurate representation of side-shading between trees, and captures the way shaded trees tend to become more flat-topped as they reduce their crown height (Oliver and Larson, 1996), while saving some simulation time. See Figure 11 for a visualization of the new crown shapes.

#### 2.1.7 Density-dependent light availability

Any representation of forest canopies and light dispersion has to strike a balance between predictive power — how much photosynthetically active radiation (PAR) does a given tree actually receive at a given moment in time? — and computing cost: by aggregating leaves on a tree-by-tree basis and disregarding differences in angle and light absorption between sun and shade-leaves (Givnish, 1988), by calculating at yearly time-step, and by considering only the vertical stratification without an explicit representation of trunk distribution across space, ForCEEPS is able to compute in a timely fashion what would otherwise take orders of magnitude longer with a more bottom-up approach from the leaf to the tree.

PHOREAU does not diverge from this general framework, which is well suited to working on large-scale inventories (that usually come without tree-level coordinates), and does not suppose any *a priori* knowledge on canopy composition. However, this simplification is not without its drawbacks. Because the light availability of a given canopy layer depends solely on the foliage area present in the layers above it, with no accounting for how this foliage is actually distributed, light competition is — in effect — boiled down to a single value: the LAI. Intuitively we understand that this does not quite tally with reality: two superposed leaves will intercept less light, all else being equal, than two leaves on a level plane; forests are not horizontally homogeneous, and gaps in the canopy may form as trees die off, allowing saplings to sprout and grow even in dense stands (Nicotra, Chazdon and Iriarte, 1999). Due to the links between patchy structures of light availability and tree species diversity and coexistence (Moora *et al*., 2007), measuring and quantifying microsite light availability has been a focus of research (Parent and Messier, 1996; Tymen *et al*., 2017), with important implications for forest management (Coates *et al*., 2003).

This structural limitation — which can be important, e.g. to accurately predict species richness in relation to management — can never be fully worked around. And, in keeping with the general philosophy of the model to strike a balance between complexity and genericity, we opted not to incorporate a complex 3D tree-level light absorption model (le Maire *et al*., 2013). However, in the transition from ForCEEPS to PHOREAU, some steps have been taken to at least partially account for the horizontal stand structure. This was done in an indirect way by using information available to the model: the stand density.

As in most gap-models, foliage area in ForCEEPS is translated into light availability using a modified logarithmic Beer-Lambert law, see Eq. 5, where light availability is a function of foliage area and a light extinction coefficient *λ*. In the original formulation of the law this extinction coefficient is calculated by integrating over the path of the light ray the absorbance and density of the materials it crosses. This calculation — which accounts for the angle of the leaves, the angle of the sun’s rays, the different absorbances between species and sun and shade-leaves, and the distribution and clumping of the leaves and trees (Smith, 1993; Dufrêne and Bréda, 1995) — is usually simplified into an empirical constant extinction parameter, which can vary from site to site (Vose *et al*., 1995; Binkley *et al*., 2013). However, in the ForCEEPS framework, where stand composition is an emergent property and not an input, a single

#### *λ* value is used regardless of site conditions

Following the methodology outlined in (Nilson, 1971; Black *et al*., 1991; Bréda, Soudan and Bergonzini), PHOREAU integrates a clumping factor Ω in its calculation of the light extinction coefficient. This clumping factor ranges from 0 (corresponding to a fully concentrated distribution) to 1 (corresponding to a perfectly homogenous distribution), and represents the aggregation of leaves within each tree and between the trees themselves. The advantage of this approach is that Ω can be calculated each year as an emergent variable, allowing the model to capture observed trends like the inverse relation between LAI and the light extinction coefficient (each additional increment of leaf area blocks marginally less light) (Dufrêne and Bréda, 1995). The clumping factor in PHOREAU is calculated using Curtis relative density (Smith, 1993; Curtis, 1982): with this formulation (see Eq. 6) for a given LAI, a dense stand with small trees will block out more light than a stand populated by a few large trees. This approach is similar to the one used in LAI estimation with MODIS or hemispherical photography, where clumping indices are also used to correct the raw measured LAI (Demarez *et al*., 2008; Chen *et al*., 2012; Zhu *et al*., 2018).

A further step would be to incorporate species-specific absorbance values, as leaves of different species react differently to incoming light (Binkley *et al*., 2013), but this would necessitate gathering data at the species level. Another possible refinement would be to incorporate the angle of incoming light in the calculation of light availability (Smith, 1980); but this would require additional inputs, in the form of site slope and exposition.

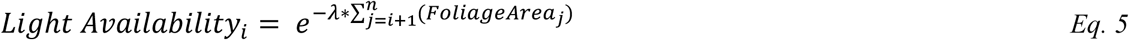

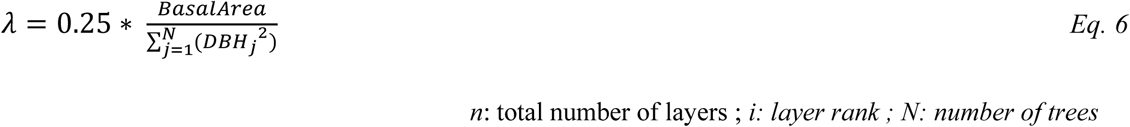

#### 2.1.8 Incorporation of Specific Leaf Area

The relation between trunk diameter, crown biomass, and foliage area in ForCEEPS are governed by a set of simple allometric relationships calibrated for a few of the main temperate European species, using experimental data collected in Switzerland by destructive sampling in the 1940s and 50s (Burger, 1951; Bugmann, 1996). The refinements that ForCEEPS implemented regarding crown plasticity and explicit vertical stratification were built upon this foundation but did not challenge its underlying assumptions (Morin *et al*., 2021). This became problematic as the model — and PHOREAU in particular — incorporated more species from a larger geographic range: understory or Mediterranean species in particular that were not represented in the initial calibration dataset. This was directly reflected in model predictions, for example with an overestimation of *Quercus ilex* or *Pinus halepensis* mortality due to inflated foliage areas.

A simple solution to this issue was implemented by recalculating the *c*_2_ parameter (used in ForCEEPS to derive a tree’s foliage area from its diameter) using a specific leaf area (SLA) value for each species.

The retained SLA — the surface area for a given mass of leaves — are those of average adult individuals of each species over a large set of sites (Kattge *et al*., 2020; Devresse *et al*., 2024). This new formulation, see Eq. 7, allows the model to capture inter-specific differences in drought resistance strategies (Greenwood *et al*., 2017), while disregarding for the moment SLA plasticity to tree age, competition, and site conditions (Gratani, 2014).

#### 2.1.9 Climate-induced leaf loss

While ForCEEPS already implemented a mechanism for competition-driven loss of foliage area, representing the reduction of the crown height of dominated trees as their lower branches die off, this did not adequately represent climate-induced leaf loss. Unlike competition-driven branch dieback, leaf-loss caused by extreme weather conditions is not usually accompanied by branch death, does not preferentially target the leaves located in the lower parts of the crown, and can be more quickly reverted with shoot regrowth. These differences justified the implementation in PHOREAU of a new mechanism for transitory leaf-loss, distinct from the reduction of crown size, with no memory from one year to the next. The variables used to drive this leaf-loss are derived from the yearly percentage of uninjured leaves (*I*_*l*_) and leaf cavitation (*PLC*_*l*_) values calculated respectively in the PHENOFIT and SurEau submodules

(see sections 3.2 and 3.3), and representing the impact of frost and drought events. This new mechanism, see Eq. 8, allows the model to reflect strategies of drought acclimation, where defoliation can help some species tolerate drought events (Bréda *et al*., 2006; Limousin *et al*., 2022) at the cost of a lowered growth potential.

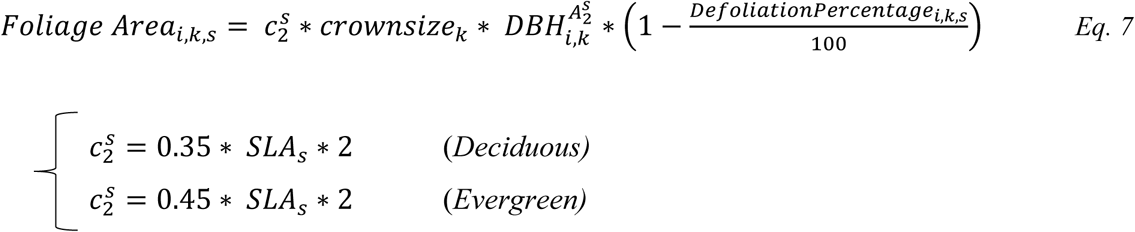

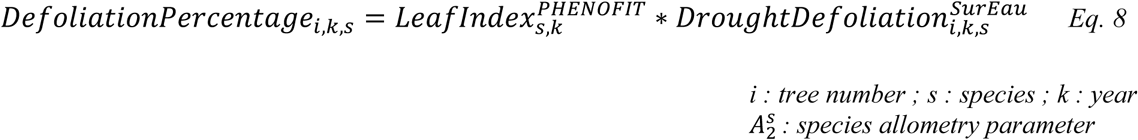

#### 2.1.10 Microclimate derived from stand-structure

By integrating fine hydraulic and phenological mechanisms in the overall framework of a forest-structure gap model, PHOREAU has the opportunity to capture the effects of microclimate on plant functioning. Because forest canopies absorb or reflect the majority of incoming solar radiation, reduce wind speeds, convert solar energy into latent heat through evapotranspiration, and block outgoing infrared radiation, climatic conditions in the understory are often buffered compared to those at the top of the canopy, with cooler more stable temperatures during the day, and warmer temperatures during cold nights. This climate dampening effect is more marked for temperature extremes, and for tall, structurally complex dense canopies (De Frenne *et al*., 2021). Furthermore, it is an important factor in ability of young, understory trees to resist droughts despite their shallow root systems (Forrester and Bauhus, 2016). Because PHOREAU evaluates drought-stress at an individual level by calculating tree fluxes, it can easily make use of microclimatic data for temperature, air humidity, and light availability, to better compute plant evapotranspiration and in turn differentiate water stress among individuals of different heights. In addition, because PHOREAU simulates many small patches each sharing a soil and a canopy height profile, the incorporation of microclimate could help the model capture forest landscape mosaic dynamics, where forests with heterogeneous patches are able to host more diversity due to differentiated microclimatic effects on regeneration and drought (Pincebourde *et al*., 2016).

To derive microclimate temperature and air humidity from macroclimate, we implemented a version of the statistical model developed, calibrated and validated in Gril *et al*., (2023) and Gril, Laslier, *et al*., (2023). This model, which has the advantage of using only easily available patch characteristics, uses a simple slope and equilibrium approach, presented in Figure 5, to compute microclimate temperature at soil level (*T*_0_) from the corresponding hourly or daily macroclimate temperature (*T*^*j*^). The slope (*m*_*slope*_) captures the linear relationship between microclimate and macroclimate, while the equilibrium is the point at which microclimate is equal to macroclimate (Eq. 11). In our case, month mean temperature (*T*^*m*^) is used as the equilibrium. The slope, which acts as a buffer if is lower than 1, is computed daily using patch-level leaf area index (*LAI*), maximum tree height (*h*_*max*_), and vertical complexity index (*VCI*), as seen in Eq. 12, with corresponding coefficients calibrated over a large dataset of microclimate measurements (Gril, Laslier, *et al*., 2023). VCI is obtained following Van Ewijk, Treitz and Scott, (2011) by calculating the weighted logarithmic average of foliage area proportion per patch canopy layer (*p*_*i*_), normalized by the total number of layers *n*, as shown in Eq. 13 and Eq. 14. Finally, for any given tree height *h*, the corresponding microclimate temperature 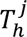 is derived from soil microclimate and macroclimate using a linear interpolation, as shown in Eq. 9 and Eq. 10.

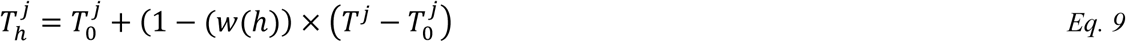

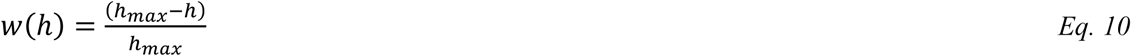

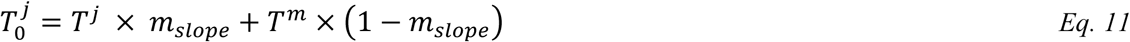

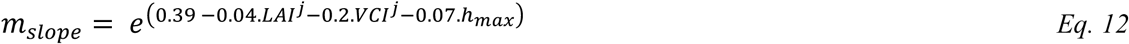

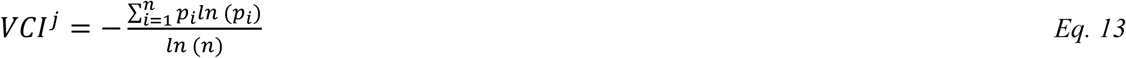

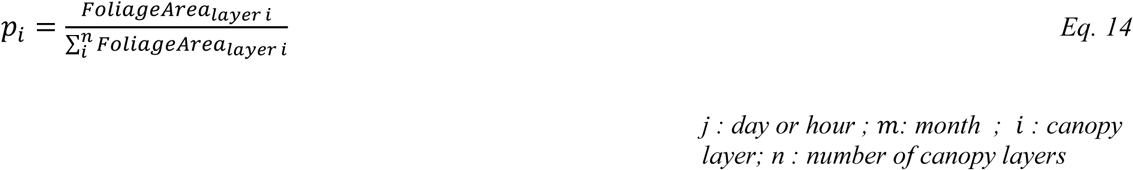

**Figure 4.**
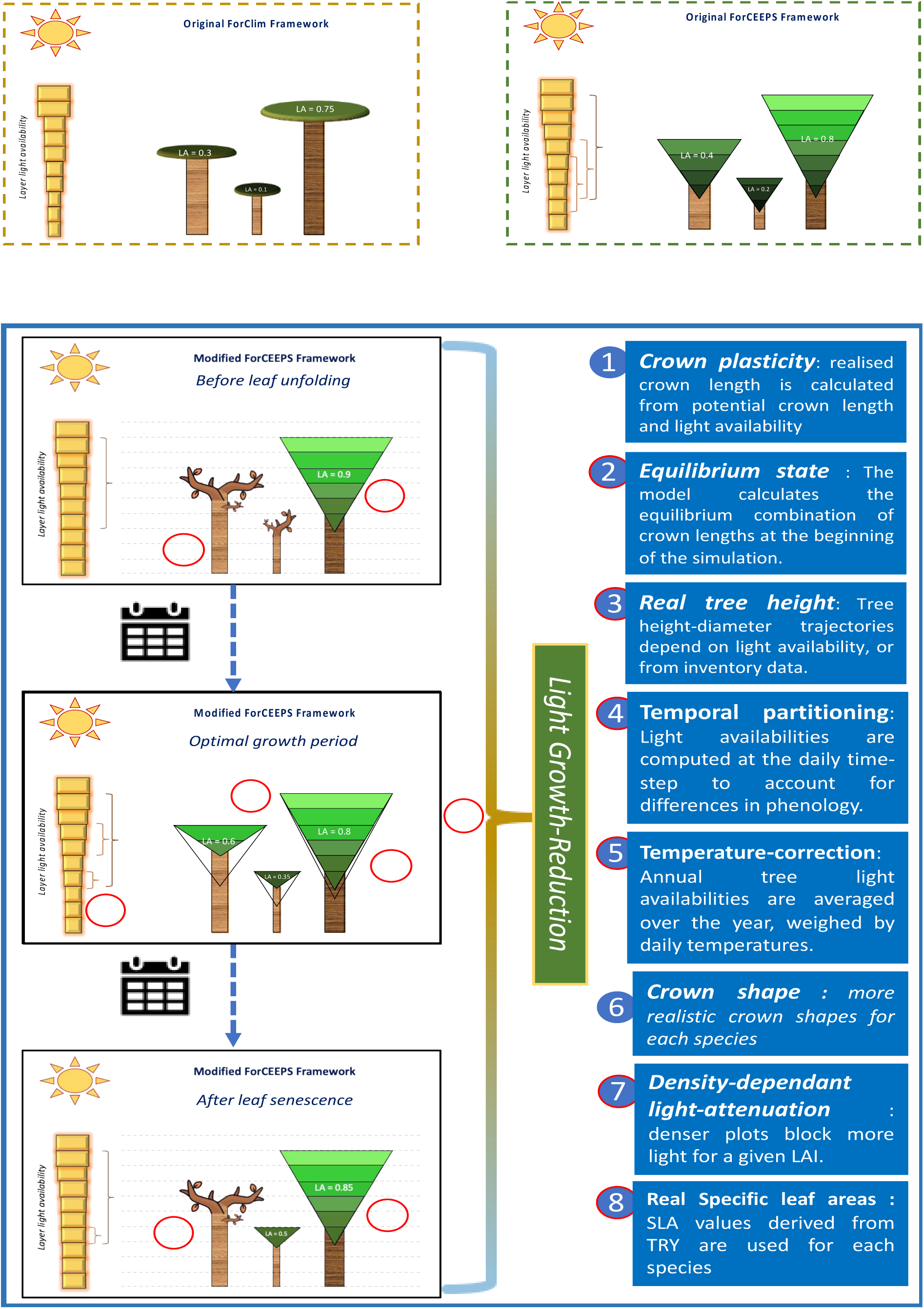
Presentation of the modifications in the light-competition module between ForCEEPS (Morin et al., 2021) and PHOREAU, with a description of the main changes.

**Figure 5.**
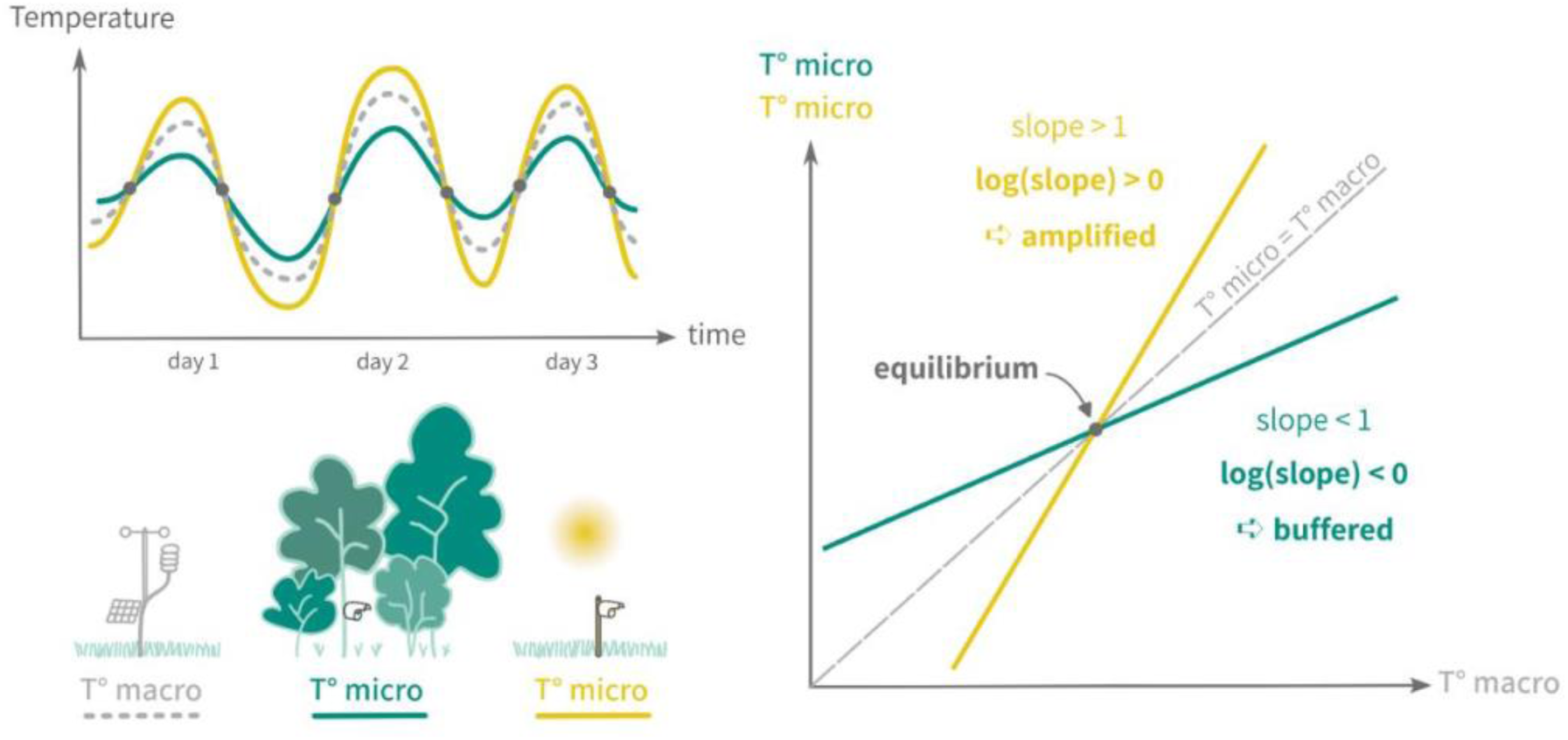
Schematic representation of the slope and equilibrium microclimate approach, reprinted from Gril, Laslier, et al., (2023).

Calculated hourly microclimate temperatures are then used to compute corrected local vapor pressure deficits (VPD) used in PHOREAU transpiration computations. These temperatures are also used in GDD calculations (see Eq. 32), as well as for seedling establishment constraints based on minimal temperatures (*W*_*Tmin*_). For seedlings, soil-level microclimate temperature is directly used; for established trees, the microclimate temperature is calculated the weighted average height of their foliage area distribution.

Because leaf unfolding and senescence dates are integrated in the calculations of *LAI* and *VCI*, the slope of microclimate buffering or amplification can change throughout the year.

While this approach presents a number of advantages, it comes with major simplifications. The most important one is certainly the linear interpolation of microclimate over the height of the stand, which neglects actual wind movement and radiation attenuation dynamics. Microclimatic data, measured at different heights below the canopy, would be needed to calibrate a more realistic non-linear function. Other simplifications include disregarding the effect of soil moisture, ignoring horizontal heterogeneity within patches, and assuming monthly mean temperatures are a good indicator of equilibrium.

### 2.2 Integrating plant hydraulics

At the heart of the PHOREAU model is the complex integration of the ForCEEPS gap model with the SurEau plant hydraulics model. This integration was made possible by the presence of both models on the Capsis Java platform (Dufour-Kowalski et al., 2012). The version of SurEau implemented on Capsis closely resembles the framework of the SurEau-Ecos v2.0 (Ruffault et al., 2022). which was developed in R as a more parsimonious alternative to the original SurEau model for large-scale applications, with a simplified representation of plant architecture, and a modified numerical scheme to reduce computing time. However, the capabilities of the Java object-oriented language allowed a number of additions compared to the R framework, presented below: it is this on the basis of this more modular, more efficient version of SurEau, that the integration with ForCEEPS was made possible, allowing the user to finely tune the level of detail by changing the number of tree organs, the time-step, and even to switch between the “implicit” (SurEau-Ecos) and the standard calculation method for water fluxes.

#### 2.2.1 Presentation of SurEau-Ecos

A detailed presentation of the principles of the SurEau model can be found in Cochard *et al*., (2021) and Ruffault et al., (2022). It is a process-based hydraulic model, in which the plant is represented as a series of organ connected like a circuit, with resistances and capacitances regulating water flow between the different reservoirs. Each plant organ — roots, trunk, stems and leaves in the original complete model — is itself separated in two compartments, with their own capacitances and resistances: the apoplasmic compartment is linked to its organ’s symplasm as well as to apoplasms of neighbouring organs, while each symplasmic compartments is linked to its corresponding apoplasm, and to the atmosphere through cuticular transpiration. In addition, each leaf symplasm is connected to an evaporative site, while root symplasms draw water from soil reservoirs through an endoderm. In the simplified model used here the system is discretized as an atmosphere, three soil layers, and only two plant compartments. The plant is represented as a leaf and an abstract “stem”, which is an aggregate of real water volumes in plant stems, trunk, and roots, and as such is directly connected to the soil layers (Figure 6).

**Figure 6.**
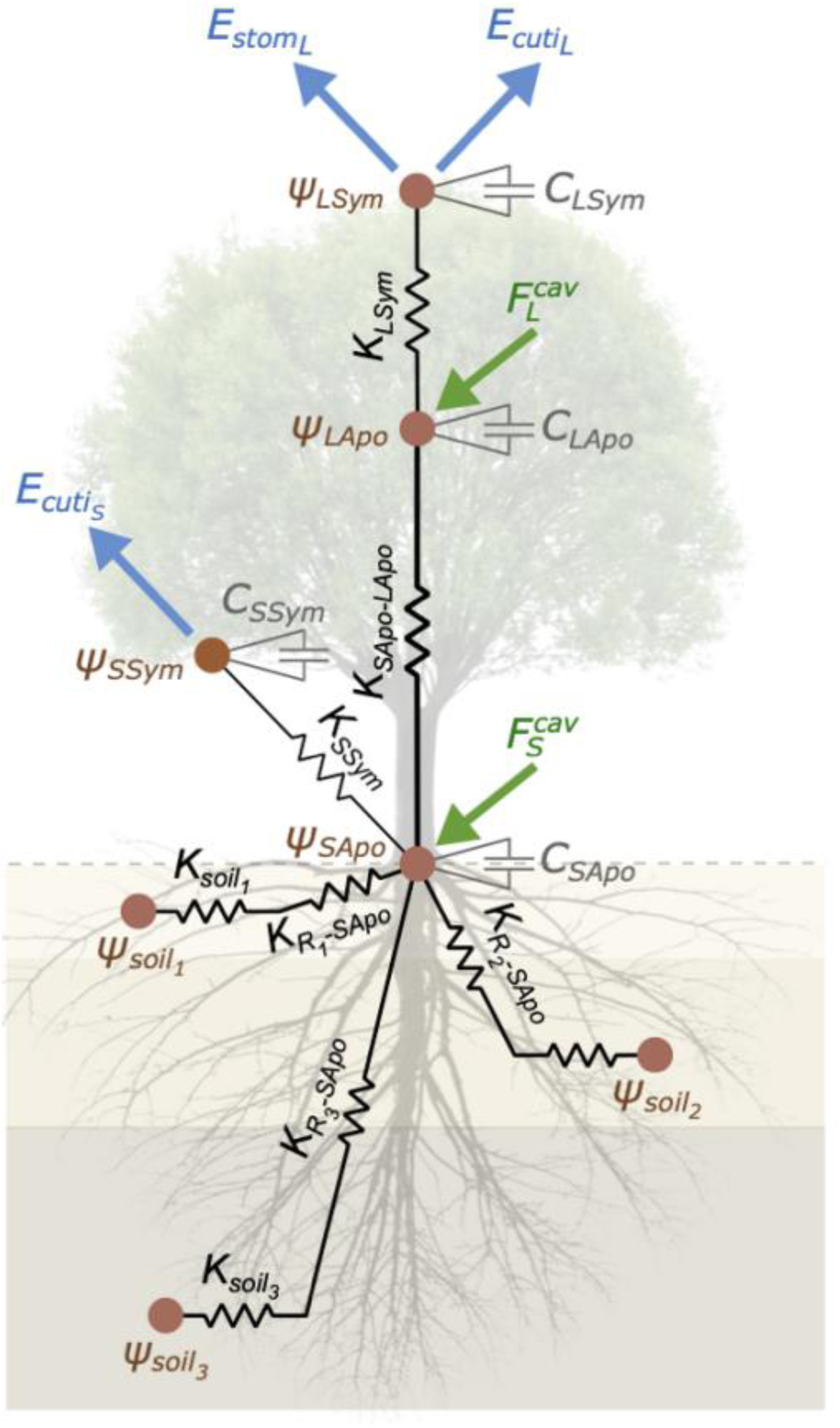
The SurEau framework, Ruffault et al. (2022)

**Fig. 7.**
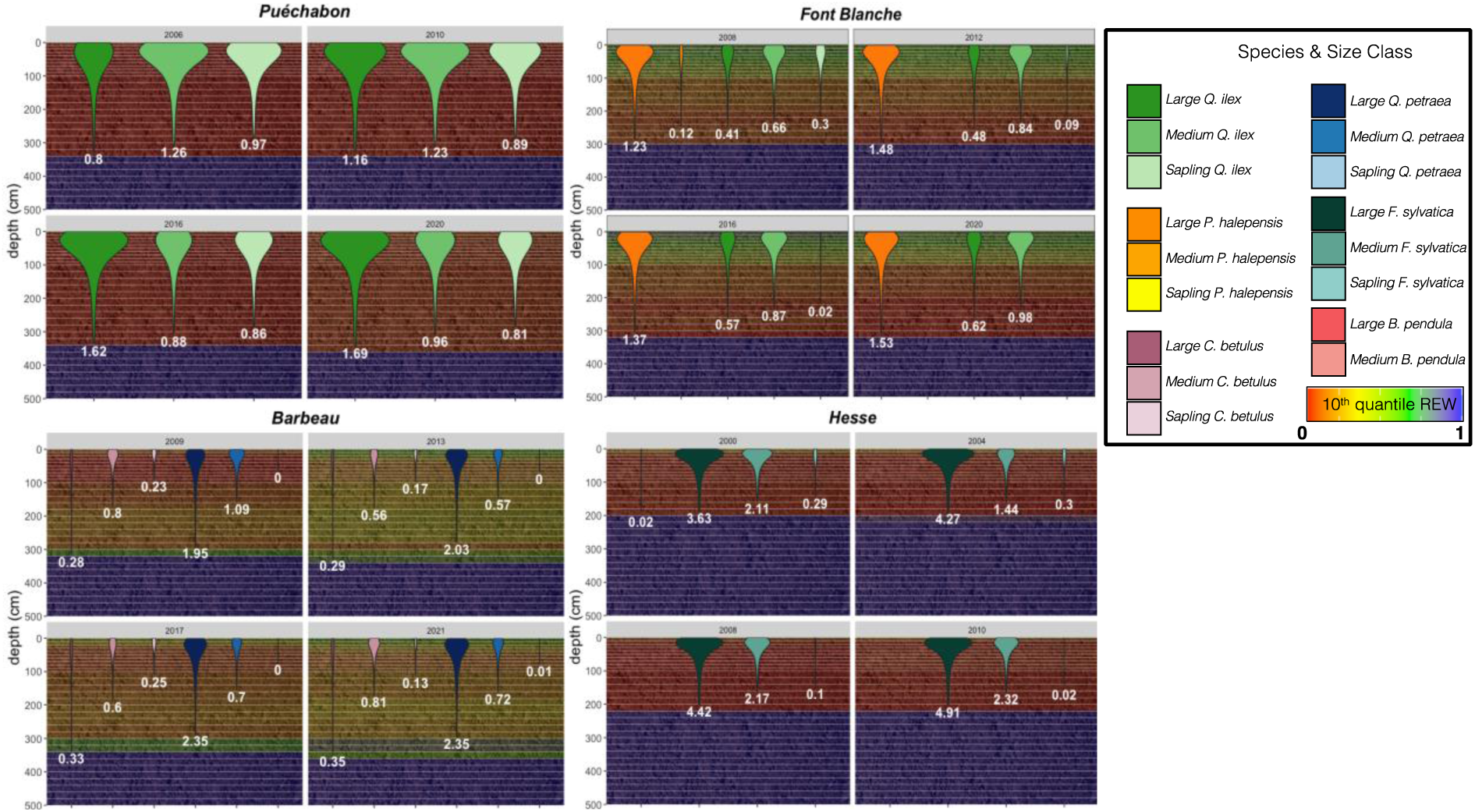
Predicted fine root area distribution over the soil profile. For the four ICOS validation sites, for certain simulation years a partial vertical sol profile is shown, with the overall dryness of each sol layer depicted as a gradient using its 10” quantile relative extractable water (REW) percentage. For each species and size class aggregate tree (refer to Annex X for details on the aggregation method), the distribution of the inverse cone along the soil layers represents the predicted location of it fine roots, with its total aggregate tine root area index (FRAU shown under.

The model functions at two distinct time-steps, configured here as a large hourly time-step, and a small 1200ms time-step. The short-time step computes water fluxes and integration of water balances, while the longer time-step computes transpiration, cavitation, leaf energy balance, and other processes that do not directly interfere with hydraulics.

At the short time-step local water balance is the result of water fluxes and compartment specific sink terms (soil transpiration, cuticular and stomatal evaporation) and source terms (precipitation, water release through cavitation). The overall flow of water driven by transpiration at the stomatal and cuticular level of all organs. Water-flows between compartments are computed as the product between the interface conductance and the gradient of water potential. Water potentials are the result of the difference between current water content and water content at full saturation on the one hand, and organ capacitance on the other (constant for the apoplasm, computed using pressure-volume curves for the symplasm). Meanwhile interface conductance depends on the level of cavitation and percent of leaf fall.

The percentage of leaf and stem loss of conductance (PLC) through vessel embolism is calculated at the long time-step, using the water potential of the organ’s apoplasm (*ψ*_*Apo*_) and an empirical sigmoid function described by species-specific inflexion and slope parameters (*P*_50_, *slope*_*cav*_) as shown in Eq. 15.

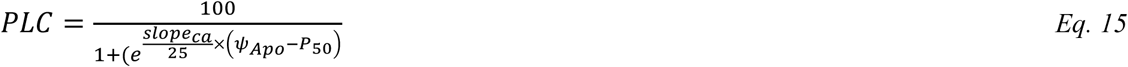

At the long time-step the various cuticular and stomatal transpirations are also computed using gas-phase conductances, and the vapor pressure deficit between the organ and the atmosphere. The leaf stomatal and cuticular conductance are connected in parallel to produce the leaf conductance, itself connected in series to other boundary and crown conductances to produce the overall canopy conductance. Leaf cuticular conductance varies with leaf temperature and its photosynthetic activity. Meanwhile, stomatal conductance is calculated as the product of a maximum stomatal conductance without water stress ((*g*_*stom*,*max*_) which depends on depends on light, temperature, and CO2 concentration, with a regulation factor *γ* based on plant water status, as shown in Eq. 16.

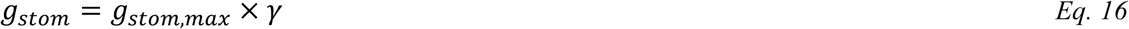

In particular *γ* represents the degree of stomatal closure between 0 and 1, computed using leaf symplasm water potential *ψ*_*Lsym*_ and a sigmoid function described by species-specific inflexion and shape parameters (*ψ*_*gs*50_, s*lope*_*gs*_), as shown in Eq. 17 (Equations are extracted from Ruffault et al., (2022), where a detailed description of the model can be found).

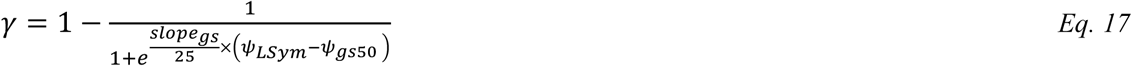

The <u>SurEau</u> model uses daily climate data as inputs, which are then disaggregated into hourly values. In addition, it also accounts for soil properties, stand characteristics, and species-specific traits. Among its outputs are the time to full stomatal closure, and the hourly level of cavitation of each organ. Importantly, this means the model is able to capture the effect of water-stress on the plant after the point of turgor loss, until xylem hydraulic failure and eventual plant desiccation. During a drought event, stomatal closure slows but does not stop the decline of xylem pressure; cavitation occurs when water transported under negative pressure undergoes a phase change from liquid to vapor, hindering the plant’s ability to transport water to photosynthetic sites.

The SurEau has been used with various future Radiative Concentration Pathways scenarios to demonstrate a potential increase in future embolism rates. Using constant climates with no precipitations, the model has also been used to test which parameters explain the variability in the resilience of trees to extreme drought, showing that stand-based traits (i.e, canopy conductance, LAI, soil available water) have the most impact on time until stomatal closure, while plant survival time after stomatal closure is mostly driven by plant-based traits that regulate cuticular transpiration and embolism resistance (Ruffault *et al*., 2022 and Choat *et al*., 2012).

#### 2.2.2 From ForCEEPS to SurEau: climate disaggregation & tree aggregation

A major consideration when developing PHOREAU was limiting the increase in computing time that the integration with SurEau would inevitably bring. In this context, several levels of aggregation were considered.

In its simplest, unrefined state, the connection between the two models can be described as follows. At the beginning of each year, all the trees currently present in the ForCEEPS plot are used to initialize a SurEau simulation. This simulation lasts exactly one year, using the same daily climate (albeit with a further hourly disaggregation) that is used in ForCEEPS. In addition to species hydraulic traits parameters (see Ruffault *et al*., 2022), shoot-specific variables, including height, diameter, leaf area, PLC, and light availability, are retrieved directly from ForCEEPS; leafing and senescence dates are obtained from PHENOFIT; and the state of the soil at the end of the previous is recovered from the previous SurEau simulation. On the basis of these inputs SurEau is run for one year and, at the end of the simulation, information is sent back from SurEau to ForCEEPS that will serve to determine tree growth and mortality (see section 4).

While this approach is workable for simulations on few sites, at shorter time-scales, it quickly becomes unwieldy when considering large datasets. This is a result of the complexity differential between ForCEEPS and SurEau, the running time of the latter representing a roughly hundredfold increase compared to ForCEEPS alone, even with the relatively large 1200 seconds time-step used to compute water fluxes. To account for this limitation, two main simplification schemes were implemented, each optional, presented in order of priority.

##### 2.2.2.1 Treewise aggregation

Because the runtime of a SurEau simulation is driven by the number of distinct water-holding compartments — the atmosphere, soil layers, and mostly importantly tree organs — the first step to reducing the runtime of a SurEau simulation is to reduce the number of initial trees. This approach requires that the global stem volumes and foliage areas remain the same at the stand level, as these are the main drivers of water-use in SurEau and *in natura (Wullschleger, Meinzer and Vertessy, 1998).* The aggregation method ensures this through by summing and averaging, at the cost of some precision in the description of the competition for water.

The degree of simplification is specified at the start of the PHOREAU simulation by choosing a number of *classes*: this is the maximum number of aggregate trees created per species at the start for each SurEau run-year. It follows that, for example, a three-class aggregation in a stand with 4 species will result in SurEau initializing with at most 12 trees, which is a more manageable number. To preserve the overall structure of the stand, trees are distributed within classes on the basis of trunk diameter: for an *n*-class aggregation, for each species, the range of diameters between 7.5 cm and the largest diameter at breast height is decomposed between *n* − 1 segments of same size: classes are then created by grouping all the trees with a diameter at breast height located between the extremities of a given segment, and the last class is composed of all the juvenile trees smaller than 7.5 cm. A consequence of this method is that a class may contain no tree for a given year, and that trees may move between classes from one year to the next as they grow in size.

After the distribution, a single aggregated tree is created for each class. The volume of this aggregate tree is the sum of the volumes of all the trees in the class; its height the average of their heights; its foliage area the sum of their foliage areas; its root depth the average of their root depths; its root biomass the sum of their root biomasses; and finally its light availability the average of their light availabilities. See Fig. 8 for an example case.

**Figure 8:**
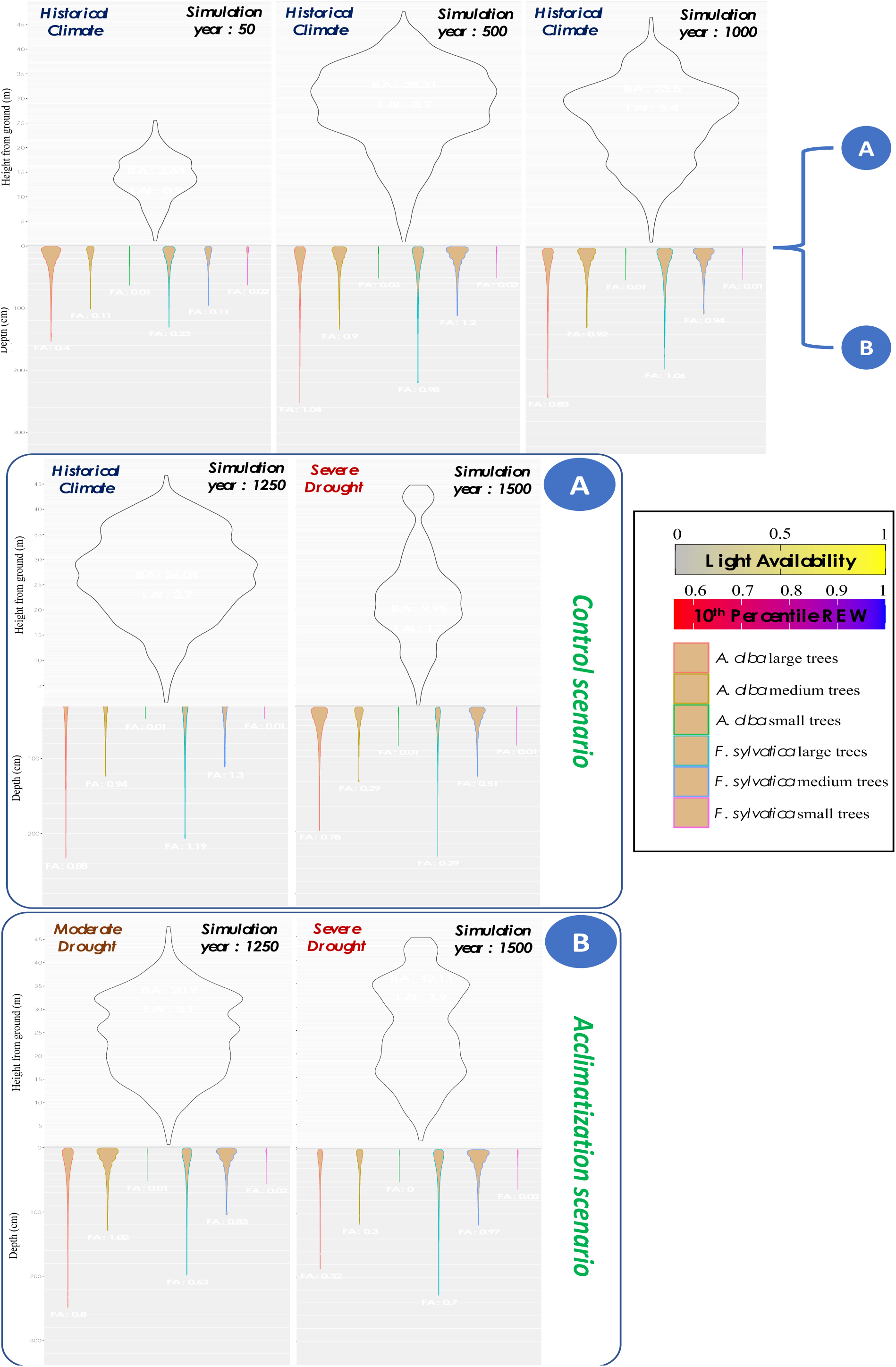
Drought acclimatation proof of concept. Simulated distribution and sum of basal area (m^2^/ha), LAI and fine root area (m^2^/ha) are shown at different steps of two 1500 yearlong simulation over 50 independent 1000m^2^ mixed F.sylvatica and A. alba stands, starting from bare ground, using edaphic and climatic conditions from a beech-dominated Vosges stand (RENECOFOR HET 88, 1969-2020). Yearly climates were sampled at random from historical data. For both the control (A) and acclimatization (B) scenarios, incoming rainfall was reduced by 66% from the 1250th year onward. For the acclimatization scenario, an additional 33% reduction was applied between the 1000th and 1250th year.

##### 2.2.2.2 Dry-year selection

The second optional way of optimizing PHOREAU performance revolves around modifying the rate at which SurEau is called from ForCEEPS. By default, the two submodels are run on a 1-to-1 basis, with SurEau being called at the beginning of each year; but a more parsimonious approach is to run SurEau only for the driest years of the simulation. This simplification is based on the idea that the impact of drought on forested stands, and especially on tree mortality, does not follow a linear curve, but rather depends on climate extremes, physiological thresholds and tipping points (Hartmann *et al*., 2018). Because this approach requires a prerequisite ranking of all of the years of the simulation according to their dryness, we use an integrative Drought Index calculated for each year (Morin *et al*., 2021). The rate of SurEau calls — every two years, five years, etc., — is set by the user before the start of the simulation, with a trade-off between runtime and the accuracy of drought-response predictions. At the start of the simulation, the driest year among the first *n* years is selected as the year SurEau will be called; then, at the start of the *n* + 1 year, the driest year among the next *n* years is selected, and so on.

#### 2.2.3 Growth and mortality feedbacks

Once a SurEau simulation has been initialized from ForCEEPS, it proceeds to runs for one year like a standard SurEau-Ecos simulation, with the significant differences detailed in sections X and X. All through the simulation, and at the end of year, data is collected that is then sent back to ForCEEPS to determine the effects of drought stress on growth, mortality, and defoliation. The complete list of SurEau variables memorized and transferred is listed in Table X.

##### 2.2.3.1 Drought feedback on growth

In assessing the effects of drought events on trees, PHOREAU distinguishes between short-term adaptations and long-term non-reversible consequences — respectively feedbacks on growth and on mortality. The independence of these two mechanisms is vital to avoiding confusion between two sources of mortality: that caused by carbon starvation — represented in PHOREAU by diameter growth falling under a certain threshold — and that caused by hydraulic failure. A tree subjected to years of water stress may maintain its conductive vessels but die off due to a lack of carbon intake; another may die following a single month of acute water stress despite strong carbon reserves. By establishing a clear distinction between these two pathways, PHOREAU is able to account for the different drought response strategies observed among species.

In PHOREAU, the impact of water-stress on growth is assessed using a factor of stomatal closure. Compared to the original ForCEEPS formulation which computes a drought index *DrI* and a drought growth reduction factor *GR*_*drought*_ using a simple monthly water budget (Bugmann and Solomon, 2000), this new mechanism takes advantage of the detailed hydraulic framework of SurEAU to account for competition for water as well inter-specific differences in dealing with water-stress. While no longer used to control growth, the original drought index remains used as a proxy for global stand water availability when determining seedling establishment, and when ranking years according to overall dryness (see Section 3.4).

Schematically, as soil water reserves become depleted and soil water potential decreases, trees adapt their conductance by closing off stomata in order to reduce water loss and maintain twig and leaf potentials above cavitation thresholds (Cochard, Bréda and Granier, 1996; Cochard *et al*., 2002). This regulation mechanism prevents premature branch and tree death due to runaway embolisms, as trees reduce their water loss until only cuticular transpiration remains. The relation between leaf water potential and stomatal closure is an important trait describing a species’ response to drought: constrained by a trade-off between carbon gain and risk of hydraulic failure (Brodribb *et al*., 2003; Venturas *et al*., 2018), it is correlated with the more often measured turgor loss point (*TLP*) (Brodribb and Holbrook, 2003). While the link between turgor loss and reduced growth is well-documented and was considered for PHOREAU (Cabon *et al*., 2019; Peters *et al*., 2020; Potkay *et al*., 2022), in the end stomatal aperture was selected as a continuous variable allowing for a finer feedback.

Stomatal aperture in PHOREAU is derived at each time-step from leaf water potential — but also light, temperature, and CO2 concentration — using a sigmoid curve described by two species-specific traits: *P*_*gs*12_ the water potential causing 12% stomatal closure, and *P*_*gs*88_ the water potential causing 88% stomatal closure (Cochard *et al*., 2021b). Stomatal aperture in PHOREAU is defined as the ratio between actual stomatal conductance and a species-specific maximal value. To calculate the drought reduction index *DrI* of a given tree, daily stomatal apertures ratios *γ* are averaged over the photosynthetic period, which are then then averaged over the year.

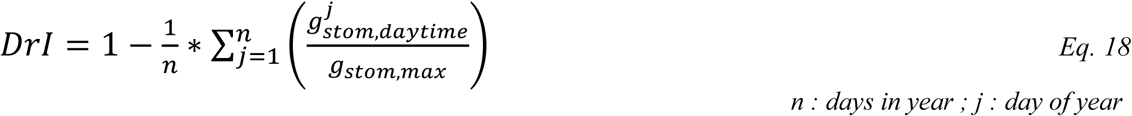

##### 2.2.3.2 Drought feedback on mortality

Drought-induced mortality in PHOREAU is derived from the percentage of cavitation, also known as the percentage of loss of conductance (PLC). This mortality mechanism is entirely distinct from the pre-existing slow-growth mortality in ForCEEPS, and the previously described drought feedback on growth. Indeed, contrary to the slow-growth mortality, which reflects carbon starvation and the long-term integrative effects of dehydration coupled with temperatures and competition for light on the capacity of trees to grow and survive (Bugmann and Solomon, 2000), this feedback is only intended to capture catastrophic water failure caused by extreme drought events, irrespective of the overall prior health of the tree. Unlike the stomatal closure used in drought feedback on growth, the cavitation of a tree’s hydraulic system is neither quickly reversible, nor does it follow a linear response to hydraulic stress. Furthermore, it occurs only after the stomata have been closed, when, under extreme stress conditions, residual water flow through the cuticle empties the plant’s water reservoirs. As water is drained from the soil and the water potential of the system becomes more and more negative, the conductance of a tree’s hydraulic system may remain stable until a certain point is reached, when it rapidly decreases as the xylem vessels are embolized and air are formed (Tyree and Sperry, 1989). This non-linear, tipping point response of conductance loss to decreasing water potentials is described by the *vulnerability curve* of the species. This curve, in the shape of an inverse sigmoid function, is described for each species using a *P*_50_ parameter. This parameter, responsible for the main differences in drought-resistance between species (Delzon and Cochard, 2014), is the water potential causing 50% cavitation in the xylem (Cochard *et al*., 2021b).

While the link between water potential and percentage of cavitation is deterministic for a given *P*_50_value, some additional steps are needed to translate this percentage into actual mortality in the model. First, the percentage of cavitation at the last time-step of the SurEau simulation is retrieved for each tree; t, as no cavitation-repair mechanism exists in SurEau (but see section 3.2.4), this will be always be the highest level of cavitation reached by the tree during the year. Second, the resulting percentage of cavitation is converted into a probability of death, which is applied at the end of the year like the other death probabilities in the model (Morin *et al*., 2021). In the event the tree aggregation option (see section 3.2.2.1) was used, each individual tree of a class receives the drought-induced death probability of its corresponding aggregate tree, and death events are drawn independently among them.

The actual conversion of PLC into a death-probability follows a logistic distribution fitted using data from Hammond *et al*., 2019. The probability distribution is parametrized using a constant steepness parameter, and a species-specific *LD*_50_parameter which corresponds of a point of no return, the lethal dose of cavitation at which exactly 50% of individuals of the species are expected to die (see Eq. 19). As a first approach this *LD*_50_ was parametrized at respectively 50% and 80% for gymnosperm and angiosperm species (Delzon and Cochard, 2014), reflecting the capacity of the latter species to operate at water potentials below the *P*_50_line. This is a result of differences in strategies between embolism-tolerant and embolism-avoidant species, as gymnosperms tend to operate at wider safety margins with vessels more resistant to embolism (Choat *et al*., 2012). Finally, an additional threshold parameter was added to avoid random mortality events for low PLC values, considering even well-watered trees show some degree of embolism throughout the year (Cruiziat, Cochard and Amiglio, 2002).

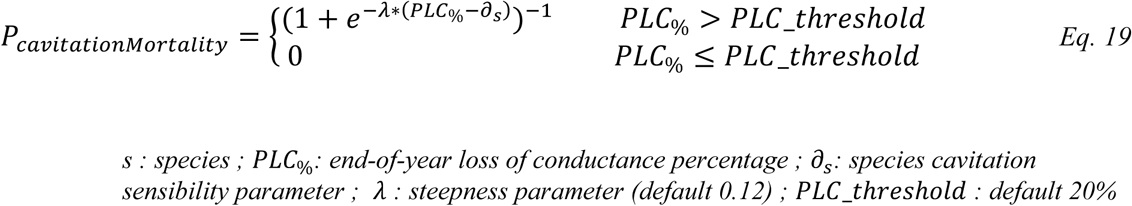

##### 2.2.3.3 Drought feedback on defoliation

Between the normal closing and opening of stomata to regulate water flow, and the runaway embolisms responsible for tree mortality after prolonged extreme droughts, trees exhibit a range of intermediate responses to water stress. Among these regulatory mechanisms, the adaptation of leaf area to moderate water stress is of particular importance for any model, such as PHOREAU, which integrates tree growth and drought-resistance.

Water limitation impacts leaf area through three main pathways: the premature shedding of leaves, the disruption of new bud formation (Bréda *et al*., 2006), and plastic biomass allocation to leaves (Martínez-Vilalta, Sala and Piñol, 2004). These mechanisms function at gradually longer time-frames: a cohort of trees may shed their leaves one year in response to extreme drought, and recover their full canopy the next; another may experience several years of decreased leaf area while its leaf phenology cycle is disturbed; and yet another cohort may have permanently shifted to produce less leaf area to adapt to chronic soil water limitations (Limousin *et al*., 2012; Martin-StPaul *et al*., 2013). This graduated temporal response is complicated by the fact it is differentially applied among species, following the classic split between drought-avoidance and drought-resistance strategies: indeed, there is evidence that while the reduction of leaf area improves resistance to moderate drought events, it may not avail against severe water stress (Limousin *et al*., 2022). Furthermore, the short-term gain in drought-resistance of a reduced photosynthetic surface may eventually offset by the negatives consequences of reduced carbon uptake (Poyatos *et al*., 2013), and the link between leaf area and a reduction of fine root biomass (Gieger and Thomas, 2002).

While the integration of defoliation has been shown to improve the predictions of tree mortality models (Dobbertin and Brang, 2001), this integration is complicated by the fact that few are able to account for the dual role of leaves in carbon-assimilation and water-use. However, unlike most mortality models, the PHOREAU model has the major advantage of being able to disentangle the contradictory effects of leaf area on growth and drought resistance, and of having an explicit representation of the root compartment with water uptake driven by fine roots and ultimately leaf area (see section 3.2.7).

From a mechanical point of a view, drought-induced defoliation in PHOREAU is derived on a daily basis for each tree from the percentage of cavitation of the leaf apoplasm. As such, the difference between drought-avoidant and drought-resistance species is already integrated in the PLC value. The resulting defoliation percentage is applied to the maximum leaf area of the tree for the given day, which is itself the result of species crown allometry, reduction of crown length due to competition for light, and the phenological stage of the leaf derived from PHENOFIT. As tree vessels begins to embolize and leaves to shed, the reduction in evaporative surface will naturally slow down the rate of embolism, as well as the water uptake as fine root area decrease. Furthermore, the list of daily drought-induced defoliation percentages are used in the light-competition module to determine light passage, and are integrated in the *GR*_*Crown*_ growth-reductor from crown length which represents leaf-loss impact on carbon assimilation (see Eq. 33). Finally, the longer-term adaptation between water stress and reduced leaf area is partially captured by the fact cavitation is carried over from year to year, with a specific repair mechanism described below. These relationships are summarized in Eq. 20.

#### 2.2.4 Year-to-year cavitation memory and repair

Unlike stomatal closure and adaptation of leaf area, the cavitation of xylem vessels is not a routine event in a tree’s life (Cochard and Delzon, 2013). The induced loss of conductivity can continue long after the end of the initial drought event, and is one of the main causes for the increased vulnerability to future drought events of previously weakened trees (Anderegg *et al*., 2013; Feng *et al*., 2021). On the other hand, internal repair mechanisms have been evidenced in many tree species, with a recovery of initial conductance over time. Because of the central role of PLC in the PHOREAU framework — driving both drought-induced mortality and defoliation— an accurate representation of its evolution over multiple-year periods was a necessary component of the model: as such, the end-of-year PLC value for a tree is carried over to the start of the next year, and it is on this basis that eventual repairs are made.

While trees have the ability of recover lost hydraulic conductance, the mechanism of this recovery does not seem to involve the actual reparation of embolized xylem vessels, but rather the formation of new vessels (Brodribb *et al*., 2010). As such, the recovery from cavitation is driven by basal area growth — or, more precisely, by the relative increase of sapwood area, which contains the living conductive vessels. While all new growth is naturally sapwood, as a tree becomes larger the relative proportion of sapwood to heartwood tends to decreases; it follows that to evaluate the rate of replacement of the conductive vessels, the model must first know the pre-existing area of sapwood. PHOREAU uses the foliage area to determine this quantity, through the application of a species-specific, constant, leaf-to-sapwood ratio, also known as the inverse of the Huber value (Cruiziat, Cochard and Amiglio, 2002). The leaf-to-sapwood ratio is applied to the potential one-sided leaf area of the tree, derived solely from its DBH and allometry parameters, and not its actual leaf area after defoliation through competition, frost or drought. This approach, presented in Eq. 20, assumes the Huber ratio to be constant: we know that this is in fact an important simplification, and that many species adapt their leaf mass per area to site conditions (Lopez *et al*., 2008).

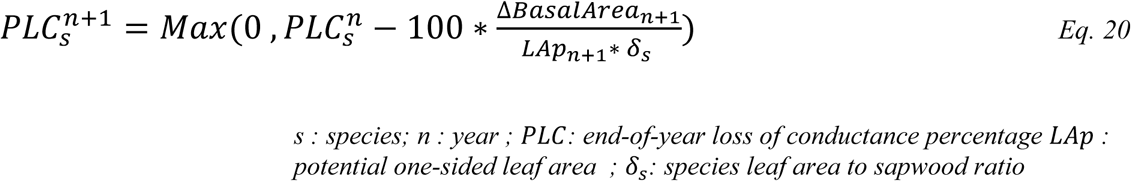

#### 2.2.5 Leaf phenology integration

PHOREAU takes advantage of the leaf phenology submodule in PHENOFIT to inform the evolution of leaf area throughout the year. Using the dates of leaf budding and leaf senescence calculated for a given species for a given year, PHOREAU gradually increments (respectively decrements) tree leaf area over an interval of days starting from the budding date (respectively senescence date), using two species-specific parameters for the length of leaf unfolding and coloration. See Eq. 21 for more details.

This formulation has the disadvantage of disregarding differences in phenology arising from differences in size or competition-status. Furthermore, it does not take full advantage of the PHOREAU hydraulic submodule to account for the effects of drought on leaf development, either through earlier leaf coloration (Xie *et al*., 2018) or shifted unfolding (Cleland *et al*., 2007). Further developments of the PHOREAU model should strive to use information from the light competition and water stress modules to inform the calculation of phenology dates.

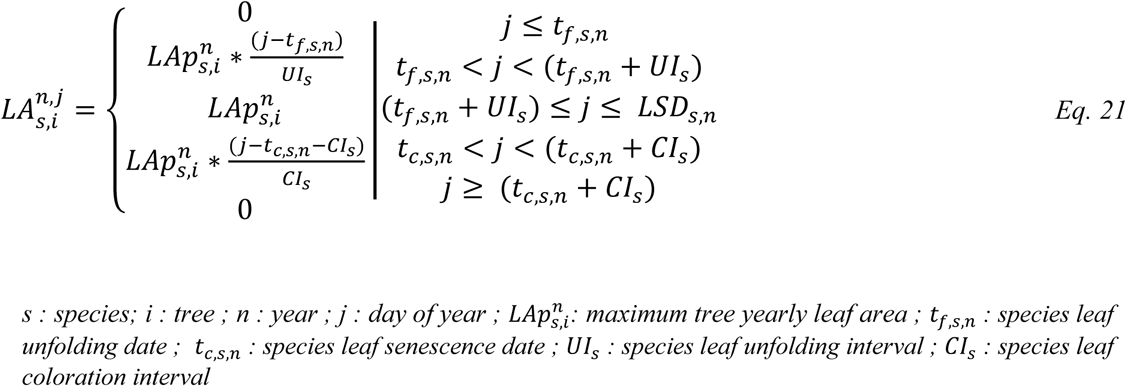

#### 2.2.6 The rain interception module

Capitalizing on the capacity of PHOREAU to predict individual-tree daily foliage area values that integrate allometry, competition, frost, phenology, and drought-defoliation effects, we implement a rain interception module that reduces incoming rain based on the daily leaf area of the stand. Modelling rainfall interception — defined as free water that evaporates from the leaves and barks of trees after a rain event — is an important component for any model trying to water cycles and tree water balance (Granier *et al*., 1999; Davi *et al*., 2005a). The intensity of the interception has been shown to grow linearly with leaf area, for values ranging from 20% to 35% of cumulated rainfall in temperate and continental climates (Bréda *et al*., 2006). While secondary factors such as irradiance, windspeed, and vapor pressure deficit impact the rate of interception *in natura*, as a first approach we have chosen a simple implementation, inspired from Medfate (De Cáceres *et al*., 2023), based solely on daily leaf area, rain volume, and potential evapotranspiration.

A canopy storage volume is derived from the foliage area of the stand. This volume is incremented at a daily time-step with incoming rainfall, and outgoing evaporated water. For a given volume of incoming rainfall, the throughfall, or the volume of water to reach the ground, is calculated with a simplified *Beer-Lambert* formula, in a similar fashion to the way light extinction is computed. Because the canopy storage volume is itself limited, any intercepted water that overflows this maximal quantity flows down the soil; a natural consequence of this property is that a given volume of given rainfall will yield a greater cumulated throughfall when concentrated in a single day, than when distributed over several days with intervening evaporation. The algorithm, presented below in Eq. 22, computes the daily stand-wide throughfall volumes that then serve as inputs to the water balance model.

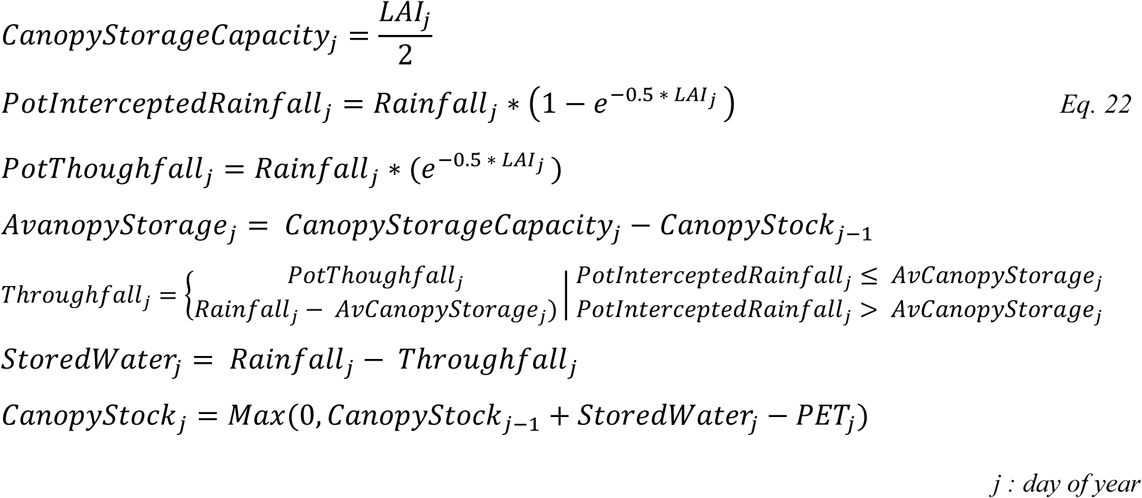

#### 2.2.7 The soil compartment in PHOREAU

The representation of root processes is vital for any model aiming to simulate the response of vegetation to climate change (Woodward and Osborne, 2000). Because of this, the framework for representing roots in PHOREAU had to be considerably expanded compared to the parent model where the rooting system was reduced to a simple fine root surface. In particular, we built upon the SurEau-Ecos framework by integrating coarse root depth alongside fine root surface, having the roots of different trees share the same soil to compete for water, and implementing plastic responses of root biomass and root depth to drought stress and aboveground growth.

The modelling of the root compartment in PHOREAU is based on the same major hypothesis as that of the canopy and light competition module: an implicit homogenous horizontal distribution of trees, with an explicit vertical stratification. In the same way the aggregated vertical distribution of foliage area entirely determines the light availability of each tree, with no accounting for its location within the stand, competition for soil water between trees in PHOREAU is a direct result of the vertical distribution of their root systems. The underlying hypothesis is that all trees compete for the same water reserves, provided their roots go deep enough; and the modeler must take care to select a simulation stand area that verifies this constraint, which will itself vary according to the size and rooting structure of the trees present in the stand.

In PHOREAU the rooting system of a tree is split between fine roots and coarse roots: this distinction is essential as the roots types have different functional roles and responses to external factors (Pregitzer, 2002). *In natura,* coarse roots (usually > 2mm in diameter) are responsible for anchoring the tree, exploring the soil vertically, and transporting water and nutrients from deeper horizons; meanwhile fine roots are responsible for nutrient and water uptake, and have been found to extend laterally more than 20 m from the tree trunk. While roots are estimated to represent 20-40% of tree biomass, fine roots make up a small fraction of this quantity; however, they play an important role in carbon sequestration due to their high turnover rate. (Jackson, Mooney and Schulze, 1997). Therefore, in addition to providing a more accurate representation of tree response to drought stress, this explicit representation of the rooting system will serve as a necessary first step to the simulation of a tree’s carbon budget.

Schematically, fine roots extend horizontally to absorb water in the available soil, while coarse roots explore deeper layers and make them available to fine root exploration. In PHOREAU, where the soil is segmented in a number of layers, this has been translated in the following way: the fine root area of a tree in a determines the conductance between this tree and a given soil layer, while the rooting depth determines which layers the tree has access to, and how its fine root area is distributed within them.

In practice this means that, for a given set of soil parameters, certain trees will be able to extract water from the full soil profile, while others will be restricted to only a fraction. This can be seen in Figure 7, extracted from the PHOREAU validation on the ICOS sites. This framework is intended to reflect the crucial role of rooting depth in resilience to drought stress (Canadell *et al*., 1996), as trees with deeper rooting systems are able to make use of relatively untouched water reserves in deeper soil layers. Furthermore, because this is implemented in a forest dynamics model where many trees share the same soil, PHOREAU will be able to use the differential rooting depths to explore the contrasting intra and inter-specific drought responses observed in nature (Johnson *et al*., 2018).

Rooting depth is a notoriously difficult trait to measure, and involves costly, time-consuming, usually destructive techniques (Maeght, Rewald and Pierret, 2013). While some rooting depth data is available in the literature (Guerrero-Ramírez *et al*., 2021), its scarcity makes it difficult to disentangle environmental, allometric, and genetic factors; what is driven by aboveground biomass, from what is driven by water availability and groundwater table depth (Fan *et al*., 2017; Freschet *et al*., 2021; Li *et al*., 2022). To circumvent this difficulty in obtaining accurate rooting depth traits, we take advantage of the fact PHOREAU does not explicitly represent the position of a tree in the plot and ignores lateral distribution, by using coarse root biomass — an extensively studied trait — as a proxy for rooting depth, thereby implicitly aggregating the lateral and vertical extension of the root system in an integrative rooting extent variable, which is driven by shoot size and site aridity (Tumber-Dávila *et al*., 2022).

Coarse root biomass and fine root biomass in PHOREAU are calculated independently. Fine root area is derived on a 1:1 basis from leaf area. Meanwhile, coarse root biomass is calculated as the product to above-ground biomass with a root-shoot ratio, this root-shoot itself calculated as ratio of realized tree height to maximum species height, positively modulated by the mean of past drought indices (Morin *et al*., 2021). This formulation, shown in Eq. (23), follows the conclusions of Ledo *et al*., 2018 which identifies size and past droughts as the main factors driving root-shoot. These simple equations allow PHOREAU to capture several well-established characteristics of the evolution of coarse and fine root biomass.

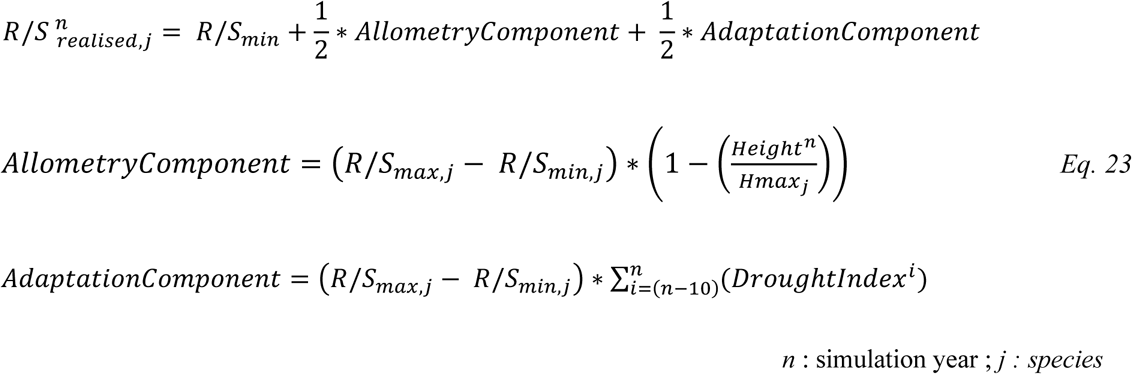

Similarly to leaf shedding, fine root area tends to decrease as an avoidance mechanism following past drought events (Hartmann, 2011; Brunner *et al*., 2015). Meanwhile, total root biomass relative to aboveground biomass (the *root-shoot ratio)* has repeatedly been shown to be positively correlated to past drought events (Mokany, Raison and Prokushkin, 2006), and tree species adapted to more xeric climates have higher root-shoot ratios and deeper roots than those adapted to wetter conditions. These results are in accordance with Optimal Resource Partitioning theory (OPT), which predicts should trees will increase their absorptive capacity relative to their transpiring surface under short water supply (Coomes and Grubb, 2000; Hertel *et al*., 2013).

Another observation captured by deriving root biomass from relative height in PHOREAU is the negative correlation between root-shoot ratio and above-ground biomass (Mokany, Raison and Prokushkin, 2006; Ledo *et al*., 2018). Because tree height in PHOREAU tends asymptotically towards the species’ maximum height following a parabolic curve, as trees become older they allocate proportionally more growth to their diameter than to their height — and to their roots in the new formulation. Following Konôpka *et al*., 2010, the maximum root-shoot was set to be greater for angiosperms than coniferous trees, who tend to have shallower roots (Schenk and Jackson, 2002) and less variation between juvenile and adult individuals. Another implication of this formulation is that the proportion of fine roots exponentially decreases with total root biomass (Li *et al*., 2003). - An emergent property of this framework is that for a given magnitude of water stress, a site which has already suffered past drought events will suffer less mortality and growth loss than a previously wet site, because of the rooting depth adaptation mechanism (Fuchs *et al*., 2020). This type of plastic adaptation is crucial to finely assess drought vulnerability at a regional level, where for the same magnitude of climate change dieback has been shown to vary among stands of similar composition depending on the level of past drought acclimatization (Piedallu *et al*., 2023). Figure 8 shows an example of this emergent behavior, by comparing simulations with two different climatic trajectories.

This integration of root plasticity, coupled with leaf shedding, is an important first step in the modelisation of tree adaptation to drought conditions. However, it by no means provides a complete picture of the various strategies used by trees *in natura.* To refine our approach the relative importance of past drought conditions relative to that of tree allometry in determining total rooting depth could be determined on a species by species basis, instead of a simple angiosperm/coniferous split. Furthermore, the magnitude of past drought conditions is currently assessed on a patch-level instead of a tree level-basis, ignoring the role of factors such as species and size in determining the actual drought stress suffered by a tree for a given set of climatic and edaphic conditions. Even then, root plasticity is only one among many plastic responses to drought conditions: regulatory responses have been identified in the ectomycorrhizal network, non-structural carbohydrate concentration, differential gene transcription and pathways, increased suberin and lignin formation in roots, and decreased fine-root turnover rate (Bréda *et al*., 2006; Brunner *et al*., 2015).

### 2.3 Integrating phenology-based reproduction

The PHENOFIT model (Chuine & Beaubien 2001), a process-based species distribution model for temperate trees, outputs the probability of presence over several years of a given species for a particular set of environmental conditions. This probability is derived from the estimated fitness of an average adult individual of that species, which is itself the product of the probability to survive until the next reproductive season, and the probability to produce viable seeds by the end of the annual cycle. The model assumes that survival and reproduction depend on the synchronization of tree development to seasonal climatic variations, with the plasticity of key phenological events such as leaf unfolding, flowering, fruit maturation, and leaf senescence. These are the result of different sub-models, using only soil data and daily meteorological data (minimum and maximum temperature, rainfall, relative humidity, global radiation, and wind speed) as inputs: these include phenological models for leafing, flowering, fruiting (Morin, Augspurger and Chuine, 2007) for leaf senescence (Delpierre et al., 2009); a frost injury model (Leinonen, 1996); a survival model; and a reproductive success model calculated as the proportion of uninjured fruits that reach maturation considering photosynthetic ability and the proportion of leaves not killed by frost (Chuine and Beaubien, 2001). A visual representation of the model can be found in Figure 1.

In PHENOFIT, the determination of leafing and flowering dates (*t*_*f*_) both use the same two-step system. First dormancy (Eq. 24), whereby the bud must undergo a chilling period, accumulating a certain quantity (*C*_*c*_) of chilling units (*R*_*c*,*t*_) from the onset of dormancy (*t*_0_) until dormancy is broken at a date *t*_1_. Secondly quiescence (Eq. 26), where the organ accumulates forcing units (*R*_*f*,*t*_) until a threshold value (*F*_*c*_) is reached, and budburst is completed. Only the parametrization and the shape of equations for the calculation of forcing and chilling unit change between leafing and flowering: they are calculated at daily time-step, using mean daily temperatures (*T*_*t*_) and species-specific parameters (*a*, *b*, *c*, *d*, *e*) as shown in Eq. 25 and Eq. 27. Leaf senescence dates *t*_*c*_ are calculated independently, following Delpierre et al., (2009).

Flowering and leafing dates are then used, alongside the daily minimum temperature after budburst (*T*_*i*_), to determine proportions of uninjured leaves and flowers (*I*_*l*_,*I*_*f*_). The probability that fruits will reach maturation (*I*_*r*_) is then calculated on the basis of the proportion of uninjured leaves which produce the assimilates accumulated in the fruits, the date of flowering from which thermal energy can begin to be accumulated, and a fitted species-specific parameter *E*_*c*_ representing the average amount of energy needed to reach maturation (Eq. 30). Finally, a yearly probability of producing viable seeds, or reproductive success (*R*), is calculated as the product of the probability that fruits will ripen and the proportion of uninjured flowers, as shown in Eq. 31.

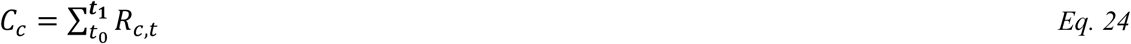

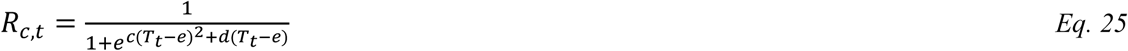

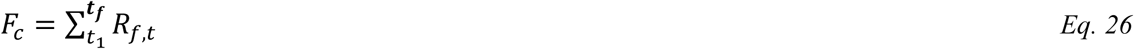

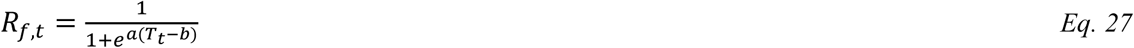

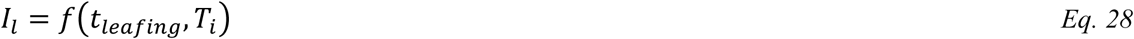

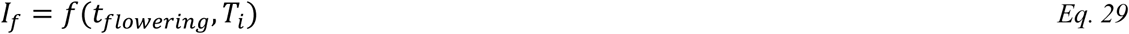

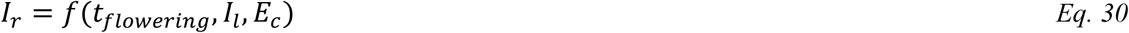

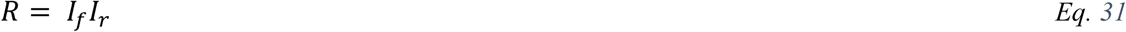

For each organ and each species, parameters are fitted with phenological observations from native populations (dates of leaf unfolding, senescence, flowering, and fruiting) or from experimental results found in the literature (resistance of plant organs to frost), and not on present species’ distribution. Reaction norms to temperature, photoperiod and water availability are inferred statistically using time series data from different sites and different years.

As the model only simulates one average individual, it does not take into account demography or biotic interactions with other species. It also does not represent the impacts of plant growth on survival and resource allocation. While it can represent the way phenological plasticity can vary from one site to another by calibrating parameters from phenological data of different provenances, we have chosen here to use only one calibration set per species: in other words, we account for the plastic response of a species to varying climate conditions, but not for the genetic differentiation of this response. As a result, adverse adaptations may be over-estimated: for example, a species’ population at higher latitude than the one used for calibrations may be simulated as flowering too late, due to higher energy requirements than in reality.

The version of the model used in the study, as well as each species’ response parameters, are distributed on the Capsis platform (Dufour-Kowalski et al., 2012). A description of the model can be found in (Chuine and Beaubien, 2001).

### 2.4 Updated model equations

Our main concern when integrating PHENOFIT and SurEau with the ForCEEPS model was avoiding overlaps, and especially a plant being penalized twice for the same factor. This meant that we could not directly use the *PHENOFIT* global plant fitness output. Similarly, we did not use the plant survival output, as lethal frost is minimal for our temperate range, and drought-caused mortality will be accounted for by the SurEau model. In the end, we used four main yearly outputs: leaf unfolding and senescence dates (*t*_*f*_, *t*_*c*_), the percentage of uninjured leaves not damaged by frost (*I*_*l*_) and reproductive success (*R*)

*Leaf phenology,* in particular leaf unfolding and leaf senescence dates, were used to control plant fluxes (see Section 3.2) and the range during which growing degree days (GDD) are accumulated for deciduous species. Evergreen species are still assumed to accumulate energy throughout the year. As the original ForCEEPS framework worked at a monthly time-step, it was necessary to update the model to calculate GDD using daily temperature data. This introduces both inter-species variability in growth, but also intra-species variability between sites and years. This change impacts both growth (through the temperature growth-reduction factor *GR*_*gdd*_) and probability of establishment (*P*_*GDD*_). See Eq. 32 for the updated calculation of annual GDD sums, including phenology and microclimate, of a tree of species *s* and average weighted foliage height *h*, with *T*_0_ the base temperature (*T*_0_ = 5.5°*c*).

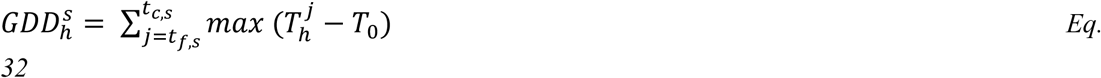

*Climate-induced leaf loss* was integrated into ForCEEPS as a modification to the previously existing crown length reduction factor *GR*_*crownLength*_, which represents the impact of leaf density on growth through carbon assimilation. While in the previous ForCEEPS framework trees could only lose leaves through a lack of light availability (the *lightcomponent*, presented in Morin et al. 2021), PHOREAU also captures drought-induced and frost-damage leaf loss, which are integrated in the updated calculation for *GR*_*crown*_ as shown in Eq. 33, Eq. 34 and Eq. 35. This is a first approach, following Wang, Zhou and Wang, (2021). We are aware this representation is incomplete, and does not account for leaf regrowth, or differential effects according to tree age and size: the absence of an explicit representation of source and sink compartments, and the lack of tree age data to implement an age-differentiated response to leaf loss, was a limiting factor.

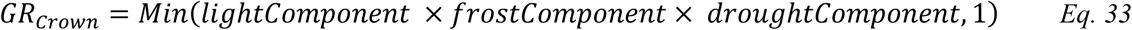

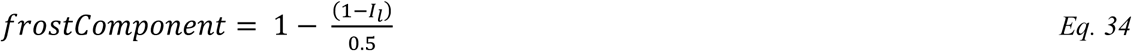

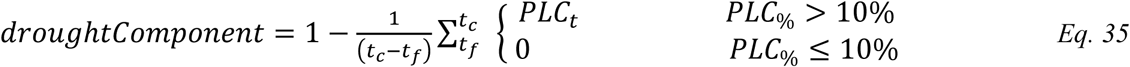

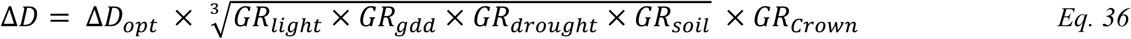

*Phenofit reproductive success,* calculated as the product of the proportion of injured flowers and the proportion of fruits that reach maturity, was used to update the different ForCEEPS regeneration modules. In particular, the yearly number of potential seedlings for each given species is multiplied by its yearly reproductive success *R*_*s*_. Once the number of potential seedlings for a given species has been determined as shown in Eq. 37, the probability of establishment of each individual seedling *P*_*est*,*s*_ is unchanged from the original ForCEEPS framework (Eq. 38, with details in Morin *et al*., 2021), but takes into account the underlying refinements in the calculation of phenology, microclimate, light availability at soil level, and soil water balance.

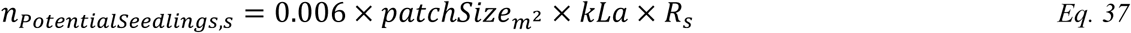

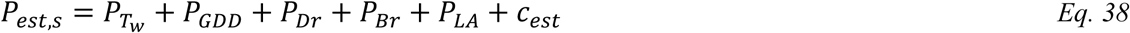

*Tree mortality* was simply updated to reflect the new cavitation mortality mechanism described in Eq. 39. With *P*_0_ and *P*_*g*_ respectively the background and growth-related mortality components described in Morin *et al*., (2021), the chance that a given tree dies on a given year is such that:

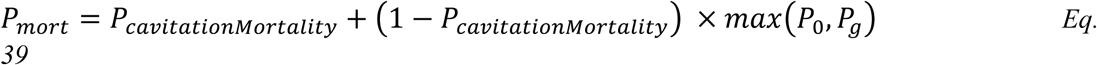

## 3 Data sources and model validation

In this study, we present a novel validation approach that evaluates our model across a broad spectrum of outputs, ranging from daily plant physiological measurements to long-term species composition predictions. This comprehensive approach allows us to avoid one of the common pitfalls of gap-model, which are often validated on a single integrative metric — such as predicted total stand basal area — and are thus limited in making robust predictions under future conditions. By directly assessing the model’s ability to reproduce intermediary variables, such as leaf area indices or soil water fluxes, we can control for common biases that may arise from errors offsetting each other under current conditions, which may not hold true when projecting into future climatic scenarios.

Because PHOREAU is intended to be continuously improved and refined over time, the validation protocol and all associated data — summarized in Figure 9 — will serve as a baseline to evaluate any future modifications to the model.

**Figure 9.**
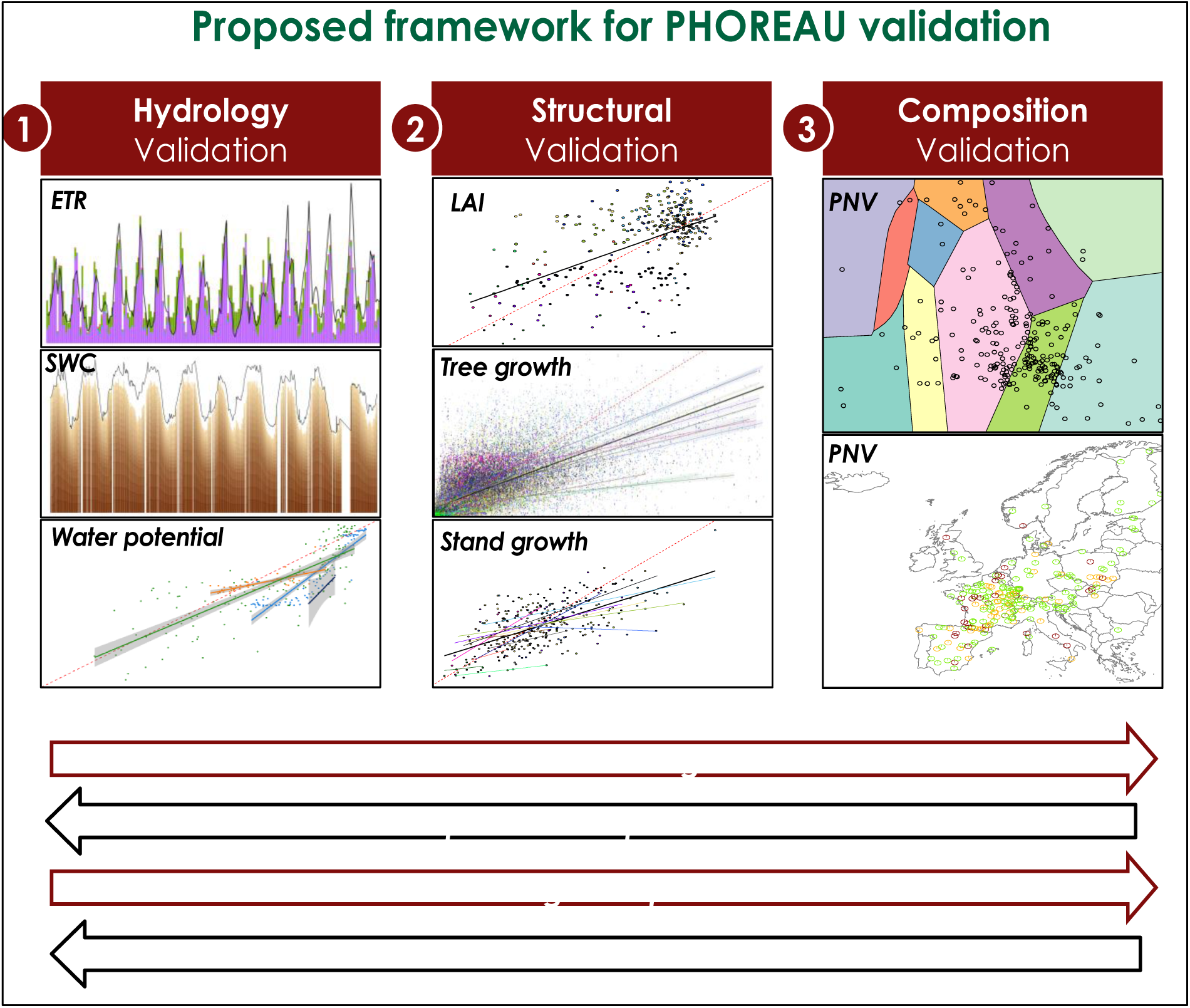
Proposed framework for PHOREAU validation. All graphs are snapshots of model results presented in this study: Real evapotranspiration (ETR); Soil water quantity (ETR); stem water potentials; leaf area index (LAI); tree and stand productivity; and potential natural vegetations (PNV).

### 3.1 Climate and soil data

Compared to the original ForCEEPS framework, the PHOREAU model requires additional climatic and soil measurements (see Table 1) at a daily instead of monthly time-step. To validate PHOREAU we used the ERA-5 Land dataset, which is a refinement of the ERA5 climate reanalysis dataset at a finer spatial resolution (Muñoz-Sabater *et al*., 2021). The hourly data was aggregated to produce daily timeseries from 1969 to 2021 at a 0.1×0.1° resolution, or roughly 9km, for all of Europe. Potential evapotranspiration were then calculated at the same resolution using the Penman Monteith equation (Monteith, 1965).

**Table 1.**
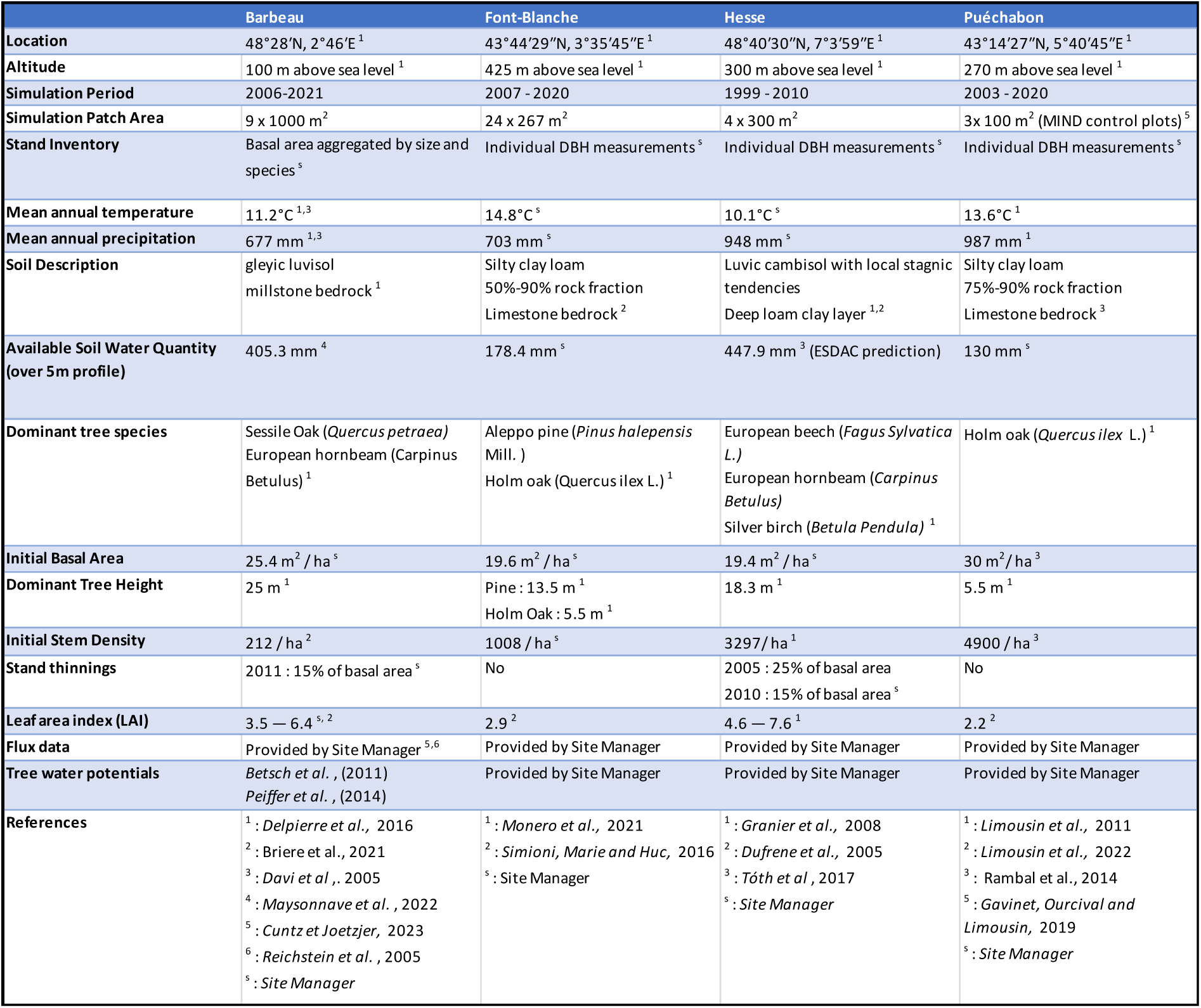
Selected stand characteristics for the four ICOS sites used in the in-depth PHOREAU validation, with associated data sources.

Instead of a single soil site water holding capacity (SWHC), PHOREAU requires, for each layer of soil (in this study 30 layers, up to a total depth of 5m), the fraction of coarse elements, as well as the parameters of the Van Genuchten water retention curve which describes the soil texture (Van Genuchten, 1980). These parameters were obtained for several depths from the European Soil Hydraulic Database (ESDAC), and interpolated over the height of the soil profile.

The resulting ESDAC soil and ERA-5-Land climate parameter files were used as a baseline for our European validation, and were directly used for the ICP II plots, for which no other climatic or soil data was available. When possible, we completed this continental-scale data with higher-resolution measurements. On-site measurements were naturally available for all four ICOS sites; for the RENECOFOR plots, we used a combination of soil measurements and the SILVAE climate time-series to refine our validation.

The SILVAE web portal (Bertrand *et al*., 2011 and Richard, 2011) offers monthly average temperature and precipitation sum data over France at a finer spatial resolution, accounting for microclimatic differences caused by differences in altitude, exposition, and wind orientation. These time-series, available for the period between 2000 and 2014, were used to correct the coarser ERA-5 Land dataset for all variables except wind-speed: either by direct mean-adjustment for the average temperature and precipitation variables, or after a prior linear regression of the variable over the mean temperature for the given month of the ERA-5 Land time-series. For the average temperature variable, between 2000 and 2014, daily values were corrected by the difference between the average of all the daily ERA-5 Land values for that month and the single monthly value of the SILVAE correction dataset; whereas for the years outside of this range where the corresponding monthly value was not available, the difference was calculated using the mean of all values for the given month between 2000 and 2014. The same method was used for the precipitation variable, except the daily values were multiplied by a ratio instead of impacted by a difference. For the other variables except wind speed the same method was used as for the average daily temperature, except the addition factor was itself first multiplied by the slope of the regression between the temperature and the variable (see Table S1 for slope and R^2^ values). The wind speed variable was not corrected due to its weak correlation to mean temperature. The workflow for climate reconstruction is summarized in Figure **10**10.

**Figure 10.**
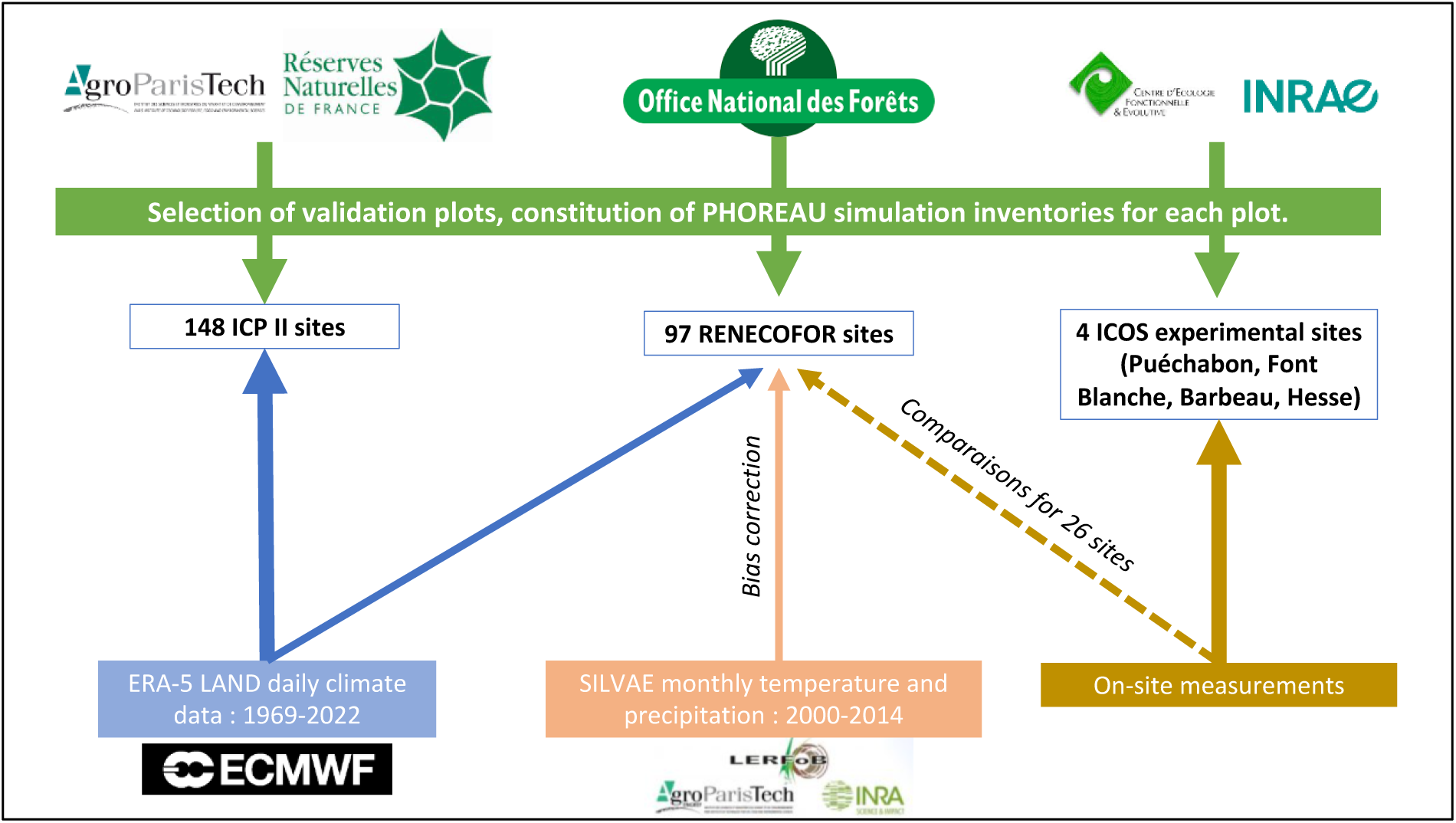
Summary of the workflow used for constructing ForCEEPS inventories and climate datasets.

Local measurements of SWHC were available up to a depth of 1 meter for all RENECOFOR plots (Brethes and Frankreich, 1997). Additional measurements were available up to 2 meters for more than half of the plots (Brethes and Frankreich, 1997; Lebourgeois, 2006; Guillemot, unpublished data), which were used to refine validation soil parameters.

On-site climate measurements were available for 26 of the 102 RENECOFOR sites (see Table S4 for the list of sites). For some of the sites the measurements periods only partially matched the simulation periods, while for others they were continuous from 2000 to 2021. For the RENECOFOR plots these measurements were not used in the main validation protocol, so as to keep a homogenous climate construction method that allowed comparisons between sites. They were however used to validate our climate reconstruction: first through direct comparisons of climate variable means and variances, and then by comparing the outputs of the ForCEEPS simulation between on-site and reconstructed climatic data (refer to Table S13).

### 3.2 ICOS sites

We used data from the Integrated Carbon Observation System (ICOS) for our most in-depth validation protocol that includes hydrological, growth, and mortality components. In particular, we selected four forested sites from the terrestrial ICOS Ecosystem: Puéchabon, Font Blanche, Hesse, and Barbeau. Together these sites represent a diversity of the climatic, edaphic, and biotic conditions that can be found in France.

The Integrated Carbon Observation System is a network of stations that use the eddy-covariance technique to measure ecosystem-atmosphere exchanges of greenhouse gases and energy at a high frequency (Baldocchi, 2003). In addition, a large set of ancillary variables needed for the interpretation of the flux data are also measured: for forest stations these include but are not limited to tree inventories, leaf area index, and soil data — all of which can be leveraged for our modelling purposes (Gielen *et al*., 2018). The large scope of measured variables in ICOS framework makes any validation based on it easily scalable, and will in the future allow testing of newly integrated PHOREAU processes such as carbon retention or vertical micro-climate interpolation. Finally, a set of rigorous specifications for the installation of the eddy-covariance tower sensors, and a common pipeline for the post-processing of the raw data through the Ecosystem Thematic Centre (ETC), ensure the high level of comparability between sites that is necessary for large-scale model validation. Eventual gaps in data were corrected by selecting, for each of our four sites, the simulation period where the most harmonized data was available.

A preliminary task was building an exhaustive database of all relevant input and output variables over the selected sites. This was made possible by the collaboration of each of the site managers, especially for non-flux data that was not always readily available on the ETC database (Reichstein *et al*., 2005; Papale *et al*., 2006). Table 1 provides a summary of the data sources used in the ICOS validation, as well as some of the main site characteristics. Figure 11 shows a simulated representation of the initial state of each inventory, highlighting the structural diversity across sites.

**Fig 11.**
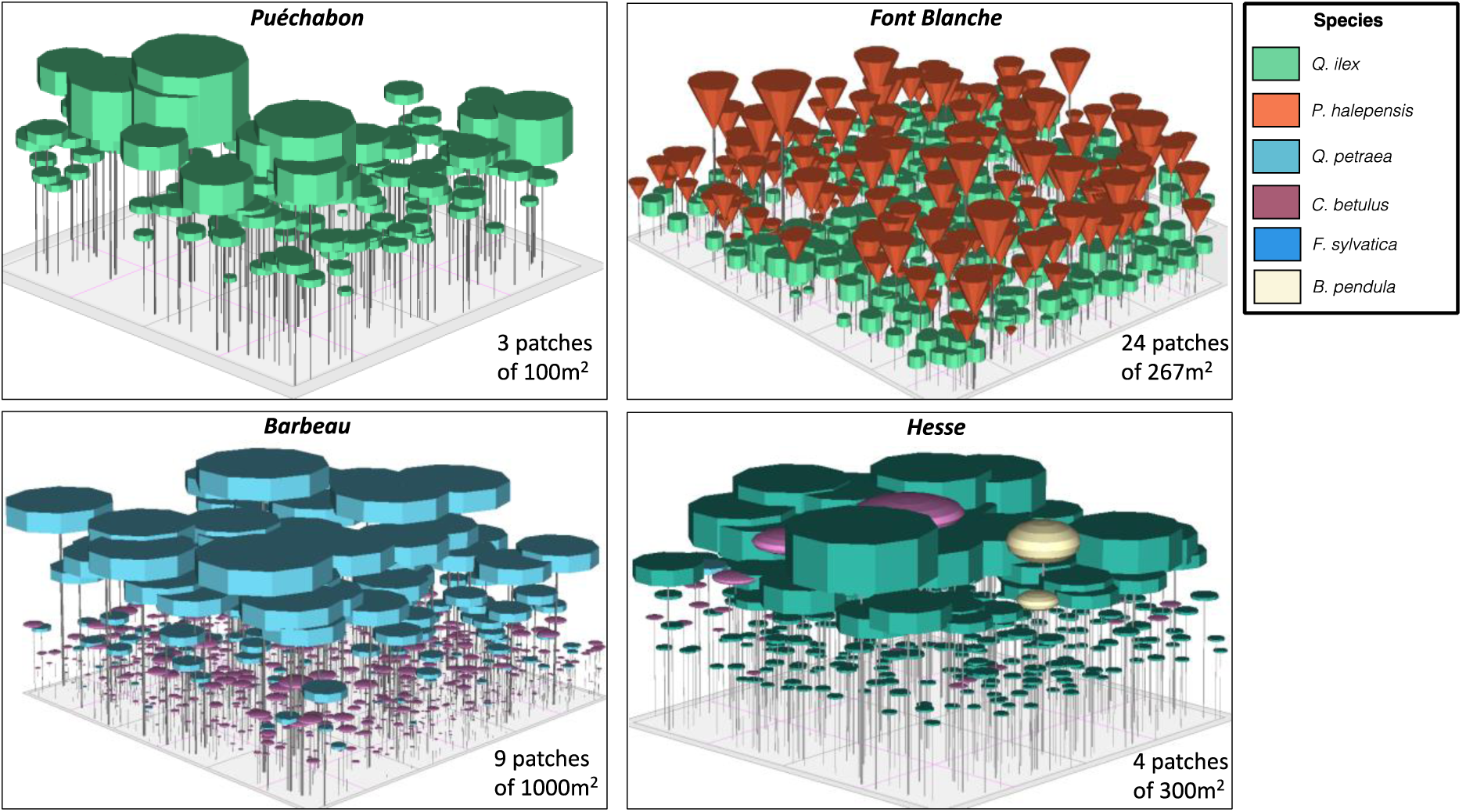
3D vizualisation of ICOS stands used for indepth validation. Vizualisation generated by the PHOREAU model, on the basis of initial inventories.

#### 3.2.1 Puéchabon

The Puéchabon experimental site (43°44’30”N, 3°35’40”E, altitude 270 m) is located in a forest of holm oak located in the South of France near Montpellier. With its last clear cut in 1942, and managed as a coppice for centuries before that, the site is characterized by a high density (5000-7000 trees/ha) of small (5.5 meter high overstorey) *Quercus ilex* trees: they make up an old forest with a basal area of 30 m^2^/ha, (Rambal *et al*., 2014), and an LAI around 2.2 with little seasonal variability. Located on a flat area, with a rocky soil of Jurassic limestone filled with clay, its small water reserve (roughly 130 mm of water over the 5 meter profile) and typically Mediterranean precipitation pattern (highly variable from year-to-year, with a measured range of 550 to 1550 mm primarily concentrated between September and April) made it an ideal candidate to study the long-term effects of drought.

Within the framework of the *Mediterranean Terrestrial Ecosystems and Increasing Drought* (MIND) project, the diameter of trees contained in twelve 100m^2^ plots have been measured on a year-to-year basis since 2003: these are distributed between three control plots, three thinned plots (33% reduction of basal area), three plots with partial rainfall exclusion (33% throughfall), and three thinned and rainfall excluded plots (Gavinet, Ourcival and Limousin, 2019). We have used these plots to run simulations from 2003 to 2020, and assess how the PHOREAU model simulates the effects of tree density on drought resistance.

#### 3.2.2 Font Blanche

The Font Blanche experimental site (5°40’45’’E, 43°14’27’’N, altitude: 420 m) is located in a mixed-forest of Aleppo pine and holm oak, with an overstorey of *Pinus halepensis* (13.5 m height) that dominates a coppice of *Quercus ilex* (6.5 m height). With a basal area of 21.3 m^2/^ha and and LAI ranging between 2.5 and 2.7 it is less dense than Puéchabon, but otherwise boasts a broadly similar soil and meteorological profile (Simioni, Marie and Huc, 2016). For our validation we used the 625m^2^ control plot (PM30) of the rainfall exclusion experiment, in addition to the main plot of 6400m^2^ that we split between 25 smaller splots of 267 m^2^ apiece to satisfy PHOREAU homogenous competition assumptions. Our timeframe for this site ranges from 2007 to 2020.

#### 3.2.3 Hesse

The Hesse Experimental site (7°3’59’’E, 48°40’30’’N, altitude: 300 m) is located in a beech (*Fagus sylvatica*) stand in north-eastern France, on a plain at the feet of the Vosges mountains. Average tree height was measured at 16.2 m in 2005, with a maximum leaf area index over 7.5 indicating a very high level of canopy closure. In comparison to the two previous sites it is characterized by a wetter, semi-continental temperate climate, with a deep loam-clay soil (Davi *et al*., 2005a; Dufrene *et al*., 2005). Unlike most sites in the ICOS network it is fertile, fast-growing and subjected to frequent thinnings, with an average tree age of only roughly 40 years in 2005, allowing us to test the capability of PHOREAU to simulate canopy and basal area regrowth after a cut. Furthermore, despite the stand having high rainfall and soil high water holding capacity, droughts events are responsible for most of the interannual variability in tree growth (Granier *et al*., 2008). We extracted from the inventory four evenly sized 300 m^2^ plots. Because the validation timeframe ranges from 1999 to 2010 when the most data was available (Cuntz *et al*., 2023e, 2023d, 2023c, 2023b, 2023a). (Betsch *et al*., 2011; Peiffer *et al*., 2014; Tuzet *et al*., 2017; Zapater, 2018), our model also replicates two thinnings that occurred in 2004 and 2009, respectively for 25 and 15% of the basal area.

#### 3.2.4 Barbeau

The Barbeau experimental site (2°46’E, 48°28’N, altitude: 100 m) is located in the Fontainebleau national forest southwest of Paris. The stand is dominated by sessile oak (*Quercus petraea*) trees that 25 m at 100 years of age, with an understory of hornbeam (*Carpinus betulus*). Mean annual cumulated precipitations of 677 mm are evenly distributed over the year, and feed into a deep soil with roots able to reach at least 150 cm in depth. We initialized our validation over 9 plots of 1000 m^2^ using an exhaustive inventory made in the winter of 2006-2007; we ran running it until 2021, including a thinning in 2011 (Delpierre *et al*., 2016; Maysonnave *et al*., 2022). Unlike the other studied sites, growth data was not available on a tree by tree basis, but instead aggregated at the stand level (Briere *et al*., 2021).

### 3.3 RENECOFOR and ICP II sites

To validate our model’s prediction of simulated tree and stand productivity against real forest-growth datasets, as well as model performance in reproducing potential natural vegetations and observed foliage areas, we used 245 plots spread across Europe, from 37.03° N to 69.58° N, and 8.17° W to 30.71° E, covering most of the major European species (Figure 12. They cover a large range of environmental conditions, with mean annual temperatures (MAT) ranging from –1.62 to 17.6 °C, mean annual precipitation sum (MAP) ranging between 405 and 2707 mm, growing degree days (GDD) ranging from 475 to 4287 °C, and available water quantities ranging from 30 to 671 mm over the soil profile. Refer to Figure 19 for the distribution of site abiotic conditions, and Table S2 for a detailed site by site breakdown.

**Figure 12.**
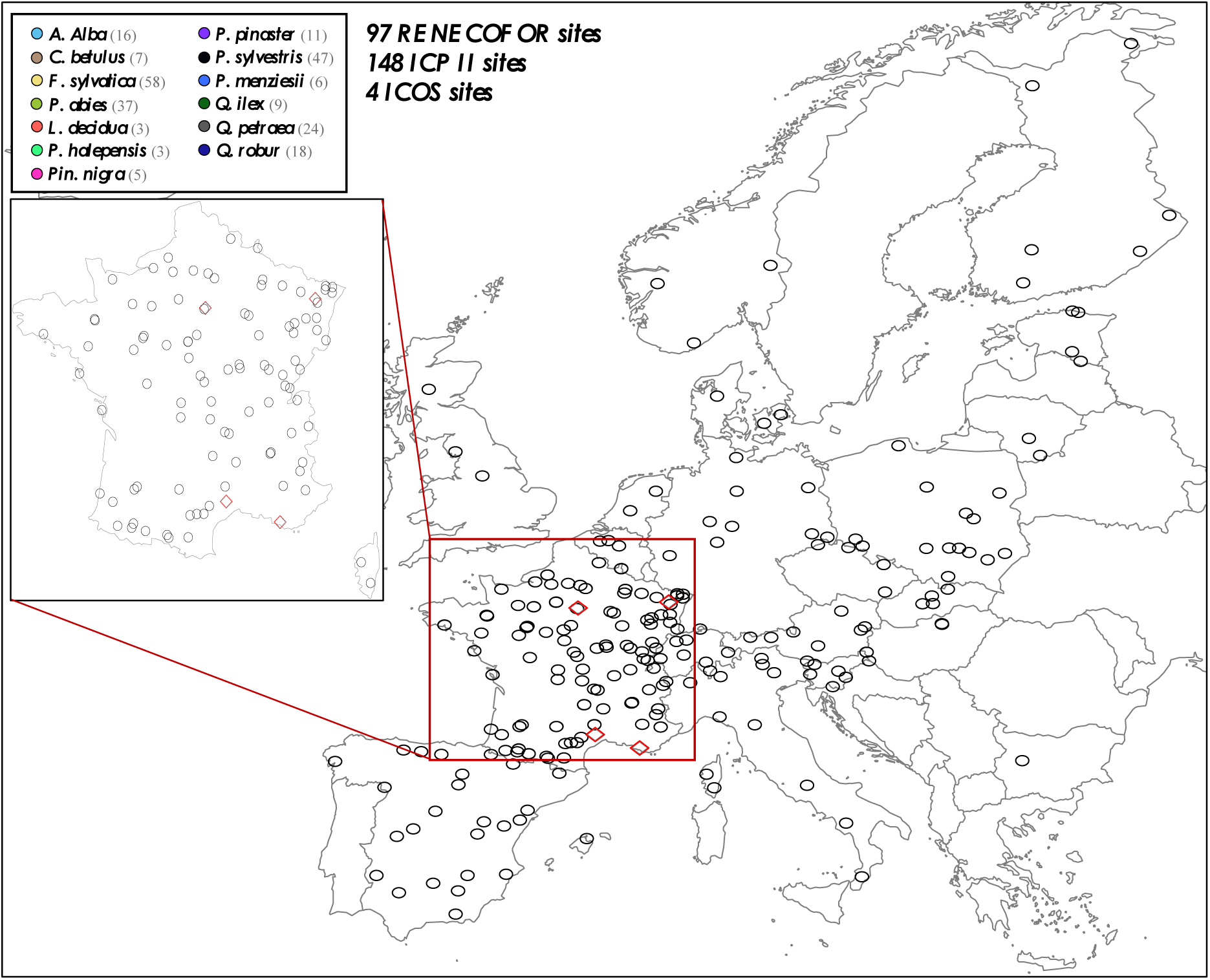
Spatial distribution of sites used for PHOREAU validation. Sites are color-coded based on the dominant species identified in the inventory (see legend in top-left). Red-bordered diamonds represent the four ICOS site (Puéchabon, Font-Blanche, Barbeau, and Hesse) selected for in-depth hydraulic validation.

*The RENECOFOR network*. Following the framework of original ForCEEPS validation, the RENECOFOR permanent forest plot network was used as the primary validation dataset (Ulrich, 1997). Comprised of 102 roughly half-hectare plots in even-aged managed forests, each comprised mostly of a single dominant species, they cover most of the main tree species and environmental conditions in France — with the notable exception of Mediterranean conditions. From the year 2000 onwards, the plots were exhaustively inventoried every five years, as well as before and after every thinning. After the removal of the plots that had suffered the most perturbations — and in particular the 1999 windstorm — 97 plots remained. With these we constructed 192 testing datasets, by grouping for each plot between 2000 and 2021 every pair of inventories that were separated by a period of at least four years within which no disturbance was recorded. The mean initial basal area of the plots was 28.3 m^2^/ha, while the time-interval between inventories ranged from 4 to 15 years, averaging at 7.1 year. As a rule, we avoided longer time-lapses, which would have mechanically improved simulation results, while giving less information on true model performance.

*The ICP II network*. In addition, we also used 148 plots from the International Co-operative Program on Assessment and Monitoring of Air Pollution Effects on Forests (ICP Forests), which comprises a network of intensively monitored forest sites (level II plots) distributed across Europe (de Vries *et al*., 2003; Schwärzel et al., 2022). These plots, located in other European countries, allowed the testing of the model over a wider range of abiotic and biotic conditions. This extension of the validation protocol, relative to the original ForCEEPS validation, was facilitated by the fact the RENECOFOR network is the French declination of the European-level ICP II program, with comparable protocols and measurements. Unlike for RENECOFOR, each plot corresponds to exactly one simulation dataset. The mean initial basal area of the plots was 28.1 m2/ha, while the time-interval between inventories ranged from 2 to 10 years, averaging at 4.6 year (refer to Table S2 for details on each individual simulation dataset).

### 3.4 Validation against intra-annual stand fluxes and tree hydraulics

For each of the four ICOS simulations, model predictions were compared to observations at two distinct levels: first for stand-level structure, focusing on the annual evolution of basal area, tree mortality, and leaf area index; second at the daily level, examining stand fluxes and tree functional dynamics. Statistical metrics were applied solely at the daily level, while annual stand structure predictions served as a baseline to identify discrepancies between observed and predicted fluxes (but refer to Sections 4.5 and 4.6 for direct validations on stand productivity and leaf area).

The performance of the PHOREAU model in reproducing the hydraulic functioning of forest stands was assessed for the following variables (from the most aggregative to the most specific): stand real evapotranspiration (ETR); evolution of soil water quantity (SWC); tree transpiration derived from sapflow; and stem water potential. Model performance was assessed using the Pearson correlation coefficient (*r*), the root mean square error (RMSE) and the mean deviation (MD) between observations and model predictions.

### 3.5 Validation against productivity

For each of the 340 selected RENECOFOR and ICP II simulation plots, five patches of 1000 m^2^ were initialized using the inventory of the first inventory campaign (see Table S2). For each patch, trees were sampled at random within the first inventory, until the basal area per hectare of the simulated patch matched that of the original inventory. Sampling was done without repetition within each patch, but with repetition among patches. Trees that were absent from the second inventory or found dead were kept in the sampling in order to match simulated plots to real inventories, but were removed after for growth comparison. As the time step for validation was deliberately kept short, model mortality — either through stress, age or density — were deactivated for this productivity validation protocol, so as to have for each sampled tree the observed and simulated final diameter.

Trees of unknown species, or of species currently not parametrized for ForCEEPS, such as *Pyrus communis* or *Ilex aquifolium,* used one of the generic sets of parameters. In addition to mortality, model seedling regeneration was also deactivated, due to the short time scales considered. The *crown A1* ratio between tree height and foliage height was initially set at the species maximum value, and initialized with the canopy bootstrap algorithm (see Figure 3).

#### Height-diameter ratio interpolation

In order to leverage PHOREAU’s ability to reproduce stand light availability and microclimatic conditions based on the structure of modelled trees, we used the newly independent tree height variable (see section 3.1.2) as an input parameter. However, height measurements were only available for a subset of trees across all RENECOFR and ICP II plots. Therefore, for trees where only circumference was measured, we applied plot-specific LOESS local regressions (Cleveland and Loader, 1996) to estimate species height-to-diameter curves from available measurements. The variability in height-to-diameter relationships among plots can be seen in Figure 13 and Figure S4, contrasted with the fixed height-to-diameter formula used in the original ForCEEPS framework. The associated statistics presented in Table S3 highlight the general tendency of the formula to underestimate tree heights in our study sites (AB = –15.7%; Table S3); this is not necessarily surprising, as the RENECOFOR and ICP II sites mostly support denser, more productive stands, where trees prioritize height growth to compete for sunlight.

**Figure 13.**
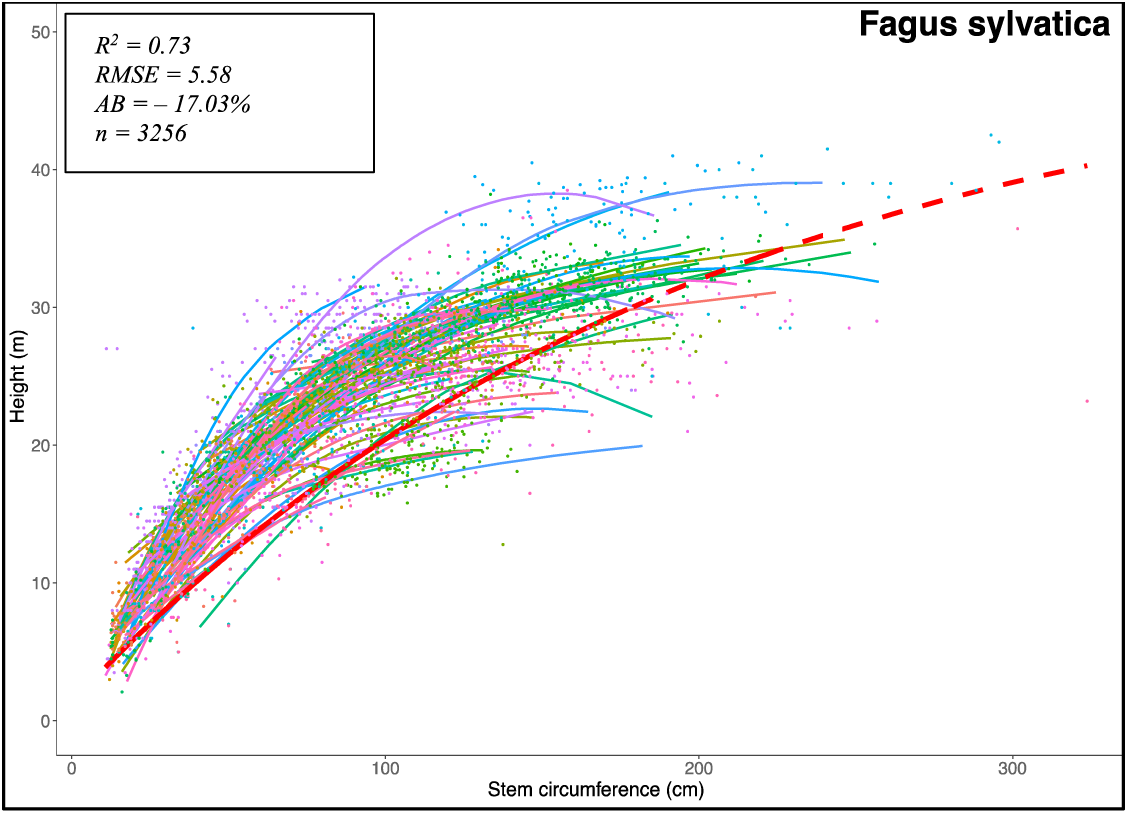
Diversity of site height-to-diameter curves for *Fagus sylvatica*. Refer to Annex X and Table X for details.

For both PHOREAU and the original ForCEEPS model, simulations were run for each site over the time periods indicated in Table S2, repeated five times for each of the five sampled patches. We compared simulated to observed basal area growth at both the tree scale and the stand scale, using predicted and observed basal area increments (BAI) normalized to mean annual values. While comparing actual, instead of averaged, annual increments would have constituted a stronger test, this data is not available for size of plots and the range of species and climatic conditions considered here. For stand-level comparisons, results were directly averaged over the five patches. The performance of both the PHOREAU and ForCEEPS model were assessed using the Pearson correlation coefficient (*r*), the root mean square error (RMSE) and the average bias (AB) between observations and model predictions.

### 3.6 Validation against leaf area index

The importance given to competition for light and leaf area prediction is one of the core principles of ForCEEPS, and the FORCLIM and FORECE models before it. However, because the initial models were focused on long-term forest dynamics, the methodology used to calibrate and validate the light competition module was based on the adequation between observed LAI values, and those reconstructed by the model after runs of hundreds or thousands of years starting from the bare ground (Kienast, 1987). Even then, LAI was not usually considered in the final validation, which was made on predicted biomass, basal area, tree density, or species composition (Bugmann, 1996; Wehrli *et al*., 2006). Notwithstanding the fact that this approach disregards past human interventions in the observed stands, it only accounts for equilibrium states, which becomes problematic when one wishes to apply the model at shorter timescales and consider the shorter-term effects of climate-change on existing forests. And while ForCEEPS did use actual inventories and short-term productivity for its validation, it did not directly test against predicted leaf area index (Morin *et al*., 2021).

This approach holds up as long as leaf area can be considered to be an intermediary variable. Because the previous models only used leaf area within the framework of their light competition modules, a given tree’s predicted leaf area only mattered insomuch as it provided shadow to neighboring smaller trees, decreasing their light availability factor. In this respect, absolute leaf area mattered less than the relative distribution between trees and species, which governed growth and final predicted compositions.

However, in PHOREAU, tree leaf area is also an integral input of another part of model: the simulation of hydraulic processes. This is because the upwards flow of water through the tree is ultimately driven by the transpiration in the leaves (Ruffault *et al*., 2022). And, in this respect, water flow is driven not by the relative, but by the absolute quantity of leaf area. Mechanically, a stand with a greater total leaf area index will tend to exhaust its water reserves faster; and tree leaf area, in ecosystems subjected to drought, is directly modulated by recent drought events (Bréda *et al*., 2006). These mechanisms, which are implemented in PHOREAU, require an accurate prediction of yearly stand leaf area index as a prerequisite condition to any simulation of hydraulic stress.

Unlike other validations of SurEau (Ruffault *et al*., 2023), the PHOREAU framework prevents the direct use of leaf area index as an input to the model; instead, the model initializes the stand LAI using solely the diameter and height information contained in the initial inventory. This makes the model suited to work on a majority of sites, where trunk diameters are measured but not leaf area, and allows it to make predictions in the future, as the LAI is recalculated on a year-to-year basis. The drawback of this approach is the addition of a new source of error when LAI is wrongly estimated. This is why, before validating the model on growth or drought-induced mortality, a preliminary validation of the leaf area index predictions was necessary.

This validation was realized on two different levels: first, by comparing model results to those obtained from satellite data for 340 sites spread over Europe featuring a large range of tree species; second, by comparing model results to LAI observations inferred from litter retrieval experiments for a few dozen sites in France.

The LAI satellite data used was retrieved from the Copernicus Global Land Service timeseries derived from daily PROBA-V satellite observations between 1999 and 2020 — first at a 1km resolution, then at 300m from 2014 (Fuster *et al*., 2020). For all RENECOFOR and ICP II sites and dates used for productivity validation (see Table S2) we compared LAI values predicted from the inventories at the start of the simulation, to those observed by PROBA-V and averaged over the summer months of the given year.

LAI validation on litter data was restricted to those RENECOFOR sites where such experiments had been realized — mostly beech and oak sites, excluding coniferous-dominated stands not suited to litter retrieval (Guillemot, unpublished data).

### 3.7 Validation against potential natural vegetation

To validate the model’s prediction in the long-term for a broad range of climatic conditions, integrating the effects of all the different processes for mortality, reproduction, phenology, microclimate and competition not directly captured by shorter-term validations protocols, we compared community compositions simulated by PHOREAU with the predicted potential natural vegetation (PNV) along an environmental gradient. Here, similarly as in Bugmann (1996) and Morin et al. (2021), potential natural vegetation is simply defined as the assumed dominant tree species, assuming no large disturbances, in a space spanned by mean annual precipitations (MAP) and mean annual temperatures (MAT), following Ellenberg (1986), Rameau *et al*. (2008), and San-Miguel-Ayanz *et al*. (2016). For this validation we used for the same 250 RENECOFOR and ICP II sites used for the productivity validation, spanning across all the different PNV conditions described in Ellenberg (1986) (Figure 12.

For each of the 250 sites, we ran 2000-year simulations starting from the bare ground. This simulation length – accounting for seedling establishment, tree growth and mortality – was necessary to ensure the communities were no longer in a transient phase, and had reached the final stage of forest succession with a pseudo-equilibrium composition. The 2000-year climate time series were obtained by randomizing the years for which climatic data was available (1969-2020), which preserved inter-annual variability in in climate, but avoided any cyclic trend. For each site we considered of 50 independent patches of 1000 m^2^. At the end of the simulation, aggregate species basal areas per hectare were extracted for each simulated site, and compared to assumed PNV dominant species.

## 4 Result Analysis

### 4.1 Prediction of above-ground tree growth

PHOREAU demonstrated satisfactory predictive capability for tree-level mean annual basal area increment (BAI) across diverse species and climatic conditions throughout Europe (Figure 14). The model achieved a strong correlation between observed and predicted values (*r* = 0.68, p < 0.001, *n* = 81655; Table S4), with satisfactory levels of prediction accuracy (RMSE = 0.00106, AB = 0.225, and AAB = 0.793). However, the model exhibited an attenuating effect on the observed variability in tree growth, tending to underestimate the productivity of the most vigorous trees while simultaneously overestimating growth in less productive trees (Figure X).

**Fig. 14.**
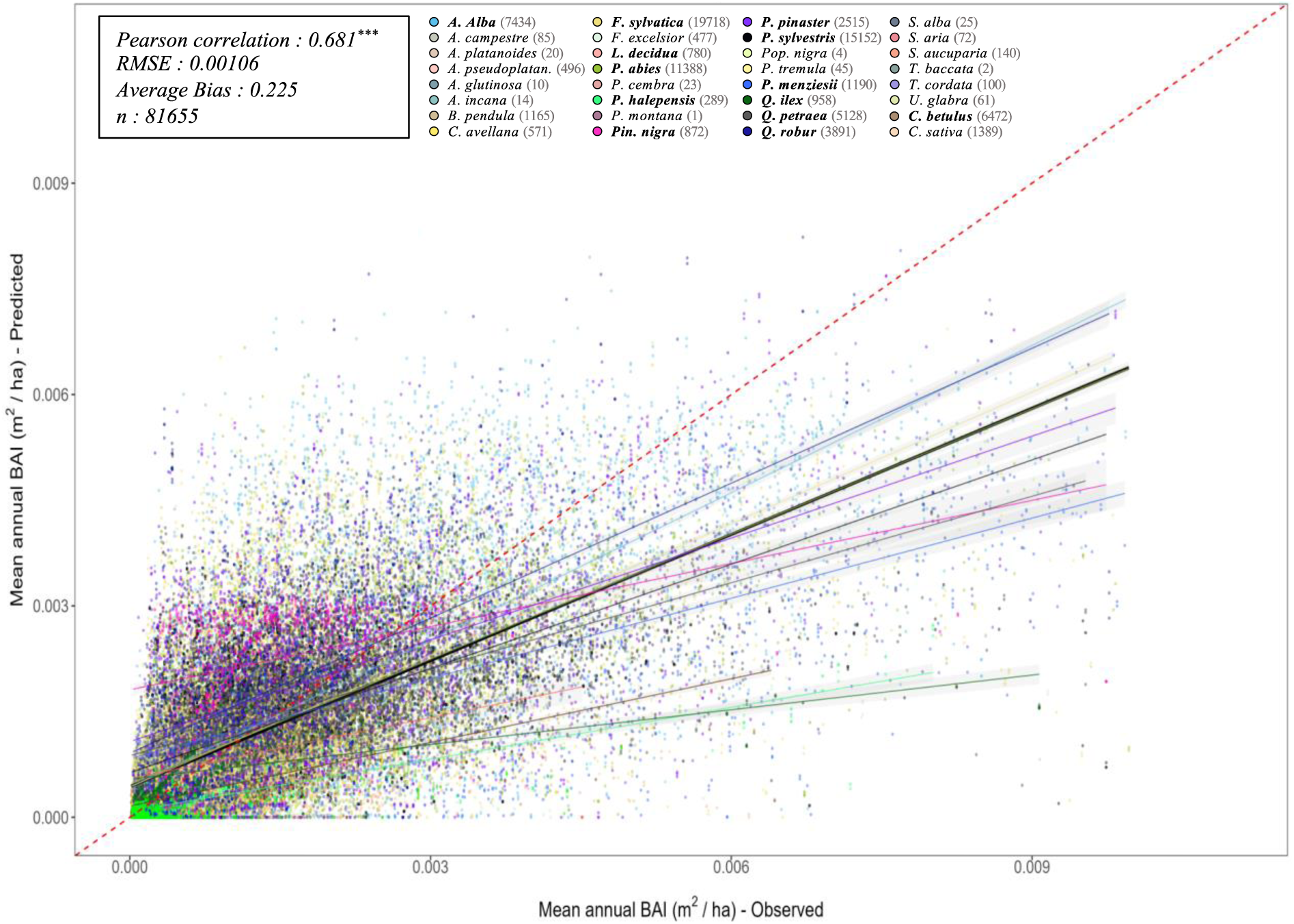
Projected (by PHOREAU) against observed mean annual tree basal increments (BAI) for all simulated trees over the 340 RENECOFOR and ICP II validation inventories. Tree points are color coded by species (see legend above). The dashed red-line is the 1:1 line; other full lines represent the regression lines of the linear model between observed and predicted tree productivity, with confidence intervals represented by the grey shaded area (in black the overall regression; coloured lines for species-specific regressions). Species-specific regressions are only shown for stand dominant species (in bold in legend) Associated statistics for the global simulation in top left, while species-specific statistics can be found in Table SI.

When assessed at the species level, the Pearson correlation coefficients varied substantially, from 0.14 for *C. avellana* to 0.913 for *U. glabra* (Table S4). Prediction accuracy also differed widely among species, with an average RMSE of 0.00103 and an AB of 0.34. Correlation coefficients were generally higher for the main species of the study compared to secondary species (average *r* = 0.60 and 0.53, respectively), with a pronounced tendency for the model to underestimate the productivity of these secondary, generally understory species, whose growth rate was not recalibrated on forest growth data in the original ForCEEPS study (Morin *et al*., 2021).

In comparison with the original ForCEEPS model, which was applied to the same dataset (Figure S1, Table S4), PHOREAU demonstrated a moderately improved performance in predicting tree productivity. It yielded higher Pearson correlation coefficients, as well as lower RMSE and absolute errors. Despite these improvements, PHOREAU’s predictions exhibited a comparatively greater average bias.

### 4.2 Prediction of above-ground stand growth

At the stand level, PHOREAU exhibited robust performance in reproducing mean annual BAI across most species and environmental conditions. Overall, there was a strong correlation between observed and predicted values across all 340 simulations (*r* = 0.62, *p* < 0.001) with a small margin of error between observations and predictions (RMSE = 0.23, AB = 3.7%, and AAB = 0.34; Figure 15, Table S5). However, the accuracy varied when species were analyzed individually. While the model generally showed no significant systematic bias (RMSE = 0.23; AB = –2.2% on average), some species exhibited notable biases and variability, particularly in the most productive plots where the model tended to underestimate productivity. This was especially evident *for P. halepensis* (RMSE = 0.35, AB = –65%) and *P. menziesii* (RMSE = 0.18, AB = –29%), though both had relatively small sample sizes. Even for *Q. petraea* (RMSE = 0.19, AB = –17%), where sample size was not a limitation, similar bias was observed.

**Fig. 15.**
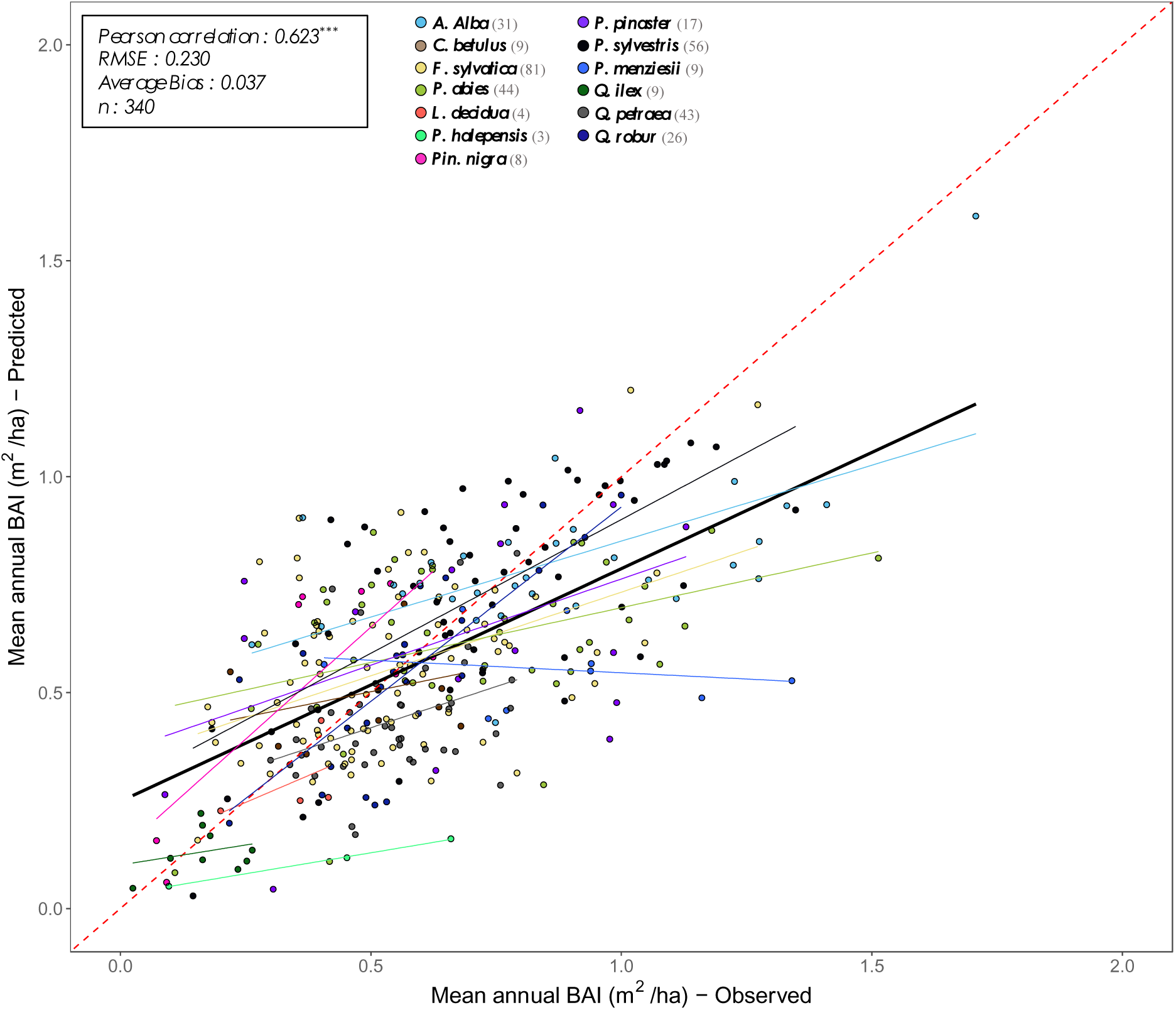
Projected (by PHOREAU) against observed mean annual stand basal increments. **(BAI)** for all 340 RENECOFOR and ICP II validation inventories. Stand points are color coded by dominant species (see legend above). The dashed red-line is the 1:1 line; other full lines represent the regression lines of the linear model between observed and predicted stand productivity, with confidence intervals represented by the grey shaded area (in black the overall regression; coloured lines for species-specific regressions). Associated statistics for the global simulation in top left, while species-specific statistics can be found in Table S2.

When examining the relationship between prediction errors and various stand characteristics (Figure 16), no strong systematic biases were identified with respect to site-specific factors such as rainfall, temperature, stand density, or simulation duration. However, the regression analysis revealed a weak but statistically significant positive relationship between errors and site water-holding capacity (WHC) (slope = 0.0034, *r* = 0.138, *p* < 0.05), suggesting a tendency to underestimate productivity on drier soils. Additionally, there was a strongly significant negative relationship between errors and initial stand basal area (slope = –0.0044, *r* = –0.21, *p* < 0.001), indicating that the model underestimates productivity in the most productive stands.

**Fig. 16.**
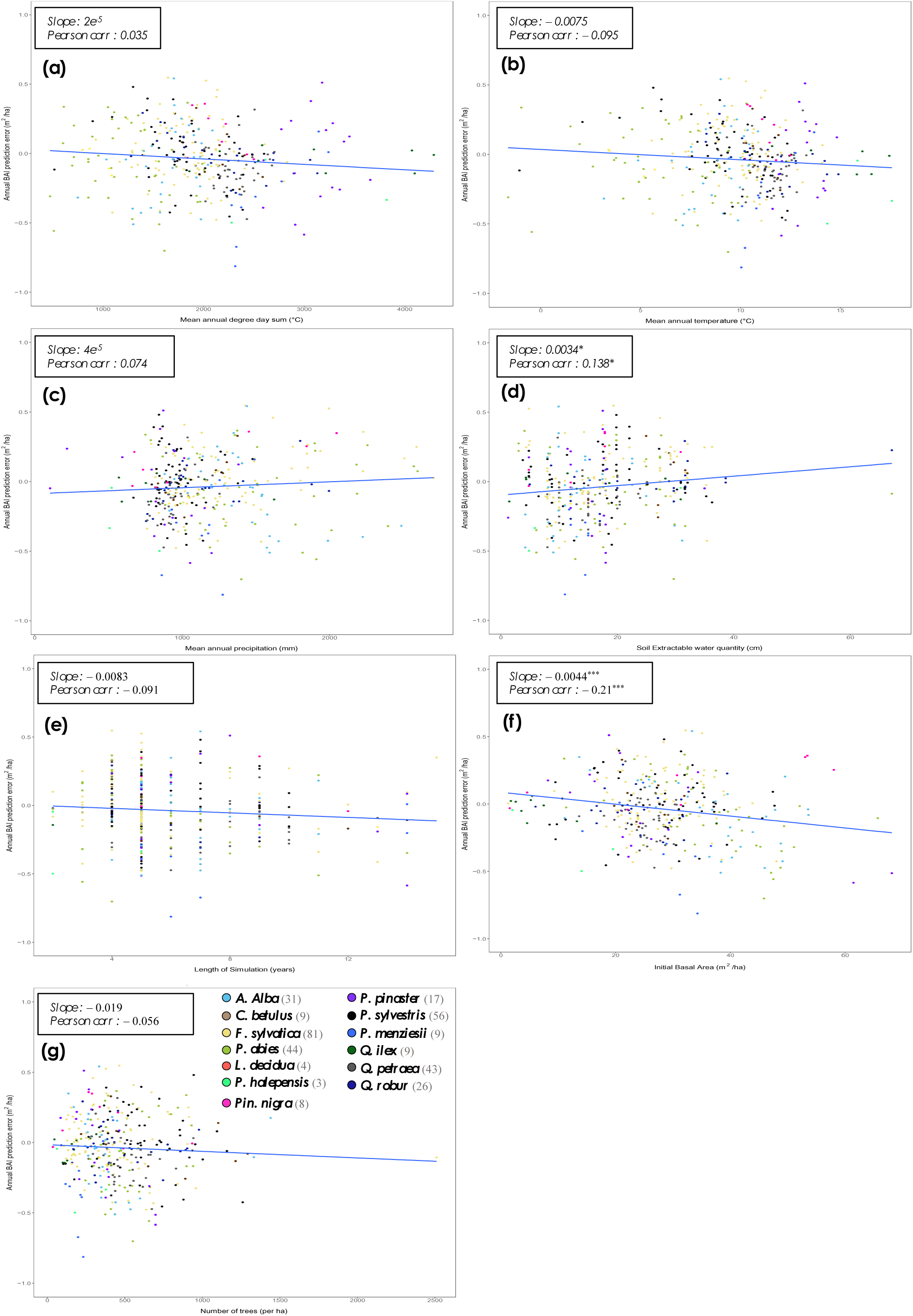
Regressions of PHOREAU stand basal area increment (BAI) prediction errors against site characteristics. For each of the 340 simulation inventories, the mean annual BAI prediction error, color coded by site dominant species (see legend in bottom left) is plotted against: (a) Mean annual degree day sum (°C); (b) mean annual temperature (°C); (c) mean annual precipitation sum (mm/’year); (d) soil maximum available water capacity (cm); (e) length of simulation; (f) initial inventory basal area (m^2^/ha). (g) initial stand density (trees/ha). The plain blue line is the regression line of the linear model of the relationship between prediction error and stand characteristic, with slope value and significance in top left, along with Pearson correlation.

In comparison to the original ForCEEPS model applied to the same dataset, PHOREAU demonstrated enhanced predictive accuracy across all evaluated metrics. PHOREAU produced a higher Pearson correlation coefficient (*r* = 0.62 vs. *r* = 0.53 for ForCEEPS), along with lower RMSE (0.23 vs. 0.316), average bias (AB = 3.7% vs. 7.7%), and average absolute bias (AAB = 0.34 vs. 0.44; see Figure S2, Table S5). These results highlight PHOREAU’s improved capability in predicting stand productivity compared to ForCEEPS

### 4.3 Prediction of leaf area index based on inventories

PHOREAU demonstrated a reasonable capacity to estimate stand leaf area index (LAI) from inventory data across diverse species and site conditions throughout Europe. When compared to PROBA-V satellite data (Figure 17), the model yielded a good correlation between observed and predicted LAI values (*r* = 0.55, p < 0.001, n = 340; Table S6), with acceptable prediction accuracy (RMSE = 1.41, AB = 0.08). Although no significant systematic bias was detected, the model tended to dampen the observed variability in LAI, slightly underestimating LAI in denser forest canopies while overestimating it in more open plots.

**Fig. 17.**
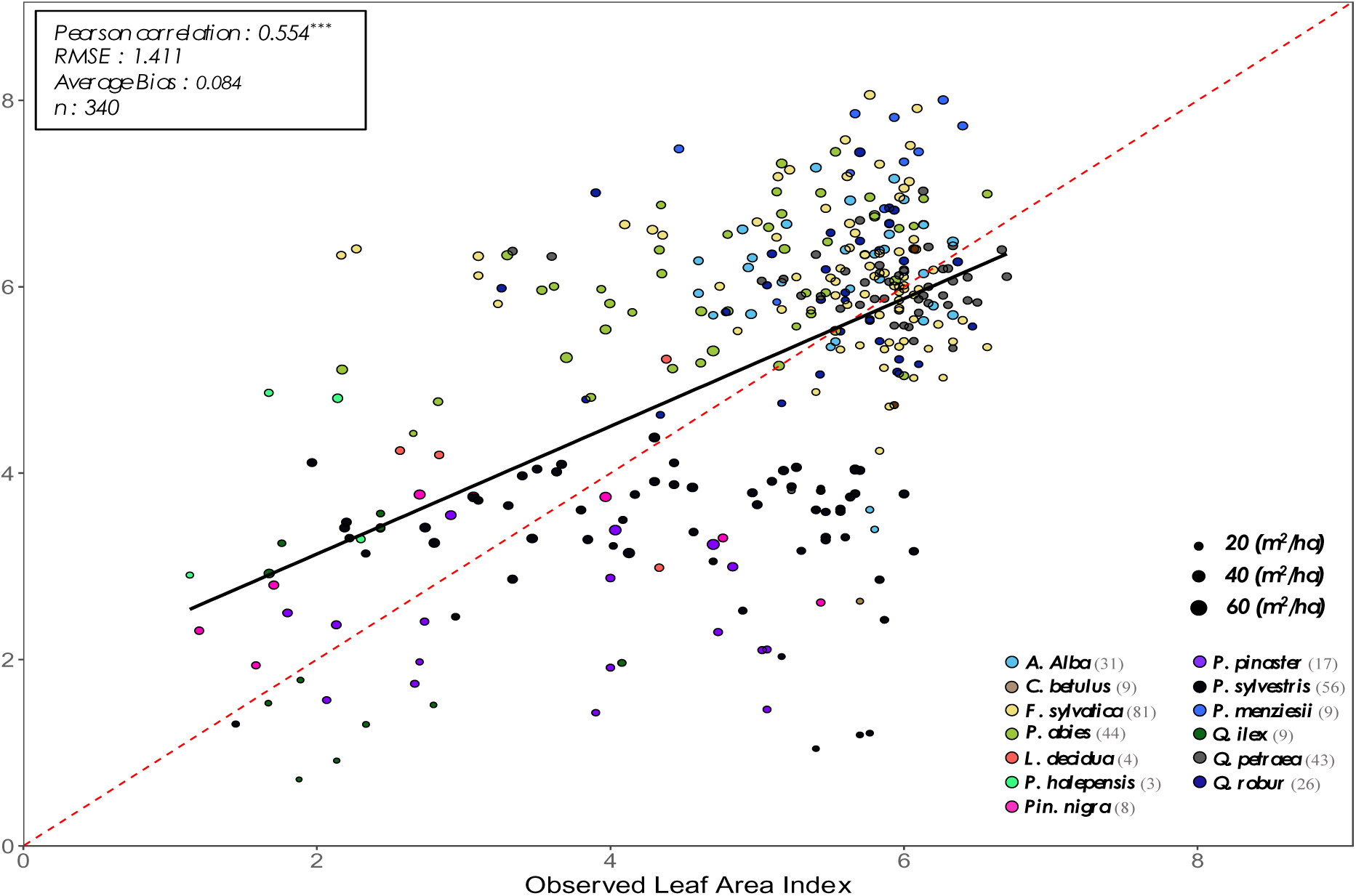
Projected (by PHOREAU) against observed satellite leaf area index. **(LAI)** for all 340 RENECOFOR and ICP II validation inventories. The y-axis shows the LAI predicted by the model from the stand inventory at the start of the simulation, while the x-axis represents the PROBA-V LAI value for the maching coordinate and inventory year, averaged between July, August and September. Stand points are color coded by dominant species (see legend in bottom left). The size of points shows inventory basal area. The dashed red-line is the 1:1 line; the black full line represent the regression line of the linear model between observed and predicted LAI, with confidence interval represented by the grey shaded area. Associated statistics in top left.

A species-specific analysis revealed notable biases for certain species. The model consistently overestimated the LAI of dense coniferous plantations, particularly for species such as *P. abies*, *A. alba*, and *P. menziesii*. Conversely, it significantly underestimated LAI for low basal area inventories dominated by *P. pinaster, P. sylvestris*, which could partially result from discrepancies between inventories observed and simulated before and after thinnings. Overall, while PHOREAU presents a notable improvement in capturing inter-species LAI variability compared to the original ForCEEPS model (RMSE = 3.42, AB = 0.49; Table S3), and does not show any significant bias compared to LAI litter inferred from litter collection (RMSE = 0.65, AB = –0.03, n = 40; Table S7, Figure 18), it struggles to distinguish smaller variations in LAI within similar-sized plots of the same species (*r* = 0.3, p = 0.047; Table S7).

**Fig. 18.**
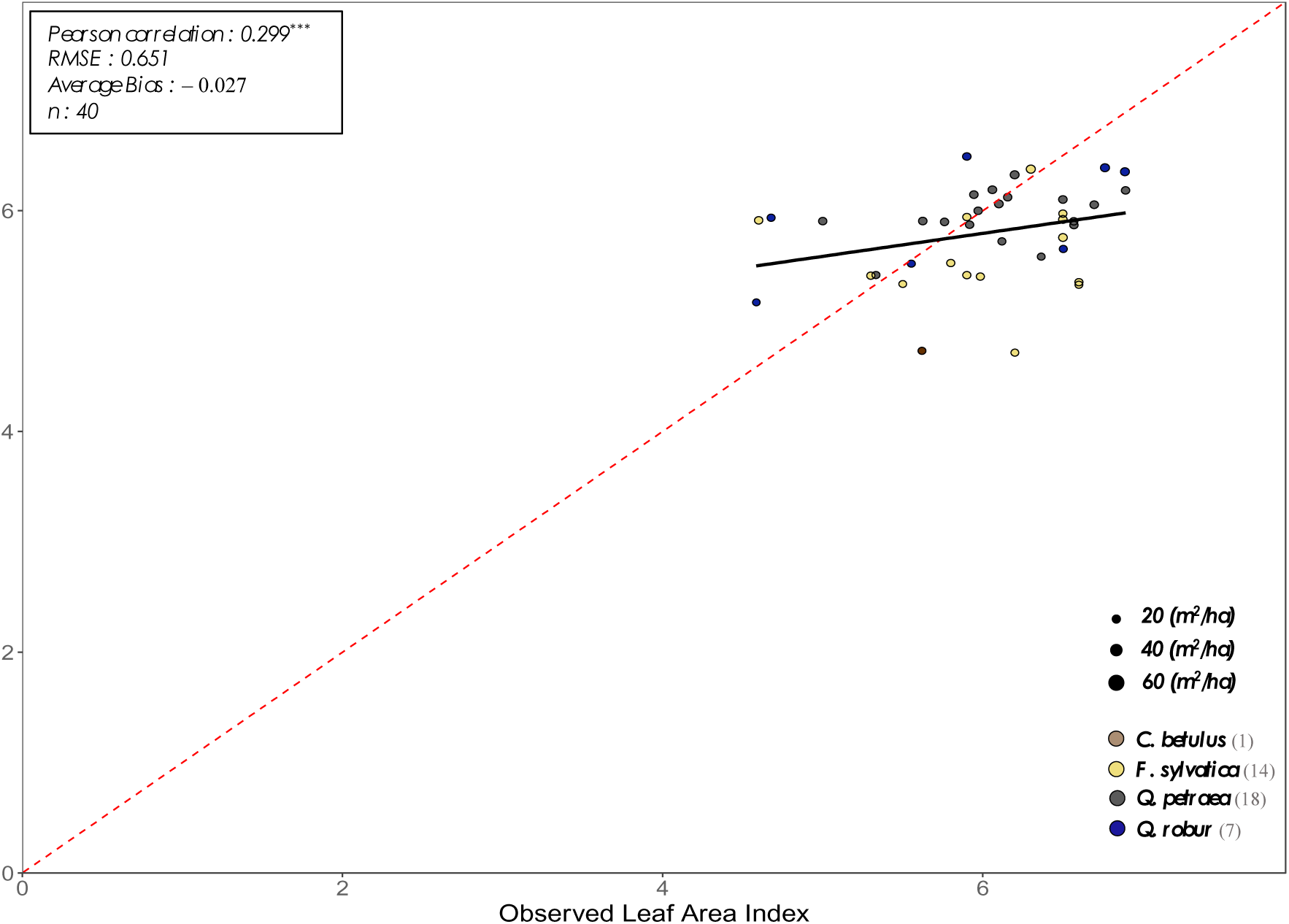
Projected (by PHOREAU) against observed litter leaf area index. **(LAI)** for available RENECOFOR inventories. The y-axis shows the LAI predicted by the model from the stand inventory at the start of the simulation, while the x-axis represents the LAI value infered from litter collection for the maching coordinate and closest available year. Stand points are color coded by dominant species (see legend in bottom left). The size of points shows inventory basal area. The dashed red-line is the 1:1 line; the black full line represent the regression line of the linear model between observed and predicted LAI, with confidence interval represented by the grey shaded area. Associated statistics in top left.

### 4.4 Prediction of long-term species composition

When comparing the distribution of predicted dominant tree species at the conclusion of 2,000-year simulations along the environmental gradient covered by 250 sites across Europe (Figure 19) it is clear that PHOREAU’s ability to accurately predict potential natural vegetation (PNV) varied depending on site conditions. Overall, the model performed well, with 62% of predictions accurately matching observed community compositions, and 24% partially accurate predictions. However, prediction uncertainty was notably higher for Mediterranean forest types, humid beech forests, and mixed montane spruce-beech forests.

**Fig. 19.**
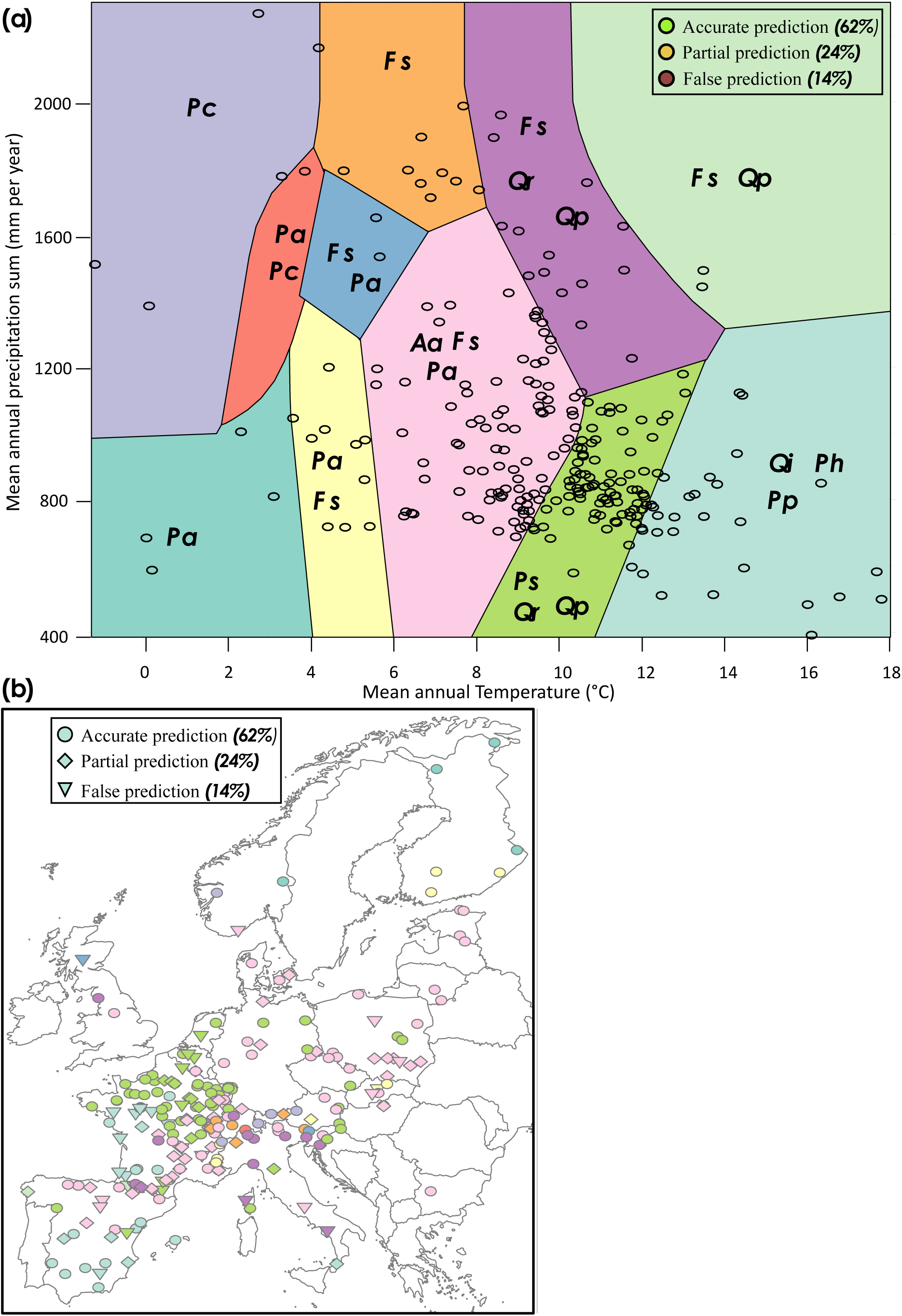
(a) Distribution of the 250 tested sites in the PNV diagram of supposed dominating species (built according to mean annual temperature and precipitation sum). PNV dominating species are Pc (P. cernbra); Pa (P. cties); Aa (A dba); Fs (F. sylvatica); Qp (Q. petrosa); Qr (Q. robur); Pp (P. pnosfer); Ph (P.hdepens is); Qi (Q. ilex) Circle colors indicate the agreement between simulated and PNV dominating species after the 2000 years PHOREAU simulations. Green: sites for which the dominating species was accurately predicted. Orange: sites for which the second-ranked (by basal area) species was accurately predicted, but not the first-ranked. Red: sites for which neither the first-ranked nor second-ranked species were accurately predicted. **(b) Geographical repartition of the 250 sites** (RENECOFOR and ICP II) used for PNV validation, colored by potential niche composition. Shapes indicate prediction success, as described above.

A detailed analysis of the predicted dominant species (Figure 20) revealed that much of this uncertainty stemmed from PHOREAU’s tendency to overestimate the competitive advantage of *Q. robur* in both hot and milder climates, while underestimating the competitiveness of *Q. petraea* and *F. sylvatica*. Despite these discrepancies, the model demonstrated strong predictive performance in extreme environments, accurately predicting species composition at both extremely cold and extremely hot sites.

**Fig. 20.**
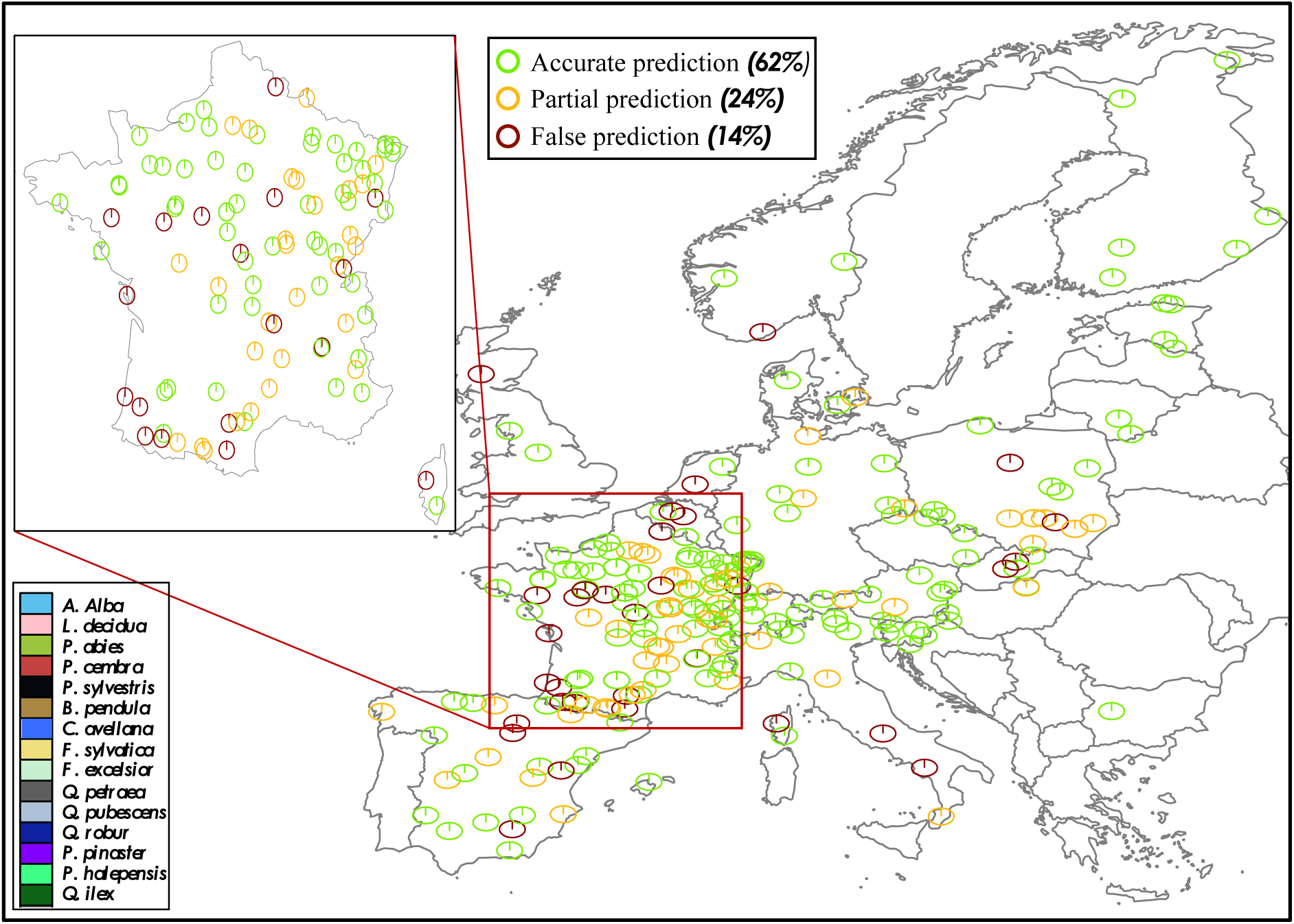
Projected community compositions after long-term PHOREAU simulations. For the 250 tested sites of the PNV validation, each pie graph represents the basal area repartition of simulated species after 2000 years (see legend in bottom left for species color code). Prediction success (according to PNV assumed dominant species) is represented by the color of the circle’s outer border. Green border: sites for which the dominating species was accurately predicted. Yellow border: sites for which the second-ranked species was accurately predicted, but not the dominating species. Red border: sites for which neither the first-ranked nor second-ranked species were accurately predicted.

### 4.5 Prediction of intra-annual stand fluxes and tree hydraulics

The results of the in-depth validation for the four considered ICOS sites demonstrated a strong ability of the PHOREAU model to replicate observed data across a wide range of metrics, from broad stand-level characteristics to specific physiological responses The model closely followed observed trends in stand basal area, despite the inherent challenge of predicting individual tree mortality (Figure 21). It accurately captured both the magnitude and variability of dieback across sites, in terms of both tree density (Figure 23) and basal area loss (Figure S5), with a marked increase in the rate of basal area loss in the latter years of each simulation; however, the model slightly overestimated mortality across all sites and particularly at *Hesse*, as well as the share of large tree death relative to medium trees and saplings. Predicted foliage area results aligned well with observations in the two open evergreen sites (*Puéchabon* and *Font-Blanche*, Figure 22). PHOREAU captured the quick regrowth in foliage area observed at *Hesse* after the 2005 cut (Granier *et al*., 2008); however, when comparing absolute values, PHOREAU noticeably underestimated foliage area in the two denser deciduous forests, consistent with prior validation results on leaf area (see Section 5.3). Despite these biases, the overall alignment between predicted and observed forest dynamics provides a solid foundation for comparing stand functioning and tree physiological responses at finer temporal resolutions.

**Fig 21.**
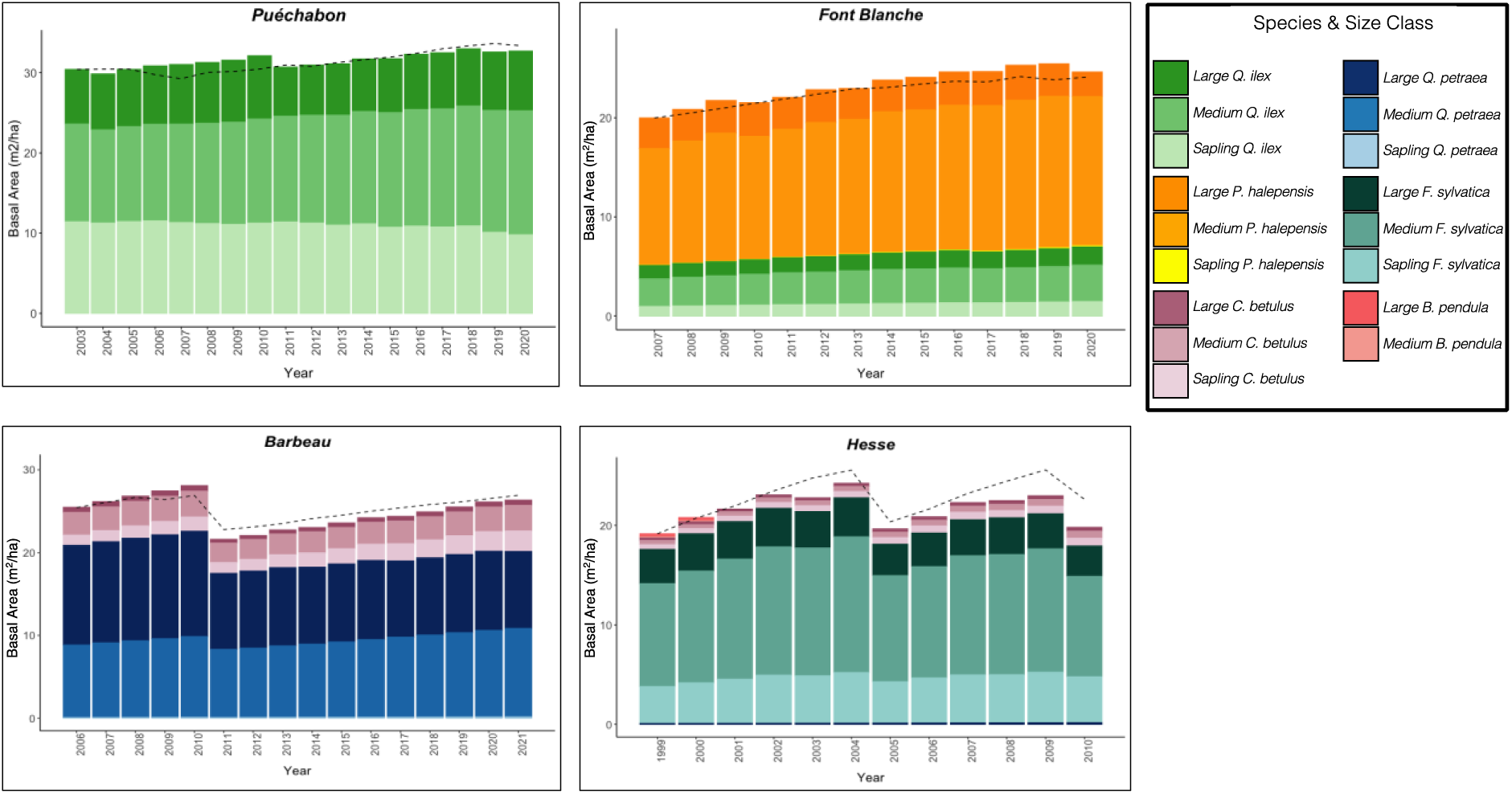
Predicted versus observed evolution of annual stand basal area. For each simulation site. the bars depict the annual basal area projections generated by the PHOREAU model, broken down by species and size class contributions (refer 1o Annex X (or details). The dashed line represents the observed annual total basal area derived from inventory data. Basal area is defined as the cross-sectional area at breast height o’ all trees per hectare.

**Fig 22.**
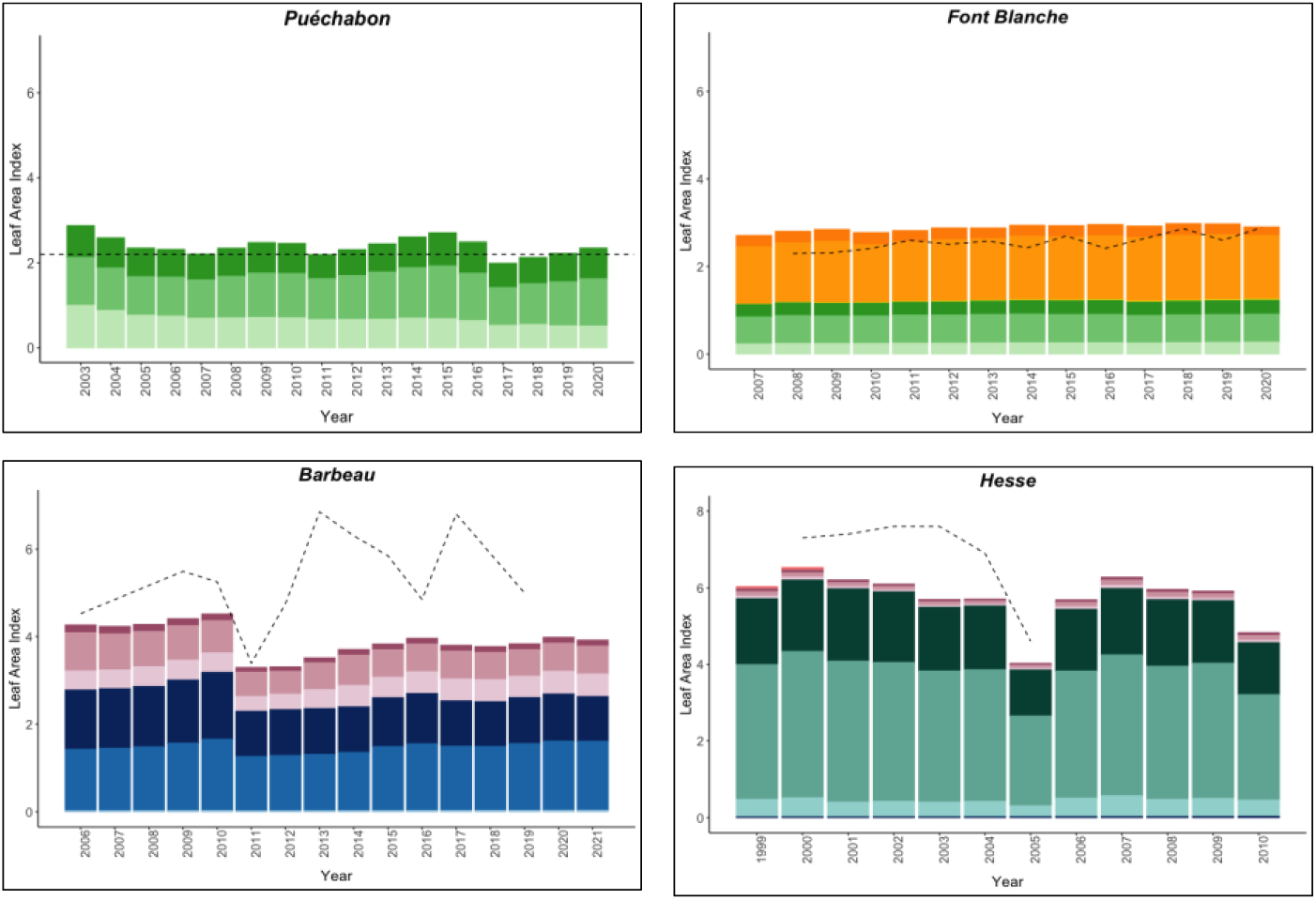
Predicted versus observed annum stand leaf area index (LAI). For each simulation site, the bars depict the annual leaf area index projections generated by the PHOREAU model, broken down by species and size class contributions (rate-to Annex X for details). The dashed line represents the observed annual stand leaf area index (data sources detailed in Annex X). Leaf area index is defined as the total ons-sided leaf area per unit of ground area.

**Fig. 23.**
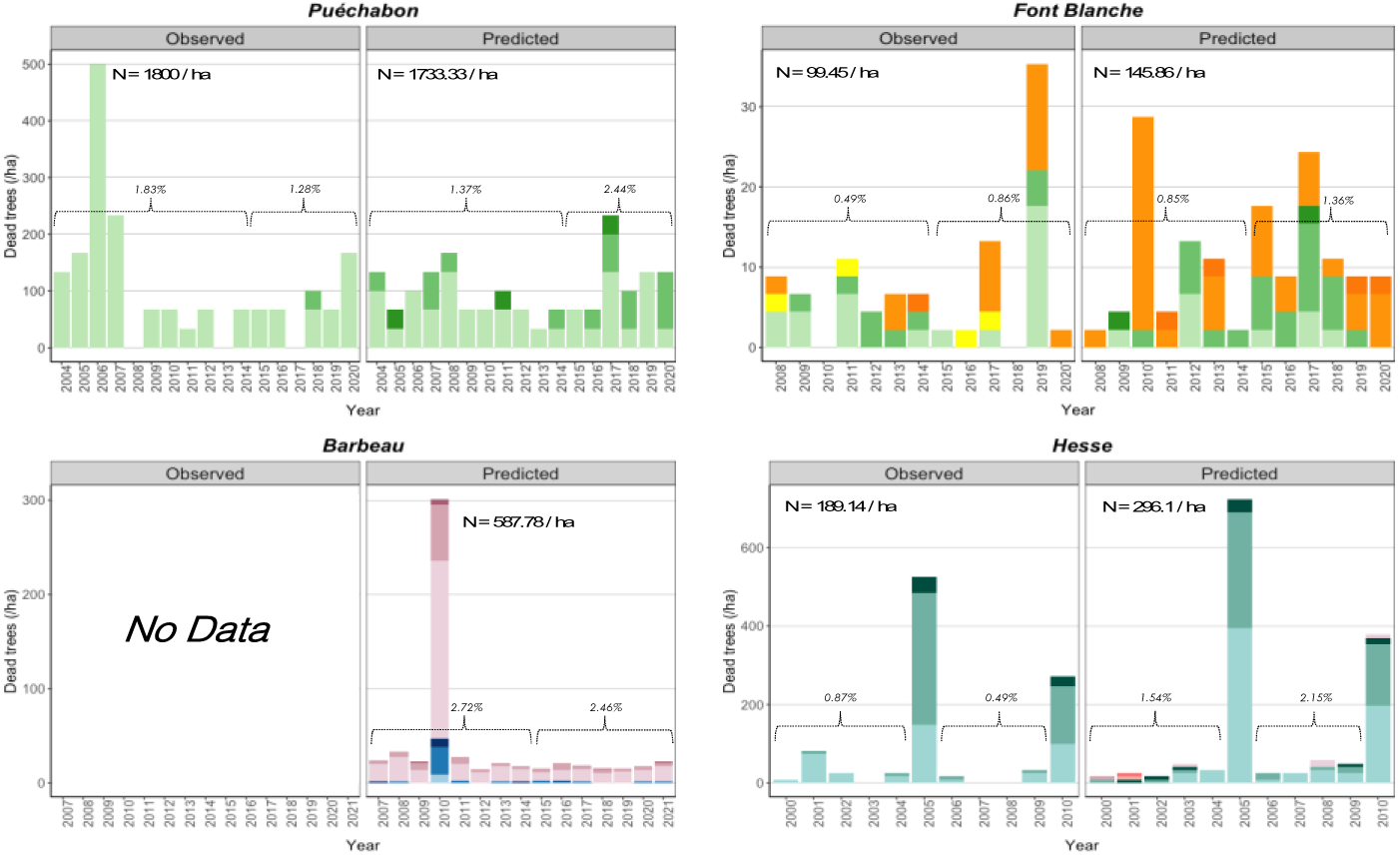
Predicted versus observed annual free mortality. For each simulation site, the bars depot the total annual number of dead trees, irrespective of cause. broken down by species and size class contributions (refer so Annex X for details) Observed values are derived from stand inventories. while predicted values are geneated by the PHOREAU model. Also shown are the annual mortality rates, calculated relative to the initial number of trees for Sao distinct time periods in each simulation, along with the total number N of dead trees by hectare Transparent bars indicate years with thinnings (see Annex X for details), which are excluded from the mortality statistics.

The PHOREAU model demonstrated solid performance in predicting daily evapotranspiration (ETR) across three of the four ICOS sites, with low mean deviations (*Puéchabon*: 0.03; *Barbeau*: –0.24; *Hesse*: 0.8) and strong Pearson correlations (*Puéchabon*: 0.64; *Barbeau*: 0.79; *Hesse*: 0.62) between observed and predicted values (Figure 25). At *Font-Blanche*, correlation was moderate (r = 0.48, p < 0.001), as the model underestimated summer ETR while overestimating winter and autumn ETR. This discrepancy, particularly the underestimation of Q. ilex transpiration (Figure 24), likely stems from biases in the model’s allocation of leaf area between Q. ilex and P. halepensis and a dampened response of P. halepensis stem water potential to summer drought (Figure **31**). Over time, the differences between predicted and observed monthly cumulative ETR became more pronounced, reflecting a drift from real-world conditions. The model also underestimated ETR during the leafless winter months at *Barbeau* and *Hesse*, which could result from excluded understory shrubs.

**Fig. 24.**
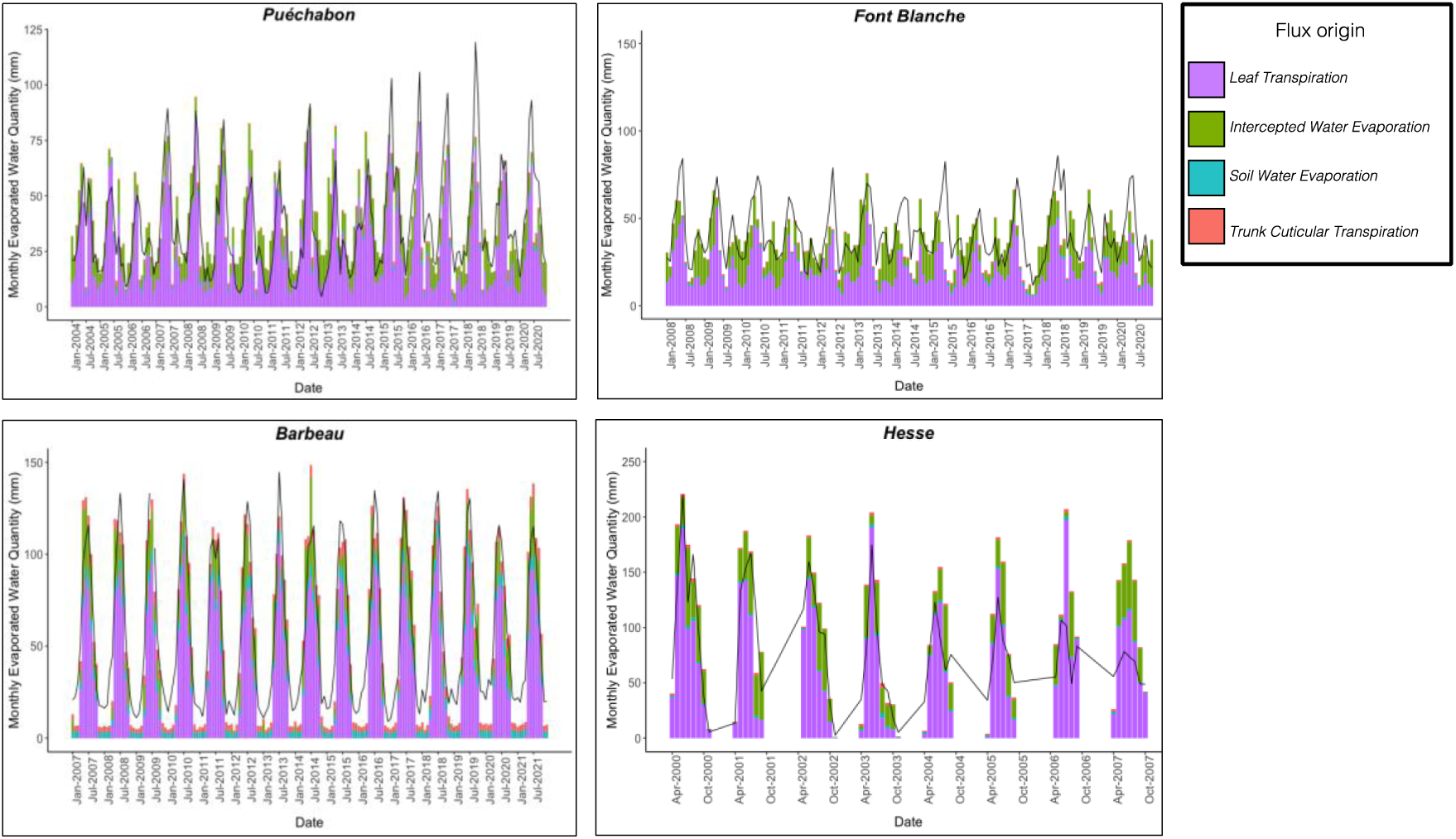
Predicted versus observed evolution of monthly real Evapotranspiration (ETR). For each simulation site, the bars depict the monthly ETR (mm) projections generated by the PHOREAU model, broken down by source of flux. Soil and intercepted water evaporation respectively originate from the first layer of soil and the water stored on the surface of leaves, while the two other sources are transpiration from different compartments of the PHOREAU tree (refer to Annex X for details). The black line indicates the observed monthly actual evapotranspiration, representing the total water vapor released from the soil and vegetation into the atmosphere, aggregated from hourly or sub-hourly measurements obtained from each site’s flux tower.

**Fig 25.**
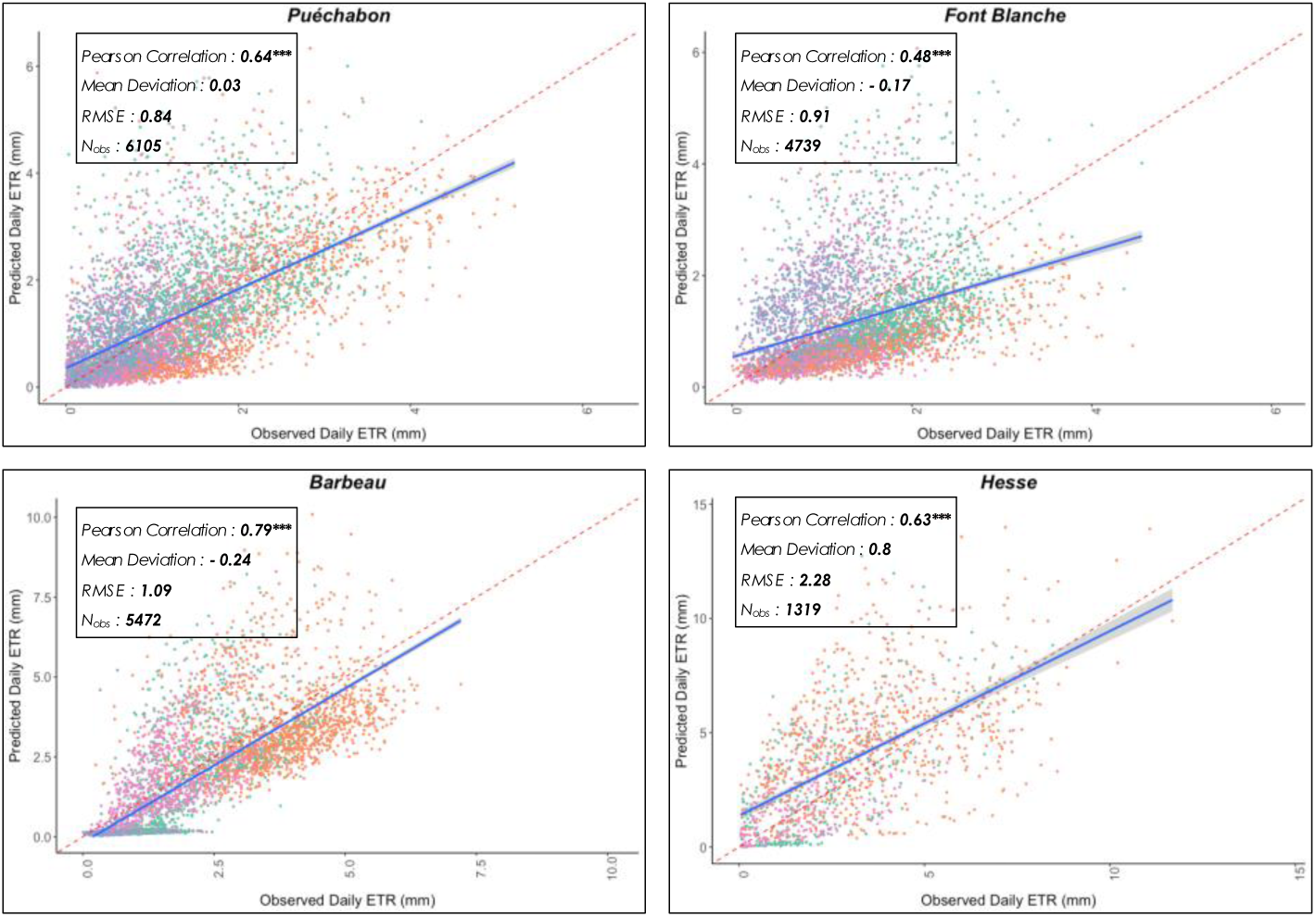
Predicted versus observed daily real evapotranspiration (ETR). For each simulation site, the plain blue line is the regression line of the linear model of the relationship between observed and predicted stand daily ETR, with confidence interval represented with the grey dashed lines; the dashed red line is the 1:1 line. See Annex X for definition of associated statistics. Colour code for the seasons as follows: 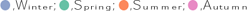

Across all sites, the PHOREAU model demonstrated robust performance in predicting the daily evolution of soil water quantity (SWC), with low mean deviations (*Puéchabon*: 15.4; *Font Blanche*: 1.03; *Barbeau*: –47; *Hesse*: –31.4; Table S12, Figure 27) and strong Pearson correlations (*Puéchabon*: 0.8; *Font Blanche*: 0.86; *Barbeau*: 0.92; *Hesse*: 0.78) between observed and predicted values. For model generally captured the seasonal refilling of soil water reserves well (Figure 26). However, at *Hesse*, model results noticeably lagged behind observations: this is consistent with the model’s overestimation of *F. sylvatica* stress during the 2003 drought (Bréda *et al*., 2006), and the overestimation of mortality and post-2003 stand ETR (Figure 24). The omission of a perched aquifer present in the site likely contributed to these discrepancies.

**Fig. 26.**
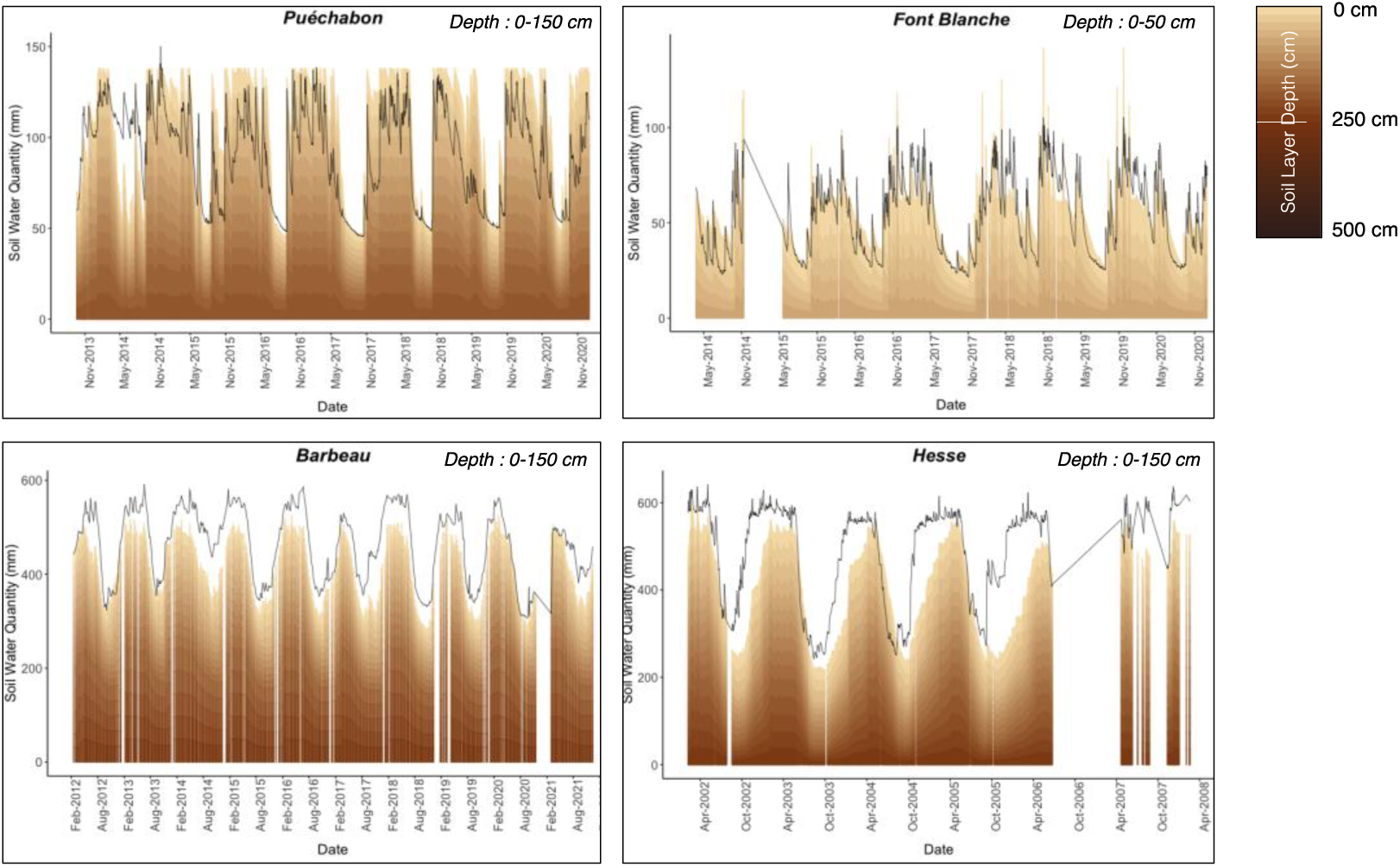
Predicted versus observed evolution of soil water quantity (SWC). For each simulation site, the black line indicates the observed daily actual SWC. The stacked bars depict the daily SWC (mm) projections generated by the PHOREAU model, with individual contributions of each soil layer stacked and color-coded by soil layer (see Fig. 6 for layer details). The projections are confined the maximum measured depth for the each site, as indicated in the upper right corner of the figure. For *Barbeau* and *Font Blanche,* observed SWC were directly obtained from site managers; for *Puechabon* and *Hesse,* they were interpolated from soil relative humidity (RH%) measured at different depths, using the same rock fractions as used in the simulation.

**Fig. 27.**
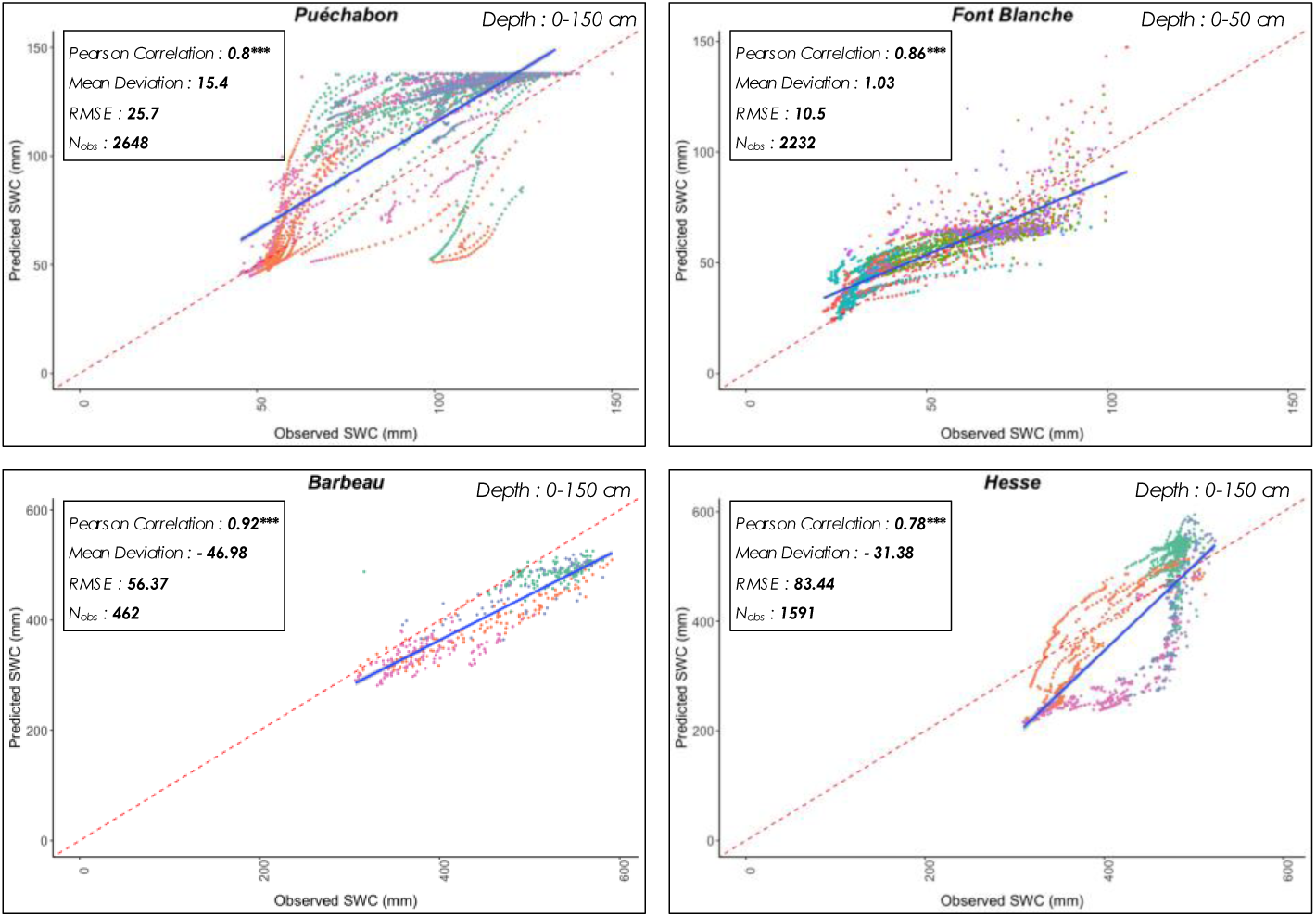
Predicted versus observed soil water quantity (SWC). For each simulation site, the plain blue line is the regression line of the linear model of the relationship between observed and predicted SWC, with confidence interval represented with the grey dashed lines; the dashed red line is the 1:1 line. See Annex X for definition of associated statistics. Colour code for the seasons as follows: 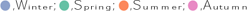

**Fig. 28.**
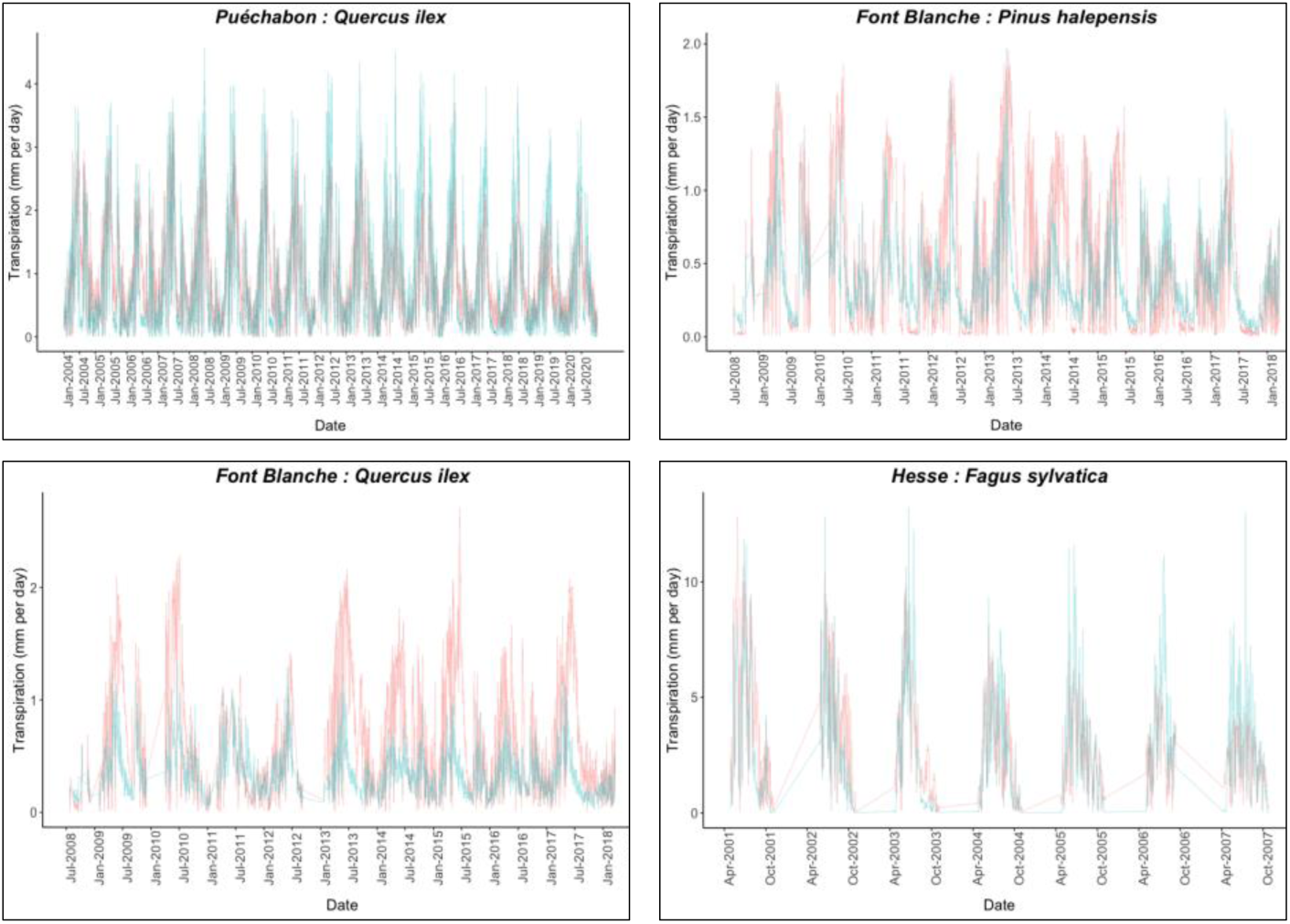
Predicted versus observed evolution of aggregate daily species transpiration. For each simulation site, the blue line depicts the aggregated daily transpiration (mm) generated by PHOREAU from all the trees of the given species. The red line depicts the observed daily transpiration value for this species, upscaled from sapwood measurements made for individual trees using stand LAI and species leaf area to sapwood area ratios.

The model also effectively captured the variability in measured stem water across species, seasons, and times of day (Figure **31**). It achieved strong correlations between observed and predicted values for both daily minimum stem potential (*r* = 0.71, p < 0.001, *n* = 208; Table S10) and predawn stem potential (*r* = 0.79, p < 0.001, *n* = 303; Table S9), with fair levels of prediction accuracy (RMSE = 0.92 and 0.89, respectively). Despite these strong correlations, the model tended to attenuate the range of observed potentials, underestimating predawn potentials (MD = –0.5) while simultaneously overestimating minimum potentials (MD = 0.53). This bias was particularly noticeable in the predawn potentials of *F. sylvatica* and *Q. petraea* (MD = –0.8 and –1.5, respectively), though the strong correlations for these species (*r* = 0.93 and 0.99, respectively; Table S9) highlight the model’s ability to reproduce relative trends in tree stress, if not absolute values.

**Fig. 29.**
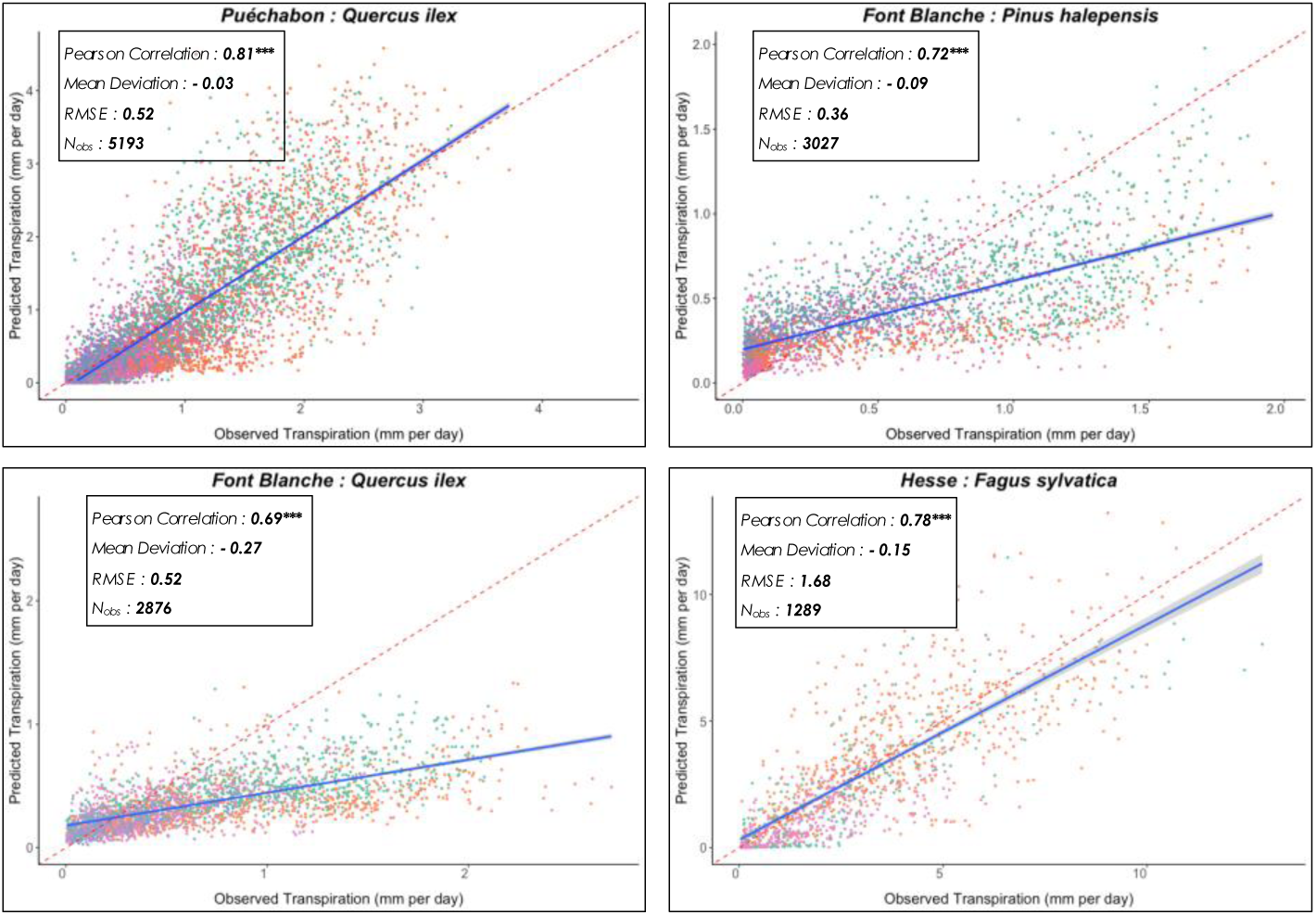
Predicted versus observed species aggregate daily transpirations. For each simulation site, the plain blue line is the regression line of the linear model of the relationship between observed and predicted species aggregate daily transpiration (mm), with confidence interval represented with the grey dashed lines; the dashed red line is the 1:1 line. See Annex X for definition of associated statistics. Colour code for the seasons as follows: 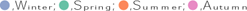

**Fig. 30.**
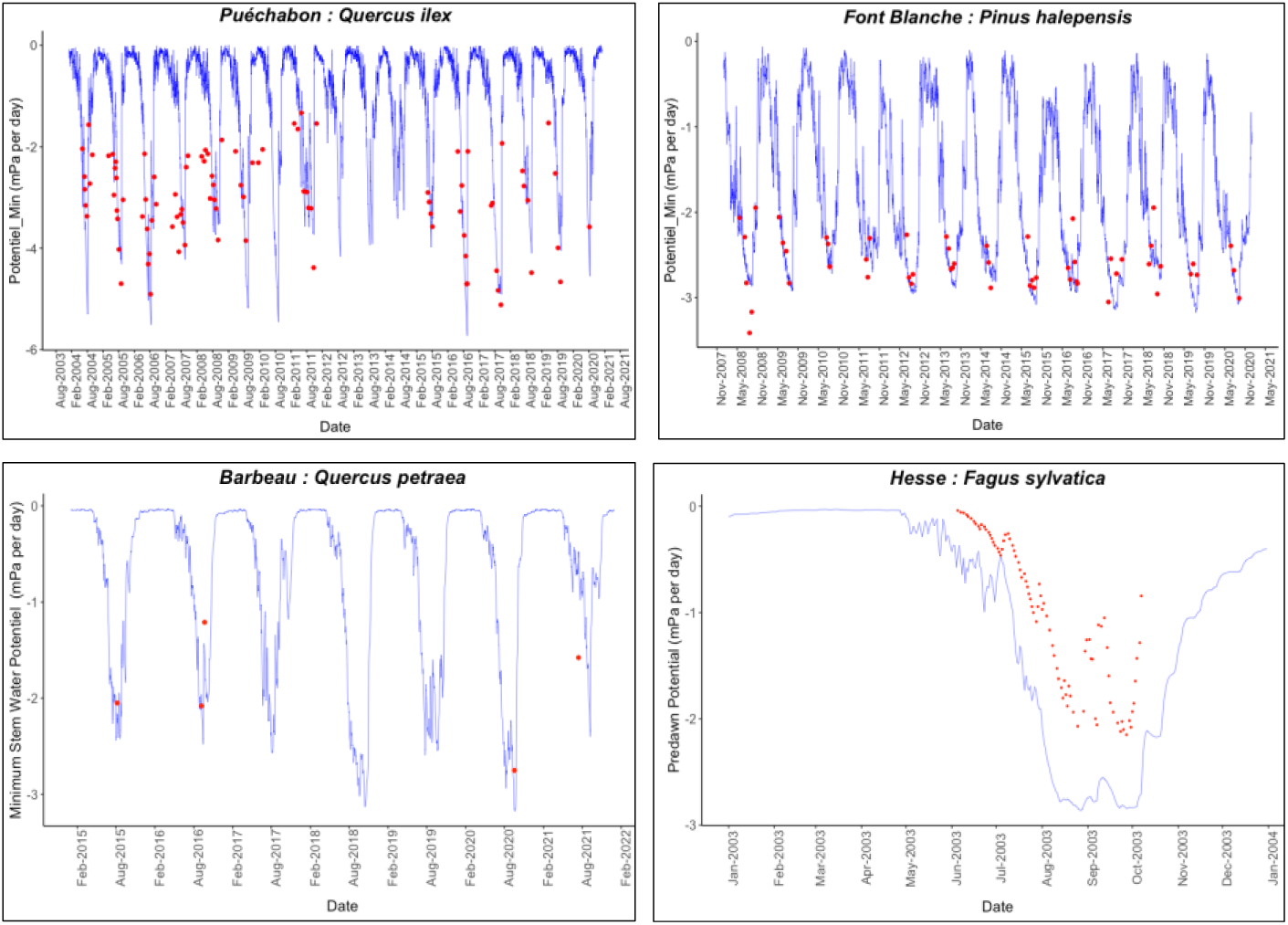
Evolution of predicted versus observed stem water potentials. For the dominant species of the four ICOS simulations, the blue line depicts the daily evolution of the stem water potentials (mPa) generated by the PHOREAU model and averaged over the aggregate trees of the species (refer Annex X for details on the aggregation method). The red points represent the observed water potentials, limited to the years for which observational data is available (data sources are detailed in Annex X). For Puechabon, Font Blanche and Barbeau sites, the minimum daily observed and predicted water potentials are shown. For Hesse, where only predawn observations are available, the maximum predicted water potential is used as a proxy.

**Fig. 31.**
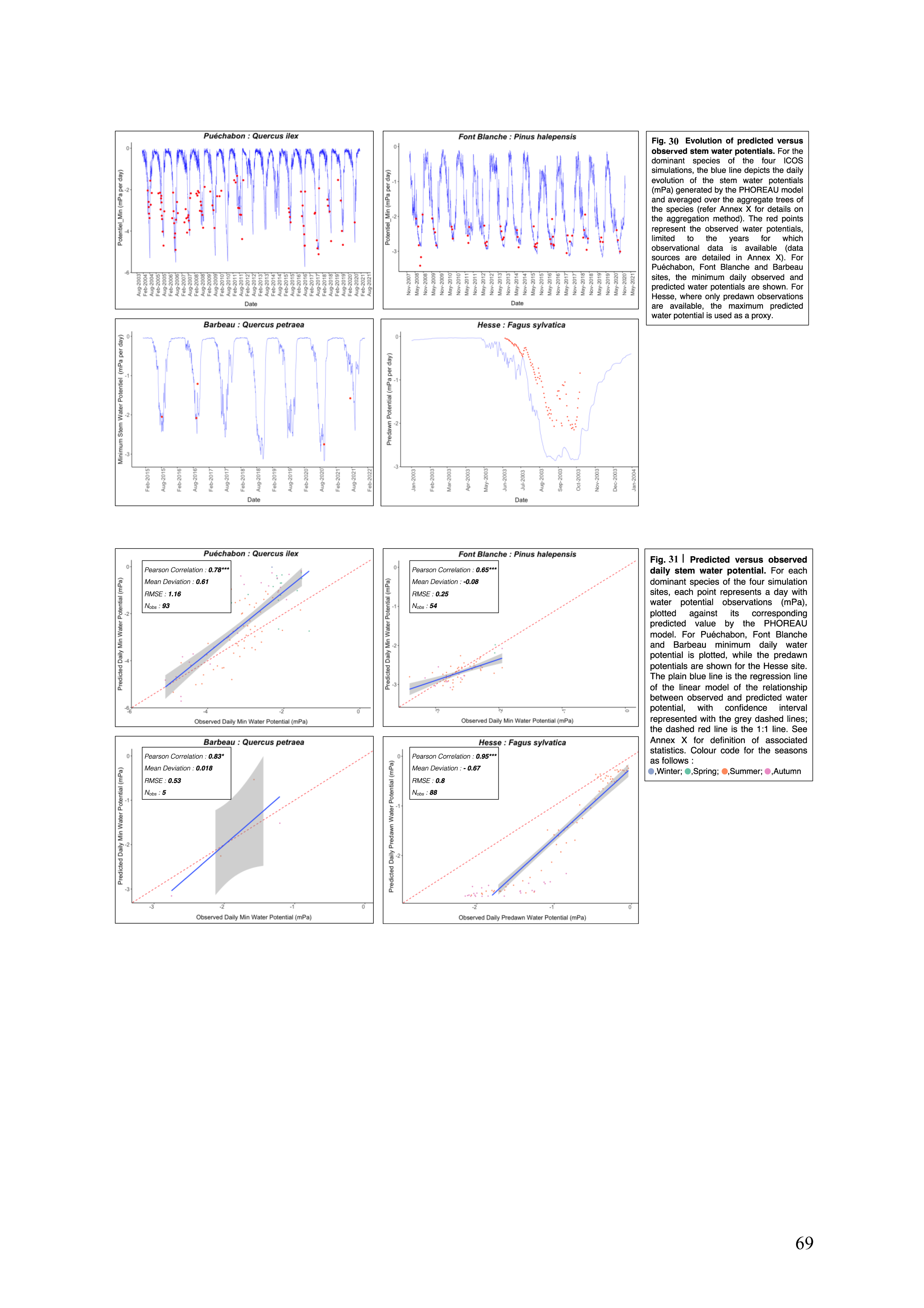
Predicted versus observed daily stem water potential. For each dominant species of the four simulation sites, each point represents a day with water potential observations (mPa), plotted against its corresponding predicted value by the PHOREAU model. For Puechabon, Font Blanche and Barbeau minimum daily water potential is plotted, while the predawn potentials are shown for the Hesse site. The plain blue line is the regression line of the linear model of the relationship between observed and predicted water potential, with confidence interval represented with the grey dashed lines; the dashed red line is the 1:1 line. See Annex X for definition of associated statistics. Colour code for the seasons as follows: 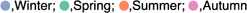

**Fig. 32.**
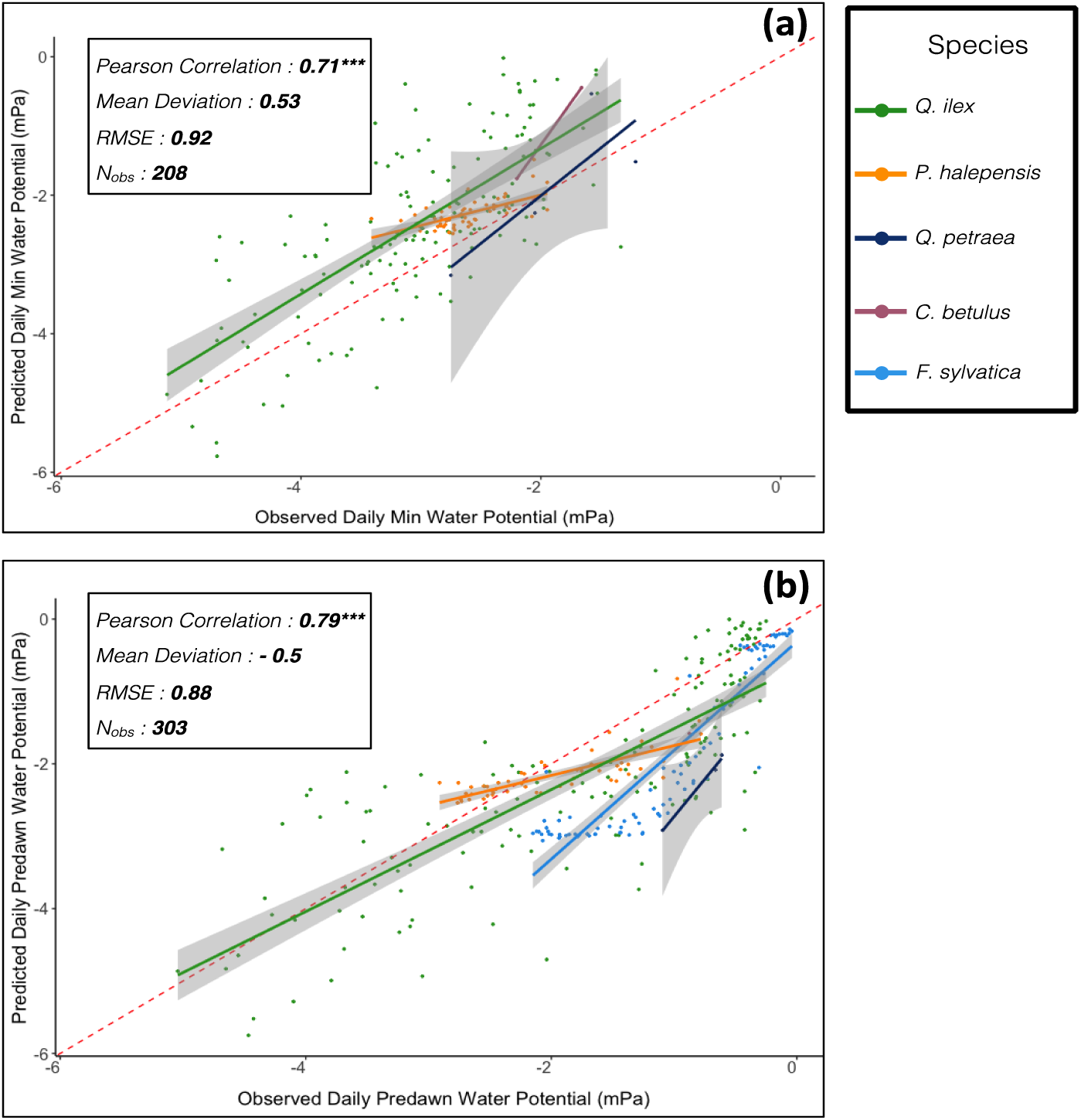
Aggregated Predicted versus observed daily stem water potential. All available stem water potentials (mPa) observations are plotted against the PHOREAU projection for the correspoding day and species. For each species, the full colored lines are the regression lines of the linear model of the relationship between observed and predicted minimum water potential, with confidence interval represented with the grey dashed lines. The dashed red line is the 1:1 line. See Annex X for definition of associated statistics, (a) Comparison with minimum water potentials. (b) Comparison with predawn water potentials.

## 5 Discussion

Depicting and understanding the role of diversity in ecosystem functioning has been a key focus of ecological studies for at least two decades (Kinzig, Pacala and Tilman, 2002; Hooper *et al*., 2012; van der Plas, 2019). In forest ecosystems, the importance of the role of diversity — both structural and compositional — on productivity and wood biomass has been firmly established by numerous studies over a wide range of conditions and methods (Nadrowski, Wirth and Scherer-Lorenzen, 2010; Morin, 2011; Paquette and Messier, 2011; Liang *et al*., 2016; Ratcliffe *et al*., 2017). In addition, there is some evidence that tree diversity could modulate the resistance and recovery of forest productivity under stress (Ammer, 2019; Jourdan, Lebourgeois and Morin, 2019; Schnabel *et al*., 2021; Blondeel *et al*., 2024), although there is no strong consensus on this point, especially regarding possible underlying processes (Decarsin *et al*., 2024). Yet despite these patterns, there remains a scarcity of data regarding the actual differences in functioning of monospecific and mixed forests, and their relative response to changing climate conditions. For these reasons, and because experimenting composition effects in mature forests is especially difficult, the evaluation of diversity effects in forest ecosystems has come to rely increasingly on process-based models (Bohn and Huth, 2017; Maréchaux and Chave, 2017; Jonard *et al*., 2020; Morin *et al*., 2021). The prospective power of these models make them key tools in testing various hypotheses on the diversity-functioning link (Maréchaux *et al*., 2021), but also in evaluating forest management practices that incorporate species mixing (Jourdan *et al*., 2021) and more generally in simulating forest-response to the long-term impacts of climate change.

While the diversity-productivity relationship is well evidenced — a global meta-analysis has shown mixed-species stands were on average 25% more productive than their respective species’ monocultures (Zhang, Chen and Reich, 2012) —, data regarding the link between species diversity and the ability to withstand extreme climatic events is more scarce and contradictory. Where some studies have linked forest diversity to a lessened sensitivity of tree growth to drought (Lebourgeois *et al*., 2013; Anderegg *et al*., 2018; Serrano-León *et al*., 2024) others have found this relationship to be strongly context-dependent (Grossiord *et al*., 2014; Forrester *et al*., 2016; Jactel *et al*., 2017), and restricted to dry environments. Moreover, with the rapid shift in climatic conditions, it would be a mistake to assume that the same patterns of diversity-productivity and diversity-resilience relationships used to support the stress-gradient hypothesis (Bertness and Callaway, 1994) will apply in the next decades to newly drought-prone sites, where water resource limitation has not had the chance to shape the co-evolution of the local species over the past millennia. In fact, the same structural and specific complementarities that are currently responsible for increasing the productivity of existing mixed temperate forests through a better usage of the light resource could become a source of vulnerability, as competition for water intensifies proportionally to the density and foliage areas of the stands (Jucker *et al*., 2014; Haberstroh and Werner, 2022; Decarsin *et al*., 2024).

### 5.1 A process-based model to investigate diversity-productivity and diversity-resilience relationships

The difficulties inherent in integrating trait-based processes in a semi-empiric framework justified validating PHOREAU on a variety of metrics not usually considered for gap models, including predicted foliage area, soil water and stem water potentials. This bottom-up approach mitigated the risk of error compensation, or equifinality, with inversely biased processes negating each other in respect to an integrative metric. Avoiding equifinality was crucial to the development of PHOREAU, because as climatic conditions deviate from the norm in future years, correlations between processes that were equifinal for historical conditions may shift, limiting the ability of the model to accurately predict the impact of climate change on forest functioning. While direct validation on growth is rarely done for gap models because of the inherent difficulty of reproducing such metrics for models not originally designed to work at such short temporal scales (Mette *et al*., 2009; Fyllas *et al*., 2014), the more granular representation of stand functioning of PHOREAU justified our validation on short-term tree and stand productivity. The good performance of the model across the wide range of species and conditions used in our productivity and PNV validation — including Mediterranean and Scandinavian forests — demonstrates its widespread applications to European forest ecosystems. Furthermore, the state-of-the-art validation dataset used in this study will be serve as a baseline to assess any further refinements to the model, as additional species traits become available.

In contrast to fully process-based models than can be parametrized using only functional traits (Davi *et al*., 2005b; Maréchaux and Chave, 2017), PHOREAU eschews a direct representation of carbon assimilation and allocation, in favor of a growth-reduction based approach. While this simplification does distort actual tree functioning and ignores the importance of carbon reserves in buffering year-on-year growth (Körner, 2003), it presents a number of advantages when considering the ecological processes that shape species composition. In addition to a significant gain in computing time, it curtails the uncertainty in model predictions that can result from equifinality, by limiting the number of variables directly impacting growth. Furthermore, by calculating tree growth, leaf area, mortality and establishment rates on the basis of well-established observed optimal values, to which process-based constraints are subsequently applied, we are able to maintain realistic stand basal and foliage areas over the length of the simulation. This model characteristic is a prerequisite to any temporal exploration of diversity-resilience relationships in drought-stressed forests: only by accurately predicting the evolution of forest structure can we decouple its effects from the effects of species-mixing (Forrester and Pretzsch, 2015) for forests functioning at eco-hydrological equilibrium. This is why our integrative validation on the ICOS sites is an important milestone in the development of hydrology-based forest models: unlike usual hydrological validations (Morales *et al*., 2005), not only did PHOREAU provide robust predictions of water fluxes for many years over a diverse set of conditions and species, it did so with no *a priori* fixing of stand leaf and basal area, instead calculating the evolution stand structure on the basis of water-stress feedbacks.

### 5.2 Limitations and future avenues of improvement

Despite good correlations and low average bias, PHOREAU predictions consistently underestimated the observed variability across almost all considered metrics, including soil water quantities, stem water potentials, tree productivity, stand productivity, and stand foliage areas. This attenuating effect is in itself not surprising given the necessary simplifications presented by any modelling approach, and results from a number of unavoidable factors: precision of climatic and soil texture data (especially for ICP II sites); utilization of single sets of species parameters disregarding intra-specific genetic and phenotypic trait variability; lack of 3D representation of competition among trees. While climatic, soil, and species traits inputs can easily be refined for more granular simulations at the local and regional level, taking into account site exposition and fertility, the strong hypothesis of the PHOREAU model regarding the horizontal homogeneity of competition for light and water inside a patch will always be an obstacle to capturing the individual dynamics of trees advantaged or disadvantaged by microtopography and spatial allocation of tree crowns and rooting systems. Despite this inherent limitation, the integration in PHOREAU of many previously disregarded or implicit processes, including explicit roots, phenology, process-based tree hydraulics, and microclimate, has allowed it to outperform the original ForCEEPS model in predicting both short-term growth and long-term species composition.

However, by introducing a more granular representation of tree functioning, PHOREAU has induced a mismatch between some of the parameters used in the model and the role they were originally intended and calibrated for. This mismatch, particularly evident for the optimal species growth rate parameter (*g*_*s*_) and for foliage allometry parameters, is responsible for the difficulty in reproducing the growth of extremely productive trees, and for the large systematic biases observed for certain species like *P. halepensis, P. menziesii, or P. sylvestris.* Because the optimal growth rate in ForCEEPS was calibrated for the main French species based on the top 10^th^ percentile of annual diameter increments measured in the NFI database (IGN, 2020) and for other species dates back to even earlier studies (Didion *et al*., 2009), it is in reality more akin to a growth rate under relatively unconstrained conditions than an actual optimum. As we updated the model’s representation of light and water use constraints to a more process-based approach, we have likely introduced constraints already implicitly present in this aggregated growth rate parameter, essentially penalizing trees twice for the same factor. As we continue to refine the PHOREAU model, a major challenge will therefore be recalibrating this parameter to better reflect actual potential growth unconstrained by competition, despite inherent difficulties in obtaining such data (Pretzsch, 2009).

Similarly, the parameters with which foliage area is derived from tree diameter have not been fully updated to reflect the new importance of foliage area in driving modelled hydraulic fluxes. Despite the many changes introduced in the representation of tree crowns and the partial validation on satellite data, the model demonstrated a poor ability to discriminate between measured litter LAI for sites of similar composition and basal area. Furthermore, neither satellite nor litter-derived total LAI measurements can be used to properly validate the predicted vertical distribution of leaf area. It is this vertical distribution, from which microclimate and individual light-competition constraints are derived, which should next be examined and validated against ground-based LIDAR (Zhao *et al*., 2011, 2015) and microclimate measurements.

Another obvious area of improvement for the model will be a deeper integration of the plant phenology component with other modelled processes. In this study leaf unfolding, leaf senescence, and probability of fruit maturation were computed yearly for an average individual of each species. This method captured inter-specific differences in phenology and temporal light partitioning, but did not account for intra-specific shifts in phenology caused by stand structure. By integrating model variables like microclimate, light availability, and water stress as inputs for an individual-based phenology calculation, PHOREAU will be able to captured well-established variations in leaf phenology between trees of different sociological status (Augspurger and Bartlett, 2003; Cole and Sheldon, 2017; Gressler *et al*., 2015; Schieber, 2012), which are responsible for the persistence of shrubs and saplings in mature forests (Gill, Amthor and Bormann, 1998; Vitasse, 2013).

### 5.3 Establishing baseline available water: retro-engineering PHOREAU to predict rooting depths

One of the main causes for the model’s attenuation of variability in stand and tree productivity was the uncertainty regarding the actual quantity of soil water available to the trees. This uncertainty is itself the result of a twofold gap in information: lack of data for the texture of deeper soil horizons, and the extremely simplified framework used to estimate tree rooting depths. By choosing to reduce the wide observed differences in rooting depths across biomes (Canadell *et al*., 1996; Schenk and Jackson, 2002; Fan *et al*., 2017) and species (Sperry *et al*., 2002; Fan *et al*., 2017) to a simple equation based only on tree size and an aggregate drought index based on past climatic conditions, we intentionally avoided any integration of model results (such as tree foliage area or percentage of embolism) in the calculation of rooting depths, as this would have resulted in an optimization of soil available water on precisely the variables we were trying to validate. Unlike other process-based models validated on stand hydraulic fluxes (Ruffault *et al*., 2023), the fact that PHOREAU produced robust multi-year predictions without using observations to control for stand leaf areas, rooting depths, or actual available water, confirms its possible applications to making dynamic predictions across a large range of forests where this data is not available. Furthermore, now that it has been validated based on *a priori* rooting distributions, the model could be used to predict tree rooting depths for a given forest inventory, climate, and soil data.

Based on the hydrological equilibrium hypothesis (EHE), which states that, in a given edaphic and climatic environment, trade-offs between vegetation water use and drought stress drive canopy density and forest composition toward an optimal hydric state (Eagleson, 1982; Caylor, Scanlon and Rodriguez-Iturbe, 2009), and following the well-substantiated hypothesis that trees function near the point of catastrophic hydraulic failure with narrow safety margins (Tyree and Sperry, 1988; Choat *et al*., 2012), a retro-engineering of PHOREAU could be realized where rooting depths are calculated by optimizing tree available water such that, for a given inventory and soil profile (Kirchen *et al*., 2017), foliage area is maximized (Grier and Running, 1977), and plant minimum water potentials are constrained to values to the point of catastrophic xylem failure. Compared to similar EHE-based statistical (Nemani and Running, 1989) or process-based (Cabon *et al*., 2018) modelling approaches, this retro-engineering of PHOREAU will natively integrate many inter- and intra-specific niche and competition processes integral to actual forest water use. It will furthermore be a necessary first step in establishing a historical baseline when using the model to predict the medium-term impact of global change on forest composition and functioning, as available water is a major determinant in predicting drought-induced die-off events (Allen *et al*., 2010; Anderegg *et al*., 2013; McDowell *et al*., 2013).

### 5.4 Unraveling the effects of trait diversity on competition and coexistence

The novel approach presented in this study, integrating plant functional traits in a forest demographic model, was developed in response to the difficulties encountered by ecologists when verifying hypothesized links between trait diversity and species competition and coexistence. While differences in traits governing resource use should, intuitively, translate into niche differences that maintain coexistence through competition reduction, attempts to directly link trait dispersion with historical species coexistence have proven challenging (McGill *et al*., 2006; Adler *et al*., 2013). This challenge arises from the fact most traits impact competition for several resources at the same time, and that even a temporary advantage in growth can actually result in a lower global fitness when considering population dynamics, with for example feedbacks on drought-induced mortality (Forrester and Pretzsch, 2015) or frost damage due to early onset leaf unfolding (Bigler and Bugmann, 2018). To overcome this difficulty, mechanistic models of resource competition with processes explicitly based on species traits have been proposed as a way to unravel the mechanisms linking trait diversity to forest functioning (Levine *et al*., 2024). Because the effects of climate change on forests will likewise be mediated by complex species mixing effects, the need to develop mechanistic models that bridge the gap between trait-based and ecology and empirical modelling has become urgent to assess the short and medium-terms effects of global warming on existing forests, and discriminate between the possible management scenarios available to forest managers.

The PHOREAU model, having been directly validated for most of its processes, could now be used as an exploratory tool to identify thresholds conditions for species coexistence, dominance, or extinction. A first parsimonious approach could simply consist in identifying, at the edges of a species’ predicted distribution, the main processes — phenology, water-use, or competition for light — limiting its fitness. A more involved exploratory protocol could follow the methodology outlined in Levine *et al*. (2024). By considering predicted species compositions for a wide range of climatic and edaphic conditions, and taking care to distinguish, for each set of condition, the different mechanistic processes which make up a species’ competitive fitness, we could establish relationships between these aggregated model metrics (for example growth reductors) and the underlying species traits. These relationships could then be used to predict the impacts of climate change on forest composition. In parallel to this approach, and as a prerequisite, predicted species compositions should be compared to actual observed compositions, albeit for a much greater set of points than those for the potential composition validation presented in this study, dissipating any remaining uncertainties regarding the representation of regeneration and mortality, which is one of the main current challenges for forest modelling (Cailleret *et al*., 2017; Vanoni *et al*., 2019).

