## Supplementary Tables2 for "PhorEau: a new process-based model to predict forest functioning, from tree ecophysiology to forest dynamics and biogeography"

(a)

| Site | Start | End | Tmp_mean_ONF | Tmp_mean_ERA5 | Tmp_mean_Correct | Tmp_var_ONF | Tmp_var_ERA5 | Tmp_var_Correct | Correlation |
|---|---|---|---|---|---|---|---|---|---|
| **CHP 40** | 2000-01-01 | 2008-07-20 | 12.45 | 13.769 | 14.253 | 40.984 | 40.479 | 38.985 | 0.984 |
| **CHP 59** | 2000-01-01 | 2021-12-31 | 10.1 | 10.485 | 10.551 | 38.066 | 39.42 | 39.086 | 0.984 |
| **CHS 10** | 2000-01-01 | 2008-07-31 | 10.969 | 11.544 | 11.644 | 40.431 | 45.435 | 44.279 | 0.987 |
| **CHS 35** | 2000-01-01 | 2008-07-31 | 11.405 | 11.797 | 12.102 | 31.656 | 31.431 | 30.638 | 0.993 |
| **CHS 41** | 2000-01-01 | 2021-12-31 | 11.597 | 12.203 | 11.827 | 41.378 | 43.013 | 40.184 | 0.985 |
| **DOU 71** | 2000-01-01 | 2008-07-31 | 10.03 | 9.753 | 10.01 | 47.655 | 46.873 | 45.923 | 0.992 |
| **EPC 08** | 2000-01-01 | 2021-12-31 | 9.195 | 9.68 | 8.936 | 43.832 | 42.635 | 41.763 | 0.979 |
| **EPC 63** | 2000-01-01 | 2021-12-31 | 7.919 | 9.604 | 8.676 | 44.336 | 45.579 | 42.463 | 0.977 |
| **EPC 74** | 2000-01-01 | 2008-07-31 | 7.305 | 9.57 | 7.431 | 51.156 | 55.304 | 50.514 | 0.97 |
| **EPC 87** | 2000-01-01 | 2021-12-31 | 9.71 | 10.35 | 9.945 | 45.959 | 43.488 | 41.598 | 0.972 |
| **HET 30** | 2000-01-01 | 2021-12-31 | 7.218 | 9.592 | 7.231 | 46.396 | 43.997 | 42.022 | 0.962 |
| **HET 54a** | 2000-01-01 | 2008-07-31 | 10.054 | 10.329 | 10.587 | 51.412 | 51.212 | 51.045 | 0.994 |
| **HET 64** | 2000-01-01 | 2021-12-31 | 13.315 | 12.353 | 13.059 | 33.658 | 36.109 | 36.596 | 0.967 |
| **PL 20** | 2000-01-01 | 2008-07-02 | 9.711 | 11.13 | 10.065 | 44.577 | 41.677 | 37.14 | 0.983 |
| **PM 40c** | 2000-01-01 | 2008-07-31 | 12.413 | 13.7 | 13.586 | 42.236 | 44.378 | 42.084 | 0.99 |
| **PM 72** | 2000-01-04 | 2008-07-31 | 11.39 | 12.112 | 11.855 | 38.658 | 38.995 | 38.871 | 0.993 |
| **PM 85** | 2000-01-01 | 2008-07-31 | 12.683 | 13.036 | 13.24 | 29.981 | 28.189 | 27.406 | 0.989 |
| **PS 44** | 2000-01-01 | 2008-07-31 | 11.887 | 12.309 | 12.588 | 32.031 | 31.611 | 30.908 | 0.993 |
| **PS 67a** | 2000-01-26 | 2021-12-31 | 10.251 | 10.857 | 11.181 | 51.069 | 53.233 | 51.64 | 0.979 |
| **PS 76** | 2000-01-01 | 2008-07-31 | 10.537 | 11.14 | 11.268 | 32.246 | 31.679 | 32.015 | 0.981 |
| **SP 05** | 2000-01-01 | 2021-12-31 | 8.437 | 5.177 | 8.289 | 52.95 | 63.715 | 51.032 | 0.975 |
| **SP 11** | 2000-01-01 | 2021-12-31 | 9.523 | 11.129 | 10.041 | 41.849 | 40.703 | 40.25 | 0.964 |
| **SP 25** | 2000-01-01 | 2008-07-28 | 8.264 | 7.555 | 8.082 | 50.712 | 48.802 | 50.692 | 0.982 |
| **SP 38** | 2000-01-02 | 2021-12-31 | 8.901 | 7.116 | 9.411 | 53.837 | 57.132 | 56.501 | 0.963 |
| **SP 57** | 2000-01-01 | 2021-12-31 | 9.335 | 9.634 | 10.173 | 47.562 | 51.44 | 50.901 | 0.98 |
| **SP 68** | 2000-01-01 | 2021-12-31 | 8.797 | 8.981 | 8.764 | 49.635 | 52.509 | 52.653 | 0.973 |
| **cumul** | 2000-01-01 | 2021-12-31 | 10.131 | 10.573 | 10.569 | 43.241 | 44.194 | 42.584 | 0.98 |

(b)

| Site | Start | End | Tmn_mean_ONF | Tmn_mean_ERA5 | Tmn_mean_Correct | Tmn_var_ONF | Tmn_var_ERA5 | Tmn_var_Correct | Correlation |
|---|---|---|---|---|---|---|---|---|---|
| **CHP 40** | 2000-01-01 | 2008-07-20 | 7.042 | 9.398 | 9.81 | 40.297 | 35.937 | 34.14 | 0.941 |
| **CHP 59** | 2000-01-01 | 2021-12-31 | 6.105 | 6.951 | 7.004 | 32.615 | 33.035 | 32.62 | 0.948 |
| **CHS 10** | 2000-01-01 | 2008-07-31 | 6.651 | 7.79 | 7.882 | 35.031 | 37.79 | 36.751 | 0.962 |
| **CHS 35** | 2000-01-01 | 2008-07-31 | 7.49 | 8.363 | 8.648 | 27.253 | 26.74 | 25.905 | 0.965 |
| **CHS 41** | 2000-01-01 | 2021-12-31 | 7.232 | 8.309 | 7.998 | 33.511 | 34.688 | 33.019 | 0.961 |
| **DOU 71** | 2000-01-01 | 2008-07-31 | 6.149 | 6.236 | 6.467 | 37.434 | 37.731 | 36.864 | 0.975 |
| **EPC 08** | 2000-01-01 | 2021-12-31 | 6.099 | 6.225 | 5.551 | 34.934 | 35.758 | 35.844 | 0.962 |
| **EPC 63** | 2000-01-01 | 2021-12-31 | 3.118 | 5.553 | 4.818 | 36.46 | 36.534 | 35.551 | 0.932 |
| **EPC 74** | 2000-01-01 | 2008-07-31 | 4.368 | 4.807 | 3.217 | 44.513 | 47.406 | 47.552 | 0.915 |
| **EPC 87** | 2000-01-01 | 2021-12-31 | 6.168 | 6.853 | 6.513 | 35.449 | 34.676 | 33.766 | 0.954 |
| **HET 30** | 2000-01-01 | 2021-12-31 | 3.933 | 5.488 | 3.517 | 40.43 | 34.678 | 36.256 | 0.929 |
| **HET 54a** | 2000-01-01 | 2008-07-31 | 5.801 | 6.518 | 6.733 | 40.921 | 42.384 | 41.9 | 0.968 |
| **HET 64** | 2000-01-01 | 2021-12-31 | 9.273 | 8.201 | 8.726 | 28.973 | 30.585 | 30.354 | 0.953 |
| **PL 20** | 2000-01-01 | 2008-07-02 | 5.4 | 7.886 | 6.939 | 33.121 | 37.095 | 33.625 | 0.961 |
| **PM 40c** | 2000-01-01 | 2008-07-31 | 7.22 | 9.344 | 9.273 | 36.231 | 35.35 | 33.987 | 0.947 |
| **PM 72** | 2000-01-04 | 2008-07-31 | 7.54 | 8.418 | 8.182 | 31.514 | 32.465 | 32.55 | 0.975 |
| **PM 85** | 2000-01-01 | 2008-07-31 | 9.08 | 10.412 | 10.611 | 30.195 | 26.114 | 25.144 | 0.957 |
| **PS 44** | 2000-01-01 | 2008-07-31 | 7.687 | 8.767 | 9.033 | 27.511 | 27.657 | 26.744 | 0.966 |
| **PS 67a** | 2000-01-26 | 2021-12-31 | 5.203 | 7.11 | 7.381 | 40.434 | 43.407 | 41.885 | 0.923 |
| **PS 76** | 2000-01-01 | 2008-07-31 | 5.475 | 7.77 | 7.876 | 32.665 | 27.414 | 27.467 | 0.888 |
| **SP 05** | 2000-01-01 | 2021-12-31 | 3.891 | 0.176 | 3.173 | 39.154 | 57.487 | 41.531 | 0.931 |
| **SP 11** | 2000-01-01 | 2021-12-31 | 5.461 | 7.059 | 6.186 | 33.02 | 35.258 | 36.308 | 0.951 |
| **SP 25** | 2000-01-01 | 2008-07-28 | 4.508 | 3.965 | 4.402 | 42.506 | 43.115 | 43.847 | 0.969 |
| **SP 38** | 2000-01-02 | 2021-12-31 | 5.658 | 1.987 | 3.789 | 42.986 | 52.942 | 46.482 | 0.908 |
| **SP 57** | 2000-01-01 | 2021-12-31 | 5.06 | 5.915 | 6.368 | 36.223 | 42.776 | 41.644 | 0.949 |
| **SP 68** | 2000-01-01 | 2021-12-31 | 5.894 | 5.018 | 4.81 | 41.284 | 44.03 | 44.527 | 0.953 |
| **Cumul** | 2000-01-01 | 2021-12-31 | 6.058 | 6.712 | 6.727 | 35.949 | 37.425 | 36.01 | 0.948 |

(c)

| Site | startDay | endDay | Tmx_mean_ONF | Tmx_mean_ERA5 | Tmx_mean_Correct | Tmx_var_ONF | Tmx_var_ERA5 | Tmx_var_Correct | Correlation |
|---|---|---|---|---|---|---|---|---|---|
| **CHP 40** | 2000-01-01 | 2008-07-20 | 19.526 | 18.591 | 19.157 | 54.923 | 51.533 | 50.604 | 0.978 |
| **CHP 59** | 2000-01-01 | 2021-12-31 | 14.878 | 14.002 | 14.084 | 57.51 | 48.585 | 48.303 | 0.978 |
| **CHS 10** | 2000-01-01 | 2008-07-31 | 16.414 | 15.326 | 15.432 | 59.417 | 55.905 | 54.642 | 0.98 |
| **CHS 35** | 2000-01-01 | 2008-07-31 | 16.45 | 15.447 | 15.768 | 45.331 | 39.707 | 39.163 | 0.983 |
| **CHS 41** | 2000-01-01 | 2021-12-31 | 16.911 | 16.092 | 15.663 | 61.048 | 54.337 | 50.175 | 0.976 |
| **DOU 71** | 2000-01-01 | 2008-07-31 | 14.736 | 13.407 | 13.683 | 65.709 | 58.58 | 57.649 | 0.985 |
| **EPC 08** | 2000-01-01 | 2021-12-31 | 13.068 | 13.057 | 12.262 | 60.454 | 51.869 | 49.841 | 0.974 |
| **EPC 63** | 2000-01-01 | 2021-12-31 | 13.085 | 13.782 | 12.679 | 63.688 | 58.828 | 53.546 | 0.972 |
| **EPC 74** | 2000-01-01 | 2008-07-31 | 11.369 | 13.916 | 11.454 | 64.775 | 61.349 | 54.256 | 0.976 |
| **EPC 87** | 2000-01-01 | 2021-12-31 | 14.577 | 14.056 | 13.591 | 66.483 | 55.248 | 52.234 | 0.969 |
| **HET 30** | 2000-01-01 | 2021-12-31 | 11.572 | 13.949 | 11.318 | 61.554 | 55.733 | 49.547 | 0.958 |
| **HET 54a** | 2000-01-01 | 2008-07-31 | 15.162 | 14.094 | 14.388 | 75.808 | 62.786 | 62.852 | 0.987 |
| **HET 64** | 2000-01-01 | 2021-12-31 | 18.526 | 16.805 | 17.7 | 47.798 | 46.729 | 47.775 | 0.958 |
| **PL 20** | 2000-01-01 | 2008-07-02 | 15.349 | 14.308 | 13.14 | 60.686 | 46.59 | 41.467 | 0.979 |
| **PM 40c** | 2000-01-01 | 2008-07-31 | 19.283 | 18.674 | 18.518 | 60.79 | 58.19 | 54.835 | 0.981 |
| **PM 72** | 2000-01-04 | 2008-07-31 | 16.069 | 15.846 | 15.57 | 54.332 | 48.269 | 47.772 | 0.987 |
| **PM 85** | 2000-01-01 | 2008-07-31 | 16.889 | 15.69 | 15.902 | 39.004 | 33.42 | 32.949 | 0.977 |
| **PS 44** | 2000-01-01 | 2008-07-31 | 17.111 | 15.947 | 16.24 | 46.031 | 39.387 | 39.087 | 0.983 |
| **PS 67a** | 2000-01-26 | 2021-12-31 | 16.088 | 14.67 | 15.037 | 78.91 | 66.331 | 64.66 | 0.975 |
| **PS 76** | 2000-01-01 | 2008-07-31 | 16.005 | 14.537 | 14.686 | 46.389 | 39.176 | 39.858 | 0.985 |
| **SP 05** | 2000-01-01 | 2021-12-31 | 14.491 | 9.649 | 12.845 | 70.04 | 71.387 | 61.682 | 0.968 |
| **SP 11** | 2000-01-01 | 2021-12-31 | 14.381 | 15.47 | 14.172 | 56.822 | 48.759 | 47.158 | 0.954 |
| **SP 25** | 2000-01-01 | 2008-07-28 | 12.72 | 11.339 | 11.95 | 65.522 | 57.086 | 60.05 | 0.98 |
| **SP 38** | 2000-01-02 | 2021-12-31 | 13.211 | 11.619 | 14.206 | 71.663 | 63.94 | 67.943 | 0.964 |
| **SP 57** | 2000-01-01 | 2021-12-31 | 14.765 | 13.314 | 13.922 | 75.435 | 63.081 | 63.048 | 0.976 |
| **SP 68** | 2000-01-01 | 2021-12-31 | 12.614 | 12.935 | 12.71 | 66.597 | 63.263 | 62.857 | 0.973 |
| **cumul** | 2000-01-01 | 2021-12-31 | 15.202 | 14.482 | 14.465 | 60.643 | 53.849 | 52.075 | 0.975 |

(d)

| Site | Start | End | Pre_mean_ONF | Pre_mean_ERA5 | Pre_mean_Correct | Pre_var_ONF | Pre_var_ERA5 | Pre_var_Correct | Correlation |
|---|---|---|---|---|---|---|---|---|---|
| **CHP 40** | 2000-01-01 | 2008-07-20 | 0.278 | 0.264 | 0.28 | 0.395 | 0.232 | 0.286 | 0.72 |
| **CHP 59** | 2000-01-01 | 2021-12-31 | 0.227 | 0.246 | 0.272 | 0.2 | 0.157 | 0.2 | 0.666 |
| **CHS 10** | 2000-01-01 | 2008-07-31 | 0.237 | 0.25 | 0.22 | 0.246 | 0.165 | 0.138 | 0.698 |
| **CHS 35** | 2000-01-01 | 2008-07-31 | 0.225 | 0.226 | 0.23 | 0.201 | 0.165 | 0.175 | 0.676 |
| **CHS 41** | 2000-01-01 | 2021-12-31 | 0.197 | 0.202 | 0.204 | 0.195 | 0.14 | 0.152 | 0.644 |
| **DOU 71** | 2000-01-01 | 2008-07-31 | 0.385 | 0.309 | 0.402 | 0.57 | 0.246 | 0.44 | 0.751 |
| **EPC 08** | 2000-01-01 | 2021-12-31 | 0.335 | 0.291 | 0.342 | 0.414 | 0.201 | 0.301 | 0.683 |
| **EPC 63** | 2000-01-01 | 2021-12-31 | 0.272 | 0.264 | 0.328 | 0.326 | 0.223 | 0.332 | 0.624 |
| **EPC 74** | 2000-01-01 | 2008-07-31 | 0.346 | 0.403 | 0.38 | 0.523 | 0.446 | 0.432 | 0.66 |
| **EPC 87** | 2000-01-01 | 2021-12-31 | 0.375 | 0.351 | 0.356 | 0.527 | 0.324 | 0.345 | 0.706 |
| **HET 30** | 2000-01-01 | 2021-12-31 | 0.616 | 0.3 | 0.472 | 3.73 | 0.582 | 1.626 | 0.723 |
| **HET 54a** | 2000-01-01 | 2008-07-31 | 0.24 | 0.285 | 0.276 | 0.25 | 0.233 | 0.238 | 0.759 |
| **HET 64** | 2000-01-01 | 2021-12-31 | 0.374 | 0.407 | 0.346 | 0.624 | 0.501 | 0.374 | 0.722 |
| **PL 20** | 2000-01-01 | 2008-07-02 | 0.314 | 0.236 | 0.34 | 0.825 | 0.3 | 0.692 | 0.668 |
| **PM 40c** | 2000-01-01 | 2008-07-31 | 0.229 | 0.21 | 0.225 | 0.252 | 0.165 | 0.197 | 0.699 |
| **PM 72** | 2000-01-04 | 2008-07-31 | 0.223 | 0.22 | 0.218 | 0.214 | 0.163 | 0.166 | 0.724 |
| **PM 85** | 2000-01-01 | 2008-07-31 | 0.198 | 0.204 | 0.209 | 0.179 | 0.136 | 0.151 | 0.73 |
| **PS 44** | 2000-01-01 | 2008-07-31 | 0.238 | 0.211 | 0.232 | 0.253 | 0.152 | 0.195 | 0.737 |
| **PS 67a** | 2000-01-26 | 2021-12-31 | 0.195 | 0.251 | 0.214 | 0.177 | 0.17 | 0.13 | 0.629 |
| **PS 76** | 2000-01-01 | 2008-07-31 | 0.238 | 0.248 | 0.204 | 0.203 | 0.143 | 0.103 | 0.703 |
| **SP 05** | 2000-01-01 | 2021-12-31 | 0.249 | 0.262 | 0.23 | 0.501 | 0.314 | 0.262 | 0.68 |
| **SP 11** | 2000-01-01 | 2021-12-31 | 0.309 | 0.316 | 0.227 | 0.507 | 0.4 | 0.259 | 0.576 |
| **SP 25** | 2000-01-01 | 2008-07-28 | 0.389 | 0.485 | 0.445 | 0.571 | 0.506 | 0.458 | 0.741 |
| **SP 38** | 2000-01-02 | 2021-12-31 | 0.384 | 0.488 | 0.372 | 0.706 | 0.619 | 0.403 | 0.686 |
| **SP 57** | 2000-01-01 | 2021-12-31 | 0.324 | 0.321 | 0.313 | 0.386 | 0.25 | 0.256 | 0.696 |
| **SP 68** | 2000-01-01 | 2021-12-31 | 0.331 | 0.336 | 0.435 | 0.477 | 0.289 | 0.537 | 0.682 |
| **cumul** | 2000-01-01 | 2021-12-31 | 0.297 | 0.292 | 0.299 | 0.517 | 0.278 | 0.34 | 0.692 |

**Table** [redacted] Complete results of comparisons between on-site ONF measurements and reconstructed climates for the 26 sites where measurements were available. For each site climate data were aggregated between the start and end dates for on-site measurement. Mean and standard deviation calculated for observed, reconstructed, and raw ERA-5 Land time series. Accuracy of reconstructed variables against observed variable measured through Pearson correlation. Significance of the Pearson correlation coefficient (*** : $p < 0.001$; ** : $p < 0.01$;  * : $p < 0.05$; ns : $p > 0.05$) . (a) for mean daily temperatures (°C); (b) for minimum daily temperatures (°C); (c) for maximum daily temperatures (°C) ; (d) for daily precipitation sum (mm)
