## Supplementary Figures for "PhorEau: a new process-based model to predict forest functioning, from tree ecophysiology to forest dynamics and biogeography"

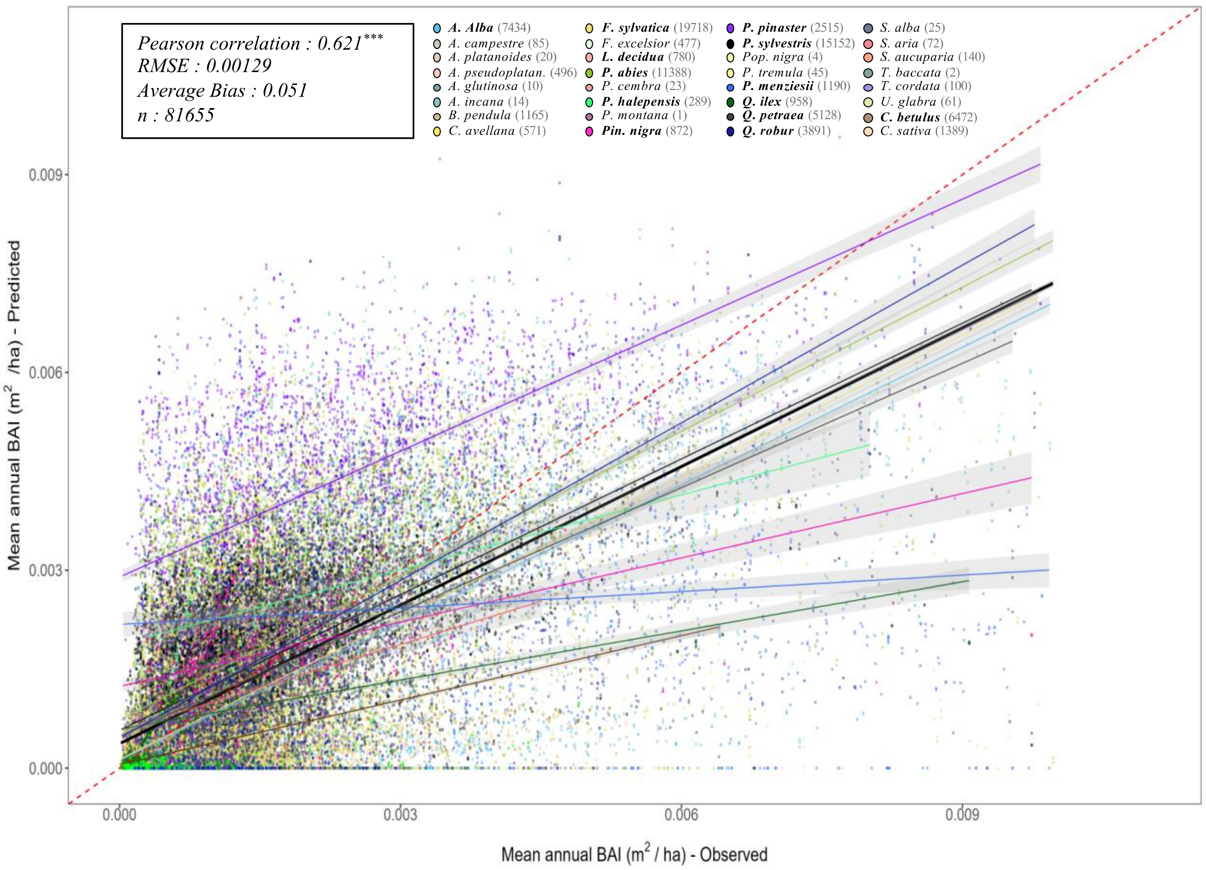


**Fig. S1 ǀ Projected (by ForCEEPS) against observed mean annual tree basal increments (BAI)** for all simulated trees over the 340 RENECOFOR and ICP II validation inventories. Tree points are color coded by species (see legend above). The dashed red-line is the 1:1 line; other full lines represent the regression lines of the linear model between observed and predicted tree productivity, with confidence intervals represented by the grey shaded area (in black the overall regression; coloured lines for species-specific regressions). Species-specific regressions are only shown for stand dominant species (in bold in legend). Associated statistics for the global simulation in top left, while species-specific statistics can be found in Table S1.

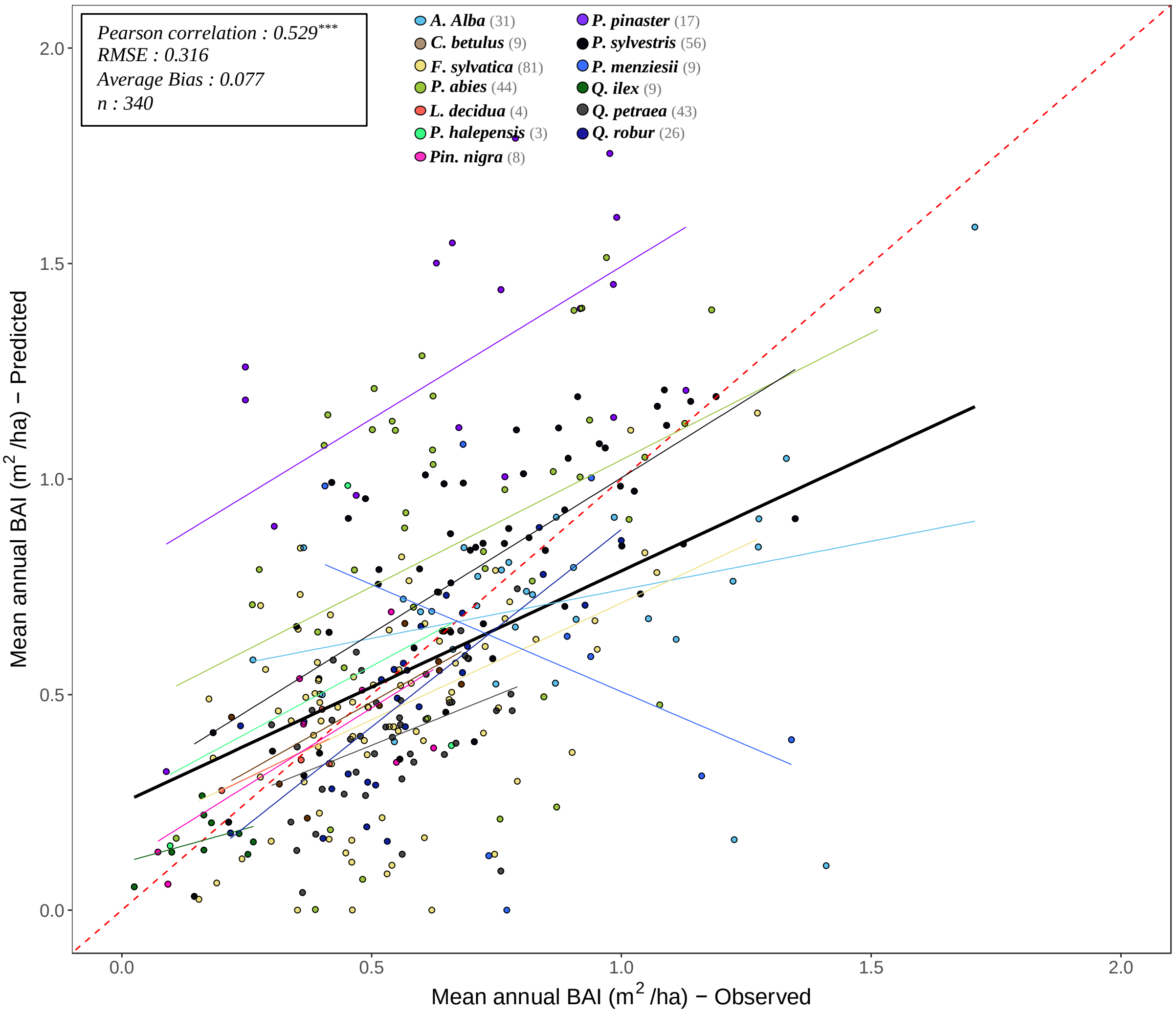


**Fig. S2 | Projected (by ForCEEPS) against observed mean annual stand basal increments (BAI)** for all 340 RENECOFOR and ICP II validation inventories. Stand points are color coded by dominant species (see legend above). The dashed red-line is the 1:1 line; other full lines represent the regression lines of the linear model between observed and predicted stand productivity, with confidence intervals represented by the grey shaded area (in black the overall regression; coloured lines for species-specific regressions). Associated statistics for the global simulation in top left, while species-specific statistics can be found in Table S2.

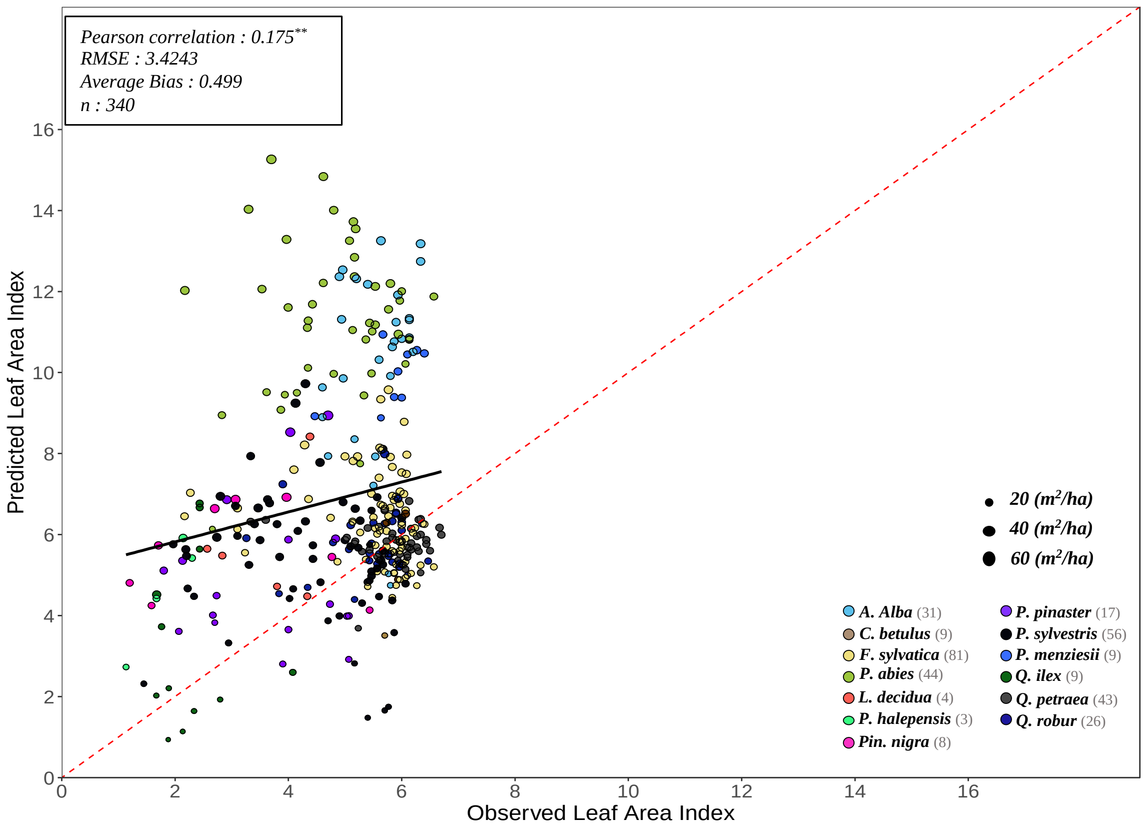


**Fig. S3 │ Projected (by ForCEEPS) against observed satellite leaf area index (LAI)** for all 340 RENECOFOR and ICP II validation inventories. The y-axis shows the LAI predicted by the model from the stand inventory at the start of the simulation, while the x-axis represents the PROBA-V LAI value for the maching coordinate and inventory year, averaged between July, August and September. Stand points are color coded by dominant species (see legend in bottom left). The size of points shows inventory basal area. The dashed red-line is the 1:1 line; the black full line represent the regression line of the linear model between observed and predicted LAI, with confidence interval represented by the grey shaded area. Associated statistics in top left.

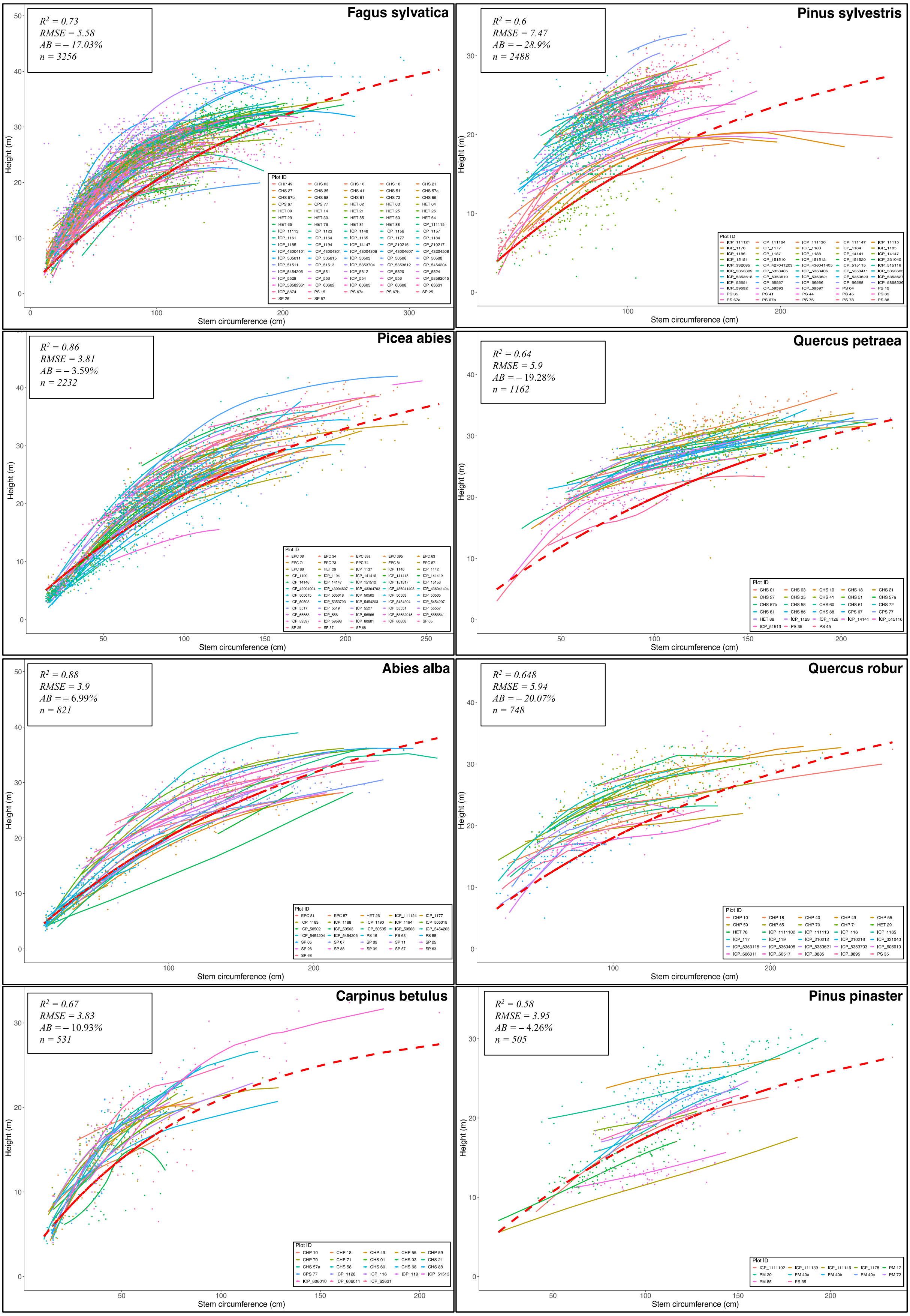
Fagus sylvatica
$R^2 = 0.73$
RMSE = 5.58
AB = – 17.03%
n = 3256
Pinus sylvestris
$R^2 = 0.6$
RMSE = 7.47
AB = – 28.9%
n = 2488
Picea abies
$R^2 = 0.86$
RMSE = 3.81
AB = – 3.59%
n = 2232
Quercus petraea
$R^2 = 0.64$
RMSE = 5.9
AB = – 19.28%
n = 1162
Abies alba
$R^2 = 0.88$
RMSE = 3.9
AB = – 6.99%
n = 821
Quercus robur
$R^2 = 0.648$
RMSE = 5.94
AB = – 20.07%
n = 748
Carpinus betulus
$R^2 = 0.67$
RMSE = 3.83
AB = – 10.93%
n = 531
Pinus pinaster
$R^2 = 0.58$
RMSE = 3.95
AB = – 4.26%
n = 505
Height (m)
Stem circumference (cm)
Plot ID

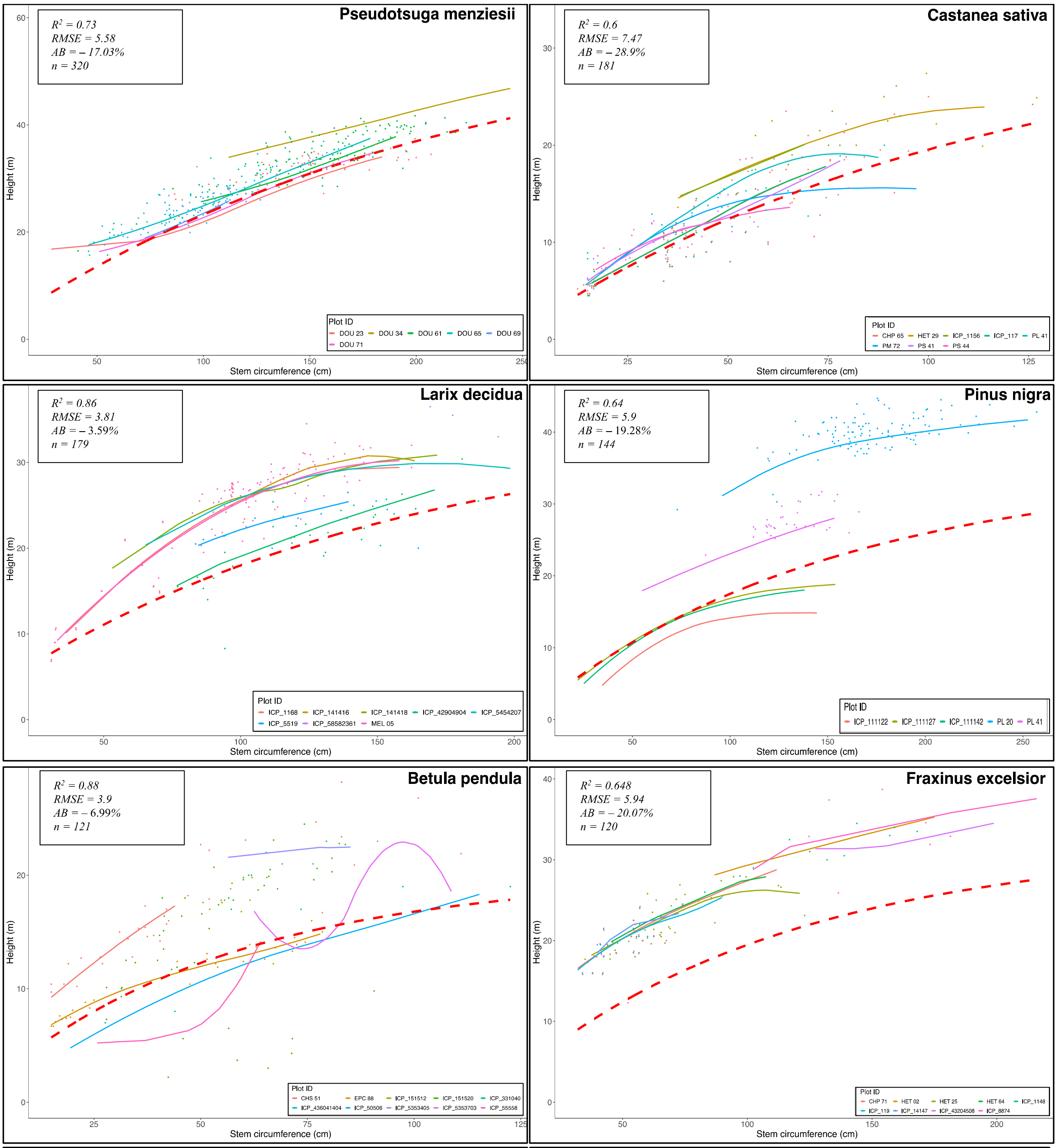


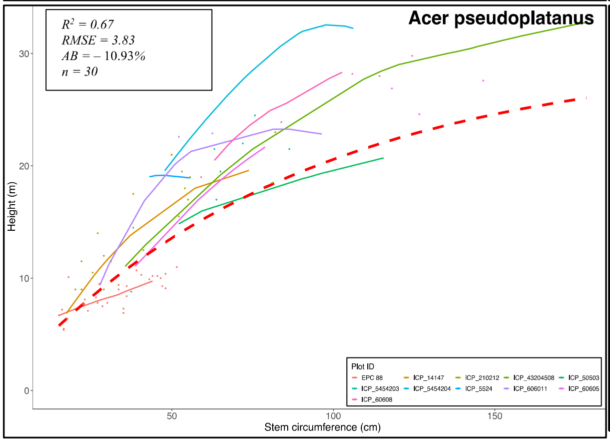


**Fig. S4 | Diversity of height-to-diameter ratios among study sites.** For each species with sufficient data, for each of the 245 RENECOFOR and ICP II study sites, points show measured height against circumference of individual trees present in PHOREAU simulation inventories. Full lines, colored-coded by site, represent local-regression curves used to calculate height for trees with no height measurement. Height-to-diameter curves were interpolated for each plot using a LOESS method n tree measurements (not limited to simulation periods). The dashed red line represents the ForCEEPS species height-to-diameter formula. Predicted tree heights from the formula were evaluated against measured tree heights to produce associated statistics, detailed in Table X.

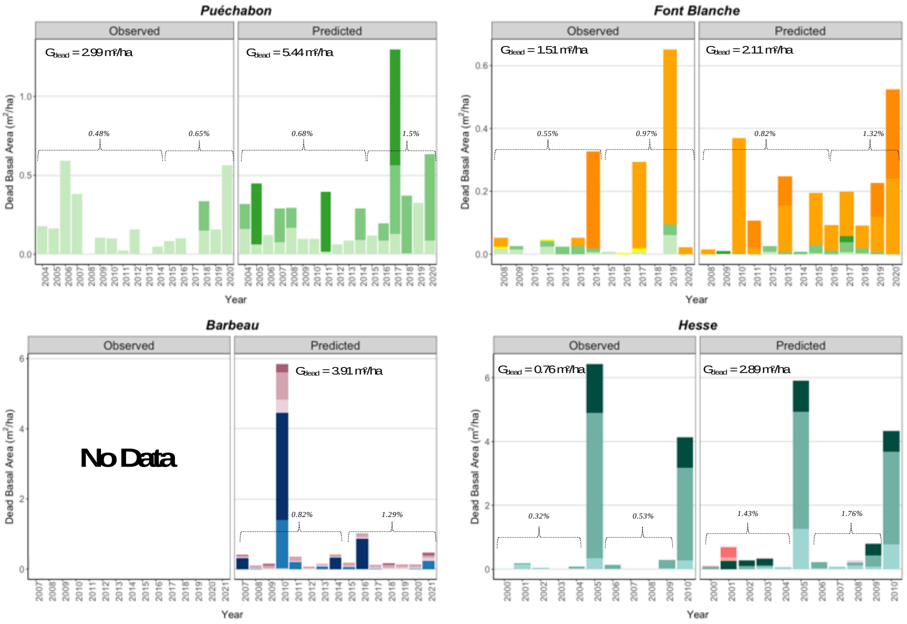


**Fig. S5 │ Predicted versus observed evolution annual stand basal area loss due to mortality.** For each simulation site, the bars depict the summed annual total basal area (m$^2$/ha) of all dead trees, broken down by species and size class (refer to Annex X for details). Observed values are derived from stand inventories, while predicted values are generated by the PHOREAU model. Also shown are the yearly basal area loss rates, calculated relative to the initial basal area for two distinct time periods in each simulation, along with the total basal area dieback per hectare ($G_{dead}$). Transparent bars indicate years with thinnings (see Annex X for details), which are excluded from the the mortality statistics.
